# In utero barcoding of mouse mesoderm reveals the clonal architecture of organ mesenchyme

**DOI:** 10.64898/2026.09.22.752134

**Authors:** Jingyan He, Johan Lorentz, Agustín A. Corbat, Sergey Isaev, Raphaël Mauron, Sandra De Haan, Lenka Belicová, Jia Sun, Bettina Semsch, Noémi Van Hul, Robert Tonoian, John Wilson, Ulrika Marklund, Marcel Martin, Neil C. Henderson, Jonas Frisén, Michael Ratz, Peter V. Kharchenko, Igor Adameyko, Enikő Lázár, Joakim Lundeberg, Emma R Andersson

## Abstract

The mesoderm coordinates organogenesis and generates the cardiovascular, visceral and musculoskeletal systems, yet in mammals this germ layer is difficult to access, impeding its genetic manipulation and study. Here, we establish embryonic day (E) 7.5 exocoelomic cavity nano-injection as a scalable route for selective genetic targeting of mouse mesoderm. Combining in utero lentiviral barcoding with single-cell and spatial transcriptomics, we reconstructed mesodermal and neural crest clonal relationships across >590,000 cells and 31,000 multicellular clones from E9.5–E10.5 embryos and E16.5 livers and hearts. In liver, capsular mesothelium was closely related to hepatic stellate cells, whereas fibroblast and vascular smooth muscle lineages diverged earlier. In heart, epicardium was clonally coupled to three spatially restricted mesenchymal sub-lineages but not to cardiomyocytes, and lineage tracing separated epicardial- and neural crest-derived populations within transcriptionally convergent valve mesenchyme. This approach enables Cre-independent manipulation of mammalian mesoderm and resolves organ mesenchyme development at clonal resolution.

**HIGHLIGHTS:** - E7.5 exocoelomic nano-injection barcodes mouse mesoderm without Cre drivers.

- Clonal maps span >590,000 cells and 31,000 multicellular clones in vivo.

- Mesothelial subsets are lineage-biased sources of liver mesenchymal diversity.

- Epicardium seeds three spatial mesenchymal lineages but not cardiomyocytes.

- Clonal tracing resolves convergent neural crest and mesodermal valve mesenchyme.

## INTRODUCTION

Understanding how the mammalian mesoderm generates the diverse cell types of the body is central to developmental biology. Mesoderm gives rise to cardiovascular, musculoskeletal, urogenital, visceral, hematopoietic and connective tissues, and provides inductive signals that coordinate organogenesis^1^. Unlike embryonic epithelia, which can be directly genetically manipulated in utero using amniotic cavity injections ^2,3^, mesodermal progenitors remain difficult to access and manipulate. This has limited systematic reconstruction of mesodermal lineage relationships during mammalian organogenesis.

Current models of mesodermal development rely heavily on genetic lineage tracing using Cre drivers. Although powerful, these approaches are constrained by enhancer specificity, timing, recombination efficiency and the fact that marker expression does not necessarily imply shared clonal ancestry ^4,5^. These limitations are particularly evident in organ mesenchyme, where transient progenitor populations express overlapping markers and generate transcriptionally related derivatives. As a result, the developmental origins of several mesenchymal populations remain unresolved.

The liver and heart provide two prominent examples of organs with complex mesenchymal populations. In the liver, septum transversum-derived mesenchyme and mesothelium have been implicated as sources of hepatic stellate cells, fibroblasts and vascular smooth muscle cells ^6–9^. In the heart, the proepicardium gives rise to the epicardium, which has been proposed to generate fibroblasts, pericytes, vascular smooth muscle cells and other intracardiac populations ^10,11^. However, these conclusions are largely based on marker-driven lineage tracing, and it remains unclear which cell types share recent ancestry, which diverge earlier, and whether transcriptionally similar mesenchymal populations can arise from distinct developmental origins.

Here, we developed a novel method for genetic manipulation of the mouse embryonic mesoderm via ultrasound-guided embryonic day (E) 7.5 nano-injection into the exocoelomic cavity. We show that its continuity with the intraembryonic coelom provides access to mesodermal progenitors and enables their widespread, selective transduction. In utero nano-injection was initially developed to target and manipulate developing mouse skin ^2,12,13^ and has recently been adapted to target ectoderm, paving the way to high-throughput studies of the neural plate, neural crest and placode derivatives^3,14,15^. Using in vivo delivery of lentiviral barcodes and next generation single cell lineage (scLineage) tracing ^4,16^, we generated high-resolution clonal lineage maps of early E9.5-E10.5 embryonic mesoderm and established its contribution to E16.5 heart and liver lineages. This work overcomes longstanding barriers in developmental genetics and delivers both a powerful experimental platform for early mesoderm targeting and new biological insights into the organization of mesodermal lineages, with broad implications for understanding congenital disorders and vertebrate organogenesis.

## RESULTS

### E7.5 exocoelomic cavity nano-injection specifically targets mesoderm

At embryonic day (E) 8 the mouse extra-embryonic exocoelomic cavity is continuous with the intra-embryonic coelomic cavity ^17^, hypothetically providing an indirect access point to target mesoderm via viral injections into the more accessible exocoelomic cavity (**Figure 1A**). To map which germ layers and cell types are targeted by exocoelomic cavity injection, we injected lentivirus encoding either a fluorescent protein alone, or a lentiviral barcode library co-expressing a fluorophore, at E7.5. Immunofluorescence analysis confirmed that exocoelomic cavity injection efficiently targeted mesoderm derivatives including the heart, epicardium surrounding the heart (**Figures 1B, S1A, box #1**), Wilms tumor 1-positive (WT1+) liver mesothelial cells, and Activated Leukocyte Cell Adhesion Molecule-positive (ALCAM/CD166+) septum transversum mesenchyme, and liver mesenchymal cells (**Figures 1B, box #2, 1C, S1A box #2**). However, endocardium was not labeled (**Figure S1B**), corroborating the previously reported early segregation of progenitors committed to an endocardial or myocardial fate ^18–20^, and suggesting that endocardial progenitors are inaccessible to exocoelomic cavity injection at E7.5.

**Figure 1.**
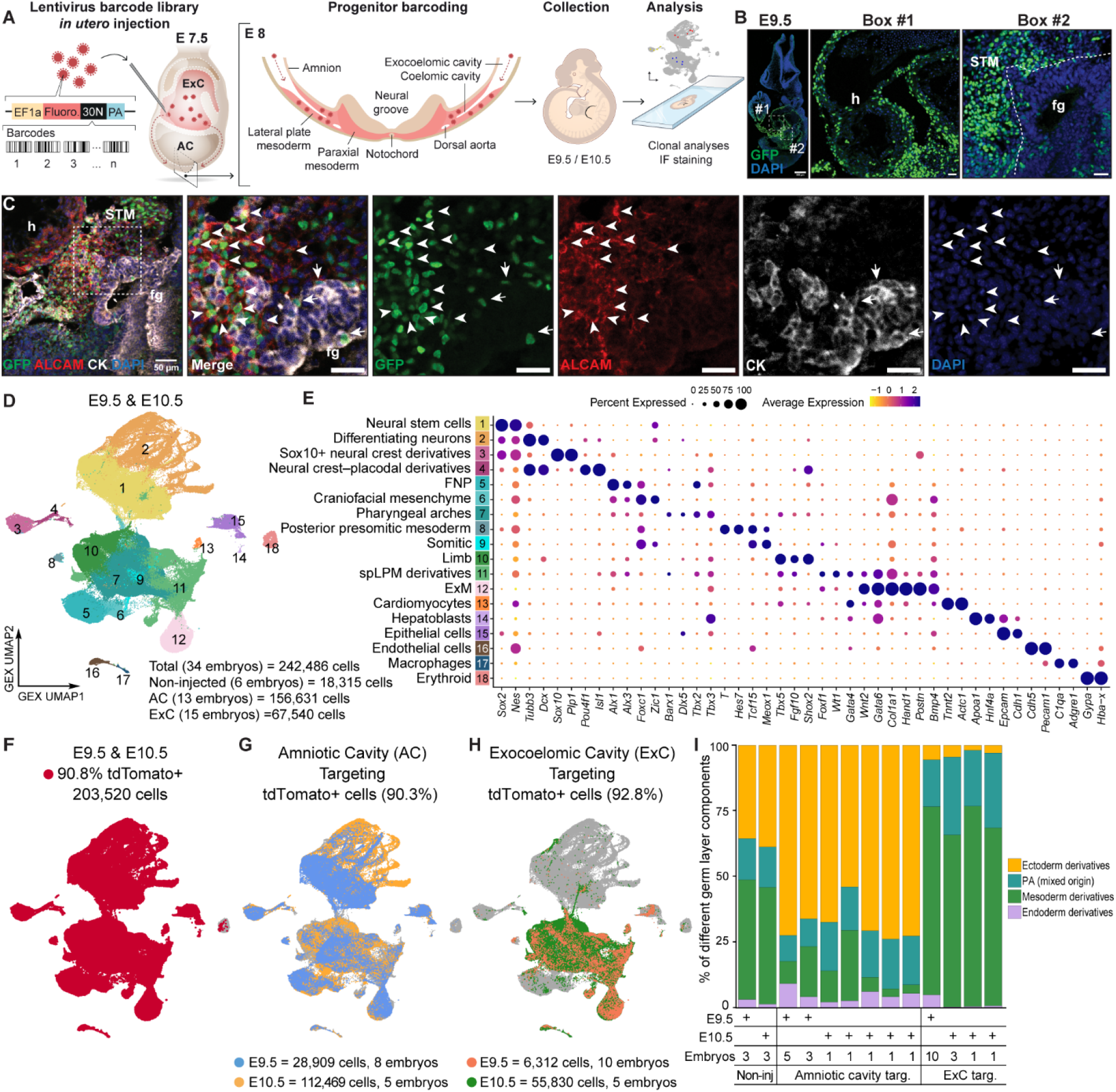
E7.5 exocoelomic cavity nano-injection targets mesodermal progenitors. **(A)** Schematic of ultrasound-guided lentiviral nano-injection into the exocoelomic cavity at E7.5, providing access to mesoderm through its continuity with the intra-embryonic coelom. **(B,C)** Representative immunofluorescence images of E9.5 embryo (sagittal view) following E7.5 exocoelomic cavity injection of H2B–GFP lentivirus, showing reporter-labeled mesodermal derivatives, including the heart and epicardium (b, box #1) and WT1⁺ liver mesothelium and ALCAM/CD166⁺ septum transversum (b, box #2, arrowheads in c, box #1)(c). Cytokeratin labels foregut epithelial cells (arrows in c). **(D)** UMAP representation of 242,486 cells collected from E9.5 and E10.5 embryos following E7.5 amniotic- or exocoelomic-cavity injection, colored by annotated cell population. **(E)** Expression of canonical markers used to annotate populations in (d). **(F)** Proportion of sorted cells expressing tdTomato RNA. **(G,H)** Numbers and relative proportions of tdTomato⁺ cells recovered from each population following amniotic-cavity (G) or exocoelomic-cavity (H) injection, shown by developmental stage. **(I)** Comparison of cell-type representation in tdTomato^+^ cells between injection routes, for different germ layer derivatives. Data comprise 34 embryos, including 13 subjected to amniotic-cavity injection and 15 subjected to exocoelomic-cavity injection. Scale bar, 300µm in b left panel, 50 µm in c left panel, and 30 µm in boxed images. AC = amniotic cavity, CK = cytokeratin, ExC = exocoelomic cavity, ExM = extraembryonic mesoderm, fg = foregut, FNP = frontonasal prominence, GEX UMAP = gene expression UMAP for single cells, h = heart, IF = immunofluorescence, spLPM = splanchnic lateral plate mesoderm, STM = septum transversum mesenchyme. See also Figure S1-S6.

To map and lineage trace targeted cells in detail, we injected the amniotic or exocoelomic cavity at E7.5 with lentivirus encoding tdTomato and a diverse library of unique 30-nucleotide barcodes^16^. Each progenitor is thus uniquely labeled with a distinct barcode that is integrated into the genome and is expressed as an RNA that can be detected with 10x Genomics single cell RNA sequencing (scRNA-seq) to resolve clonal relations with scLineage tracing ^15,16,21–23^. Embryos were collected at E9.5 or E10.5, dissociated into single cells, and tdTomato-positive cells were sorted and sequenced. After QC (**Figure S2A-F**), we obtained 242,486 cells from 34 embryos in total, with excellent read depth (average ∼6,000 genes/cell, **Figure S2E**) including 156,631 cells from 13 embryos labeled with amniotic cavity injections and 67,540 cells from 15 embryos labeled with exocoelomic cavity injections (**Figure 1D-E**). The dataset comprised ectoderm derivatives including neural stem cells and differentiating neurons from the brain and spinal cord, neural crest and its derivatives and mesenchyme derived from either neural crest or mesoderm, as well as other mesodermal derivatives (**Figure 1D,E**). More than 90% of the cells expressed tdTomato RNA (**Figure 1F**). Amniotic cavity injections enriched for neural targeting but labeled almost all cell types , with the exception of erythroid and endothelial cells (**Figure 1G,I**). In contrast, exocoelomic cavity injections led to enriched targeting of mesodermal cell types, including limb mesenchyme, pharyngeal arch mesenchyme and splanchnic lateral plate mesoderm-derived cells (**Figure 1H,I**). Both injection modes labeled extraembryonic mesoderm with variable efficiency, which we ascribe to minor experimental differences in embryo dissection rather than *de facto* targeting differences.

### In utero next-generation single-cell lineage tracing resolves mesoderm lineages

After re-clustering and further quality control, 65,654 exocoelomic cavity-labeled cells were retained for in-depth analyses (**Figure 2A,B, S3**). While limb mesenchyme was a major constituent of the dataset (44%, **Figure 2A**), heart and liver-related mesodermal cell types could be identified including *Hlx*^+^ *Gata4*^+^ septum transversum-like mesenchyme (Cluster 50, cl.50), *Wt1*^+^ *Upk3b*^+^ *Hand2*^+^ mesothelial cells (cl.511), *Wt1*^+^ *Upk3b*^+^ *Aldh1a1*^+^ epicardium (cl.15), and *Myh7*^+^ *Myl2*^+^ *Ttn*^+^ cardiomyocyte subtypes (cl.220, cl.221, cl.222).

**Figure 2.**
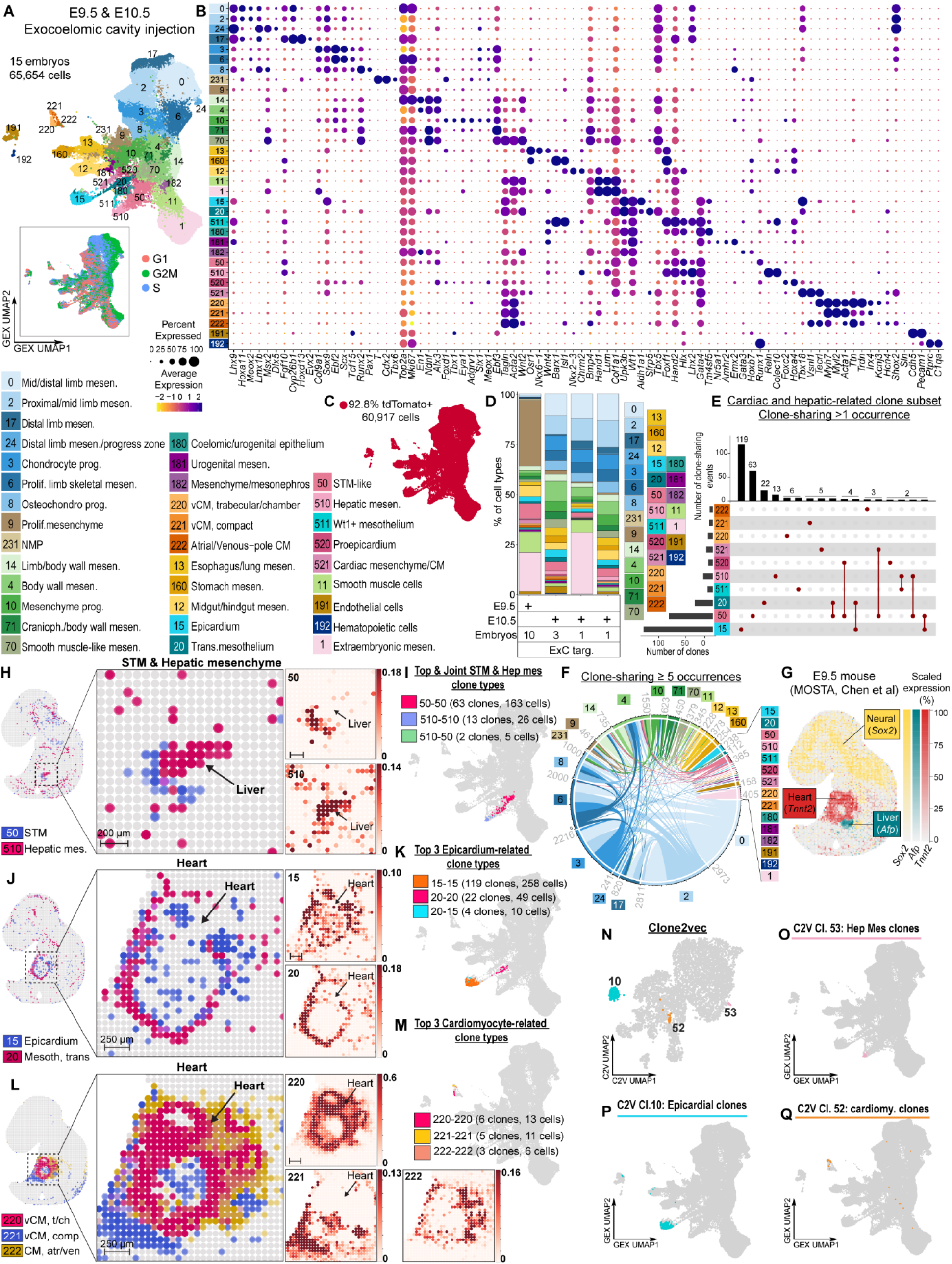
Single-cell lineage tracing resolves early mesodermal clonal relations. **(A)** UMAP representation of 65,654 cells labeled by E7.5 exocoelomic-cavity injection and collected at E9.5 or E10.5, colored by annotated cell population and cell cycle phases (boxed). **(B)** Expression of canonical markers used to annotate the populations in (a). **(C)** Proportion and distribution of sorted cells expressing tdTomato RNA. **(D)** Comparison of cell-type representation in tdTomato+ cells. **(E)** Clonal relationships among selected annotated populations. The top bars indicate the number of clone-sharing events among cell types or within a single cell type, and the left bars indicate the total clone numbers containing the indicated cell types. **(F)** Circos plot of clonal sharing (≥ 5 occurrences) for all populations at E9.5 and E10.5. Link width indicates the number of contributing cell type pairs; link color represents cell type. Grey numbers are clones per cluster, clusters without numbers have fewer than 50. **(G)** Spatial map of the published E9.5 Stereo-seq reference dataset (Chen et al, citation in text) used for transcriptomic deconvolution. **(H,I)** Spatial localization (H) and clonal relationships (I) of cl.510 hepatic mesenchyme within the liver and cl.50 septum transversum-like mesenchyme at its periphery. **(J,K)** Spatial localization (J) and clonal relationships (K) of cl.15 epicardium and cl.20 transitional mesothelium around the developing heart. **(L,M)** Spatial localization (L) and clonal relationships (M) of ventricular and atrial cardiomyocyte populations, including apically enriched compact ventricular cardiomyocytes (cl.221), centrally enriched trabecular/chamber ventricular cardiomyocytes (cl.220) and venous-pole cardiomyocytes (cl.222). **(N)** Clone2vec embedding of selected multicellular clones, colored by clonal cluster. Full clone2vec mapping is in Figure S3E,G. **(O-Q)** Composition of clonal clusters, coloured according to clonal cluster annotation in (N), containing hepatic mesenchyme (O), epicardium (P) and cardiomyocytes (Q). C2V UMAP = cell type composition UMAP for clones, Cranioph = craniopharyngeal, CM = cardiomyocytes, CM, atr/ven = cardiomyocytes, atria/venous-pole, ExC = exocoelomic cavity, GEX UMAP = gene expression UMAP for single cells, Hep mes = hepatic mesenchyme, mesen = mesenchyme, Mesoth, trans = Mesothelium, transition, MOSTA = Mouse Organogenesis Spatiotemporal Transcriptomic Atlas, from Chen et al. NMP = neuromesodermal progenitors, Osteochondro = osteochondrocyte, STM = septum transversum mesenchyme, trans.mesothelium = transitional mesothelium, vCM = ventricular cardiomyocytes, vCM, t/ch = ventricular cardiomyocytes, trabecular/chamber, vCM comp = ventricular cardiomyocytes, compact. See also Figure S3,S4.

More than 90% of the subset cells expressed tdTomato RNA (**Figure 2C**), and we identified 5,371 multicellular clones based on shared barcode expression^16^, encompassing 20,760 cells (**Figure S4A-C**). As expected, based on cell type abundance, the most common clone types comprised of limb mesenchyme populations (**Figure 2D**, **Figure S4D**). Focused analysis of cardiac and hepatic-related clones showed that these cell types predominantly shared clones with themselves (**Figure 2E,F**). Deconvolution of the cell types on published E9.5 Stereo-Seq data ^24^ (**Figure 2G**), using Stereoscope^25^, corroborated intra-hepatic localization of cl.510 hepatic mesenchyme and liver-peripheral localization of cl.50 septum transversum-like mesenchyme (STM) (**Figure 2H**), while clonal analysis showed that these populations predominantly shared clones with themselves (13 hepatic mesenchyme clones and 63 STM clones), with only two joint clones (**Figure 2E,I**). Deconvolution placed the cardiac cl.20 transitional mesothelial cells at the heart edges, while cl.15 epicardium enveloped the heart and exhibited a broader signal (**Figure 2J**). Similar to STM, epicardium predominantly shared clones with itself (119 cl.15 epicardium clones), as did the transitional mesothelium (22 cl.20 trans.mesothelium clones), and there were only four clones shared between epicardium and transitional mesothelium (**Figure 2E,K**). Deconvolution revealed spatial enrichment of the three cardiomyocyte clusters, with cl.221 compact ventricular cardiomyocyte enrichment at the heart apex, enrichment of cl.220 trabeculated/chamber ventricular cardiomyocytes more centrally, and enrichment of cl.222 atrial or venous pole cardiomyocytes at the venous pole (**Figure 2L**). These cell types also predominantly shared clones with themselves (**Figure 2E,M**).

Finally, clone2vec analysis, which represents each clone by its cell-type composition and visualizes similarly composed clones close together in a UMAP, corroborated that limb-related clones constituted a large proportion of clones at this timepoint (**Figure S4E-G**), and identified three clonal types for hepatic mesenchyme (**Figure 2N,O**), epicardium (**Figure 2N,P**), and cardiomyocytes (**Figure 2N,Q**).

Clonal analyses of the cells barcoded with amniotic cavity injections corroborated our previously published work^3^ finding early compartmentalization of neural lineages (**Figure S5,S6**) and showed that mesenchyme of both mesodermal and neural crest origin could be labeled and clonally analysed, with an enrichment for neural crest derivatives compared to exocoelomic cavity injections (**Figures 1I,S4,S5,S6**).

In conclusion, *in utero* exocoelomic cavity barcode labelling targets mesoderm shortly before giving rise to septum transversum mesenchyme, cardiomyocytes, and the epicardium, resulting in separate and distinct clonal relations. We therefore next applied *in utero* next generation scLineage tracing to query hepatic and cardiac mesenchymal lineages during organ development and mesenchyme diversification.

### In utero nano-injection at E7.5 labels mesodermal derivatives in E16.5 liver

Liver formation is induced when septum transversum mesenchyme signals to adjacent foregut endoderm to adopt a hepatic fate ^26^. Subsequently, this mesenchymal population envelops the liver and gives rise to mesothelium encapsulating the liver. *Wt1-Cre* and *Mesp1-Cre* lineage tracing indicate that this mesothelium gives rise to stellate cells and perivascular fibroblasts^8,9^, although their precise lineage relationship remains unresolved. The liver is also extensively vascularized by mesoderm-derived endothelial cells, and is a major site of fetal hematopoiesis. The fetal liver thus contains a multitude of mesodermal derivatives including hematopoietic cells, endothelial cells, mesothelial cells, fibroblasts, and hepatic stellate cells ^7^, whose lineages we next aimed to resolve at high resolution with *in utero* next-generation scLineage tracing.

We injected exocoelomic cavities at E7.5 with the barcode lentivirus, and collected livers at E16.5 (**Figures 3A,S7)**. To identify cell types with high confidence, we performed spatial transcriptomics of E16.5 livers for cell type deconvolution and spatially aware annotation (**Figures 3A,S8)**. In the ExC-injected condition, the majority of the 155,747 cells were mesoderm derivatives (99.5%, **Figure 3B-D**). Hematopoietic cells including hematopoietic stem/progenitor cells, erythroid, myeloid, and B cells constituted 63.6% (99,024 cells), while mesenchymal cells, including mesothelial cells, fibroblasts, vascular smooth muscle cells and stellate cells, constituted 25.8% (40,221 cells). 88% of cells in targeted livers were positive for tdTomato RNA (**Figure 3E**), and exocoelomic cavity injection enriched for targeting of mesothelial and mesenchymal cells (Cl.2 and 3 were enriched 8-fold relative to non-injected liver, **Figure 3E,F**), predominantly at the expense of erythroid cells. Their reduced, but not absent, labelling is consistent with the strong contribution of the inaccessible first wave of yolk-sac haematopoiesis (generating primitive erythroid, megakaryocyte and macrophage lineages), with residual labelling potentially arising from later hematopoietic waves that are accessible to exocoelomic cavity injection via labelling of hemangioblastic endothelium and multipotent hematopoietic progenitors. Immunofluorescence imaging of E16.5 livers, from embryos injected at E7.5 into the exocoelomic cavity with a lentivirus encoding a histone-2B-GFP reporter, confirmed enriched targeting of Podoplanin-positive (PDPN+) mesothelial cells, ALCAM-positive submesothelial cells, Desmin-positive mesenchymal cells and perivascular fibroblasts/portal fibroblasts (**Figure 3G**).

**Figure 3.**
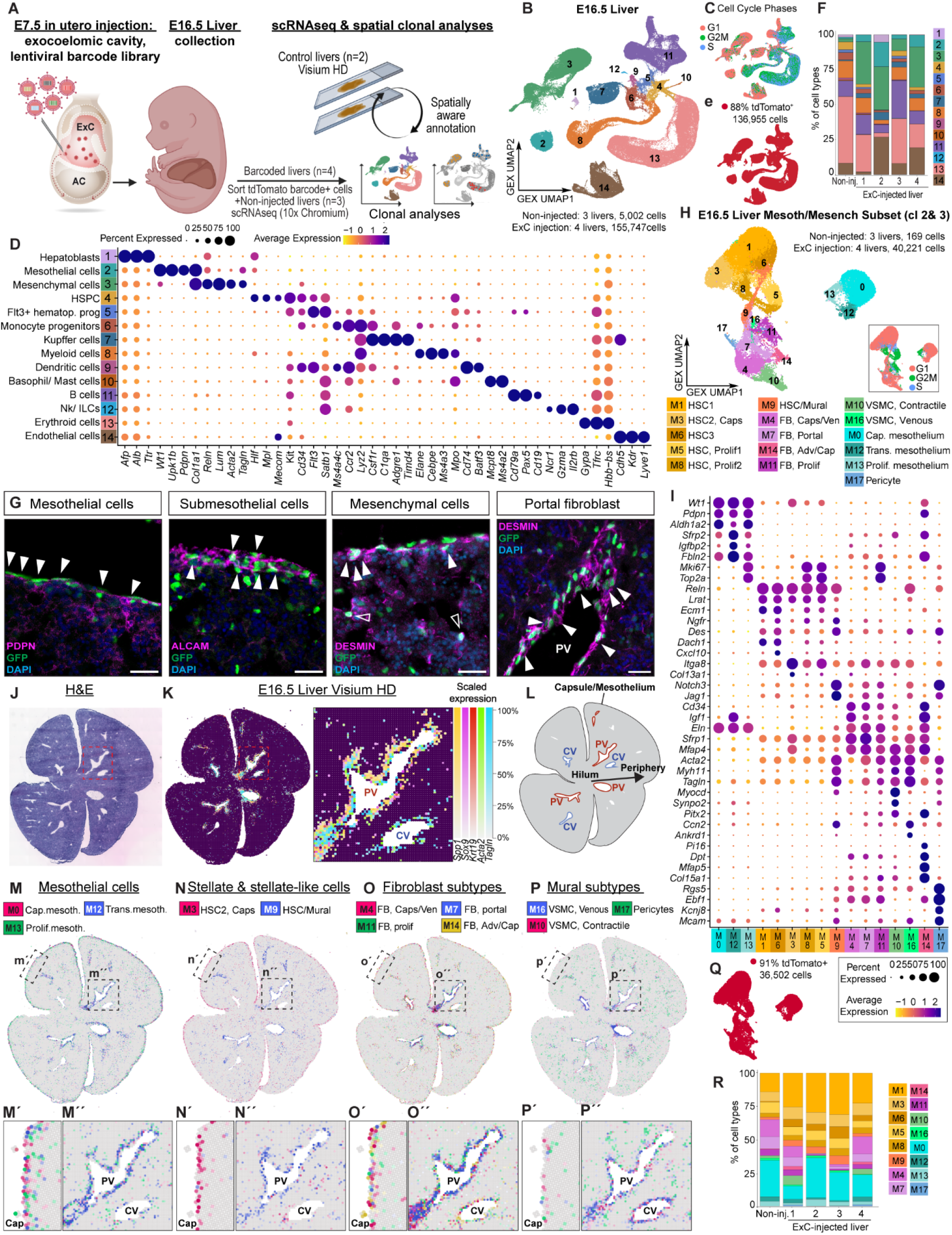
Exocoelomic cavity injection targets spatially organized mesenchymal populations in the fetal liver. **(A)** Experimental workflow. The exocoelomic cavity was injected at E7.5 with barcode lentivirus, and tdTomato⁺ cells were flow-sorted from E16.5 livers for scRNA-seq and lineage tracing. Visium HD spatial transcriptomics was performed to localize the identified populations. **(B,C)** UMAP representation of 160,749 cells from injected and non-injected E16.5 livers, colored by annotated cell population (**B**) and cell cycle phases (**C**). **(D)** Expression of canonical markers used for cell-type annotation. **(E)** Numbers and distribution of tdTomato⁺ cells. **(F)** Cell-type representation in tdTomato^+^ cells across experimental conditions and biological replicates. **(G)** Representative immunofluorescence images of E16.5 liver following E7.5 exocoelomic-cavity delivery of lentivirus encoding H2B–GFP. **(H)** UMAP representation of 40,390 mesothelial and mesenchymal cells after subsetting and reclustering, colored by subcluster. **(I)** Expression of markers defining populations in **h**. **(J-L)** Visium HD analysis of E16.5 liver, showing tissue morphology in Hematoxylin and Eosin staining (H&E, J), spatial domains (K) including portal veins (PV) surrounded by *Spp1*+ *Sox9*+ *Krt19*+ ductal plate and central veins (CV), and annotation of structures in liver sections (L). **(M)** Spatial localization of capsular mesothelium (M0), transitional portal-associated mesothelium (M12) and proliferating mesothelium (M13). **(N)** Spatial distributions of hepatic stellate cell populations, including capsular M3 stellate cells and periportal stellate/mural-like M9 cells. **(O)** Localization of fibroblast populations with capsular, perivascular or portal enrichment, including M4, M7, M11 and M14 fibroblasts. **(P)** Spatial distributions of mural populations, comprising peri-venous M16 VSMCs, capsular/peri- venous M10 VSMCs and parenchymal M17 pericytes. **(Q)** Numbers and distribution of tdTomato⁺ cells within the mesothelial and mesenchymal subset. **(R)** Relative abundance of mesenchymal subpopulations across injected biological replicates and non-injected controls. Scale bars, 30 µm. Cap = capsule, FB = fibroblasts, HSC = hepatic stellate cells, HSPC = hematopoietic stem and progenitor cells, Nk/ILCs = natural killer/innate lymphoid cells, Prolif. = proliferating, Trans. = Transitional, VSMC = vascular smooth muscle cells. See also Figure S7-S10.

Subsetting, re-embedding, and re-clustering the mesenchymal clusters (Cl. 2 and 3) revealed several subpopulations of mesothelial cells, hepatic stellate cells, smooth muscle cells and fibroblasts (**Figure 3H,I**). We performed high-resolution spatial transcriptomics on E16.5 mouse liver using Visium HD and deconvolved the spatial data with Stereoscope^25^, using cell types defined by the scRNA-seq (**Figures 3J-L**,**S9,S10**). The three mesothelial cell subtypes expressed distinct genes and occupied different anatomical locations, with proliferating *Wt1*^+^ *Pdpn*^+^ *Mki67*^+^ mesothelial cells (Mesenchyme subcluster 13, M13, Prol.mesothelium), and bona fide *Wt1*^+^ *Pdpn*^+^ *Aldh1a2*^+^ mesothelial cells (M0, Cap.mesothelium) predominantly located in the capsule layer enveloping the liver, while transitional *Wt1*^+^ *Pdpn*^+^ *Fbln2^+^* mesothelial cells (M12, Trans.mesothelium), were enriched around the portal vein (**Figure 3M**). Six subclusters of *Reln*^+^ hepatic stellate cell-like cells were identified, of which one (M3) exhibited a capsular localization (M3, *Reln*^+^ *Lrat*^+^ *Itga8*^+^ HSC2, Caps, **Figure 3N**), while M9 *Reln*^+^ *Des*^+^ *Ngfr*^+^ *Notch3*^+^ *Jag1*^+^ *Acta2*^+^ *Tagln*^+^ HSC and mural-like cells were enriched in the peri-portal region (**Figure 3N**). Two were cycling (M5 and M8), of which one (M5) occupied a capsular localization (**Figure S9,S10**) while the other exhibited a broad hepatic distribution. The remaining two *Reln*^+^ *Lrat*^+^ stellate cell subclusters (M1 and M6) were differentially enriched for *Ngfr* and *Cxcl10*, and were broadly distributed throughout the liver (**Figure S9,S10**). Similar to both mesothelial cells and stellate cells, we identified multiple fibroblast populations, with preferential capsular or periportal enrichment, including *Cd34*^+^ *Igf1*^+^ *Mfap4*^+^ *Col15a1*^-^M4 cells occupying both the capsular and peri-venous regions, while M7 *Sfrp1*^+^ *Mfap4*^+^ fibroblasts were enriched around portal veins (**Figure 3O**). In contrast, M14 exhibited features of *Pi16*⁺ adventitial fibroblasts (*Pi16*, *Cd34*)^27^ and *Col15a1*⁺ basement membrane-associated subtypes (*Col15a1*), together with the pan-universal fibroblast marker *Dpt*, while retaining capsular identity through *Wt1* and *Pdpn* expression. Nonetheless, their marker expression is reminiscent of portal fibroblasts with mesenchymal stem cell features, described by Lei et al^28^. Both M14 and *Mki67*^+^ *Top2a*^+^ M11 proliferating fibroblasts were present in the capsular and peri-venous regions (**Figure 3O**). Finally, there were three subtypes of mural cells (in addition to the mural-HSC-like M9), with two types of *Myh11*^+^ *Tagln*^+^ vascular smooth muscle cells, including *Ccn2*+ *Ankrd1*^+^ M16 peri-venous VSMCs, and capsular/peri-venous M10 VSMCs expressing contractility-associated *Myocd* and *Synpo2*, as well as *Rgs5*^+^ *Ebf1*^+^ *Kcnj8*^+^ *Mcam*^+^ M17 pericytes, which were scattered throughout the parenchyma (**Figures 3P**,**S9,S10**). In the mesenchymal subset, 91% expressed tdTomato RNA (**Figure 3Q**), and the proportions across cell types were largely similar across replicates and compared to non-injected mice (**Figure 3R**). We therefore next assessed lineages for these cell types, based on clonal sharing of barcode expression.

### Clonal architecture reveals distinct routes of liver mesenchymal differentiation

There were 3,887 multicellular clones in the liver data, comprising 93,180 cells (**Figure S11A-C**). While the mean clone size was 24 cells, the median clone size was 3, a disparity that was driven by the presence of some “mega-clones” composed of >1,000 cells of the hematopoietic or endothelial lineage (**Figure S11D,E**). The few but exceptionally large hematopoietic clones are consistent with the extensive expansion of hematopoietic cells that occurs in the fetal liver.

The most prevalent clones were M1 hepatic stellate cell clones (202 clones, 477 cells) and M0 capsular mesothelium clones (199 clones, 683 cells; **Figure 4A-C**), while the most prevalent multicellular clone types spanning multiple populations comprised stellate cell subtypes (**Figure 4A,B**). In each class (HSC, mesothelium, fibroblasts, mural cells), self-sharing clones were the most common clone type (**Figure 4B-E**). Mesenchymal and hematopoietic cells displayed distinct clonal architectures (**Figure 4A**).

**Figure 4.**
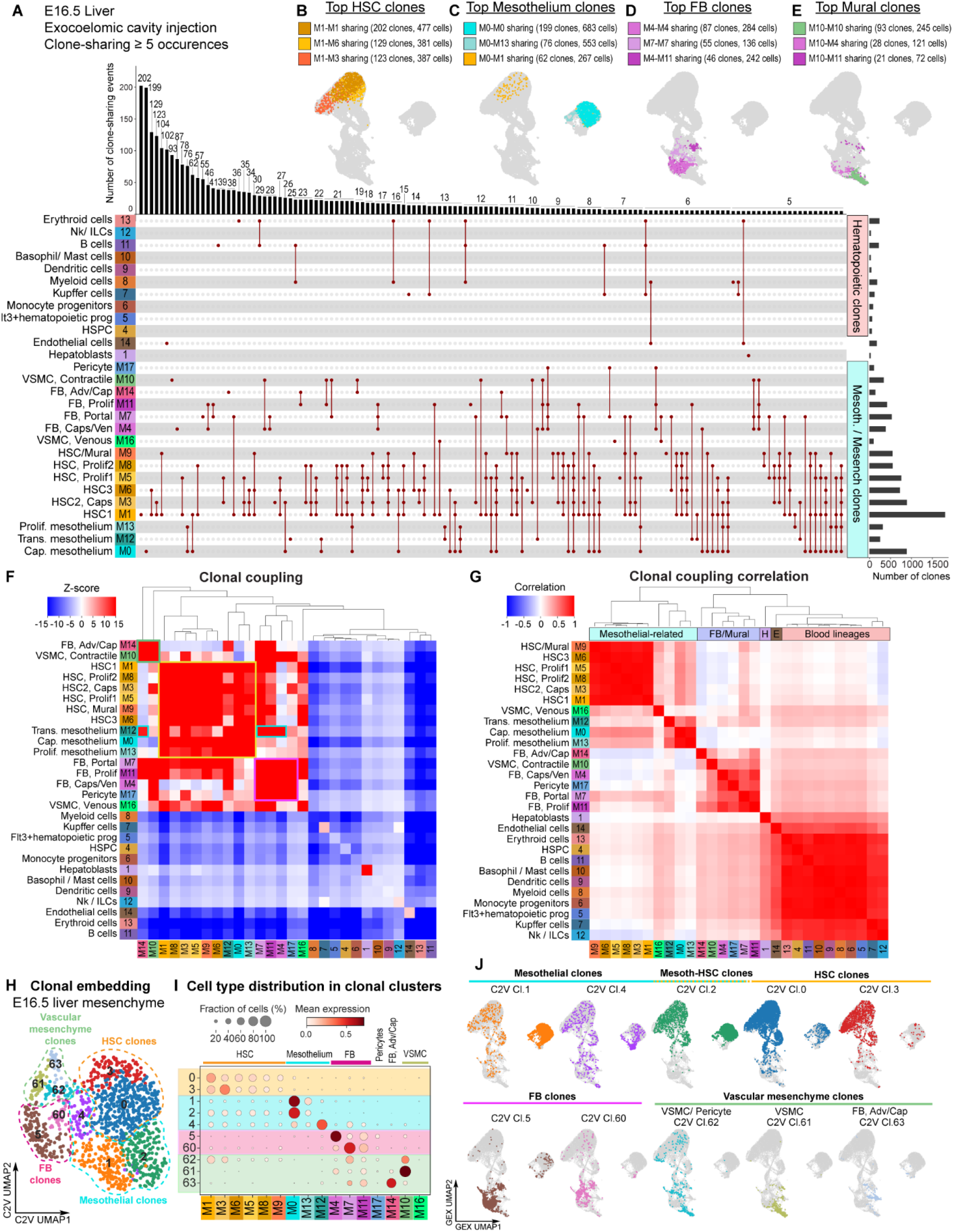
Clonal architecture reveals lineage-biased diversification of fetal liver mesenchyme. **(A)** plot depicting clonal relationships among E16.5 liver cell populations following E7.5 exocoelomic-cavity barcode labelling, occurring ≥ 5 times. Analyses comprise 3,887 multicellular clones encompassing 93,180 cells. **(B-E)** Clonal composition of most prevalent hepatic stellate cell (B), mesothelial (C), fibroblast (D) and mural cell (E) populations. **(F)** Clonal coupling analysis of liver mesenchymal populations. Clonal coupling Z-scores quantify the enrichment of observed barcode sharing relative to randomized datasets preserving cell-population abundances, with a positive (red) score indicating enriched coupling and a negative (blue) score indicating under-represented coupling. **(G)** Correlation of clonal coupling profiles across liver cell populations. **(H)** Clone2vec (C2V) embedding of multicellular liver clones (clone size ≥5), colored by clonocluster. **(I)** Relative contributions of annotated cell populations to each clonocluster. Dot size represents the fraction of clones containing the indicated cell type. Dot color represents the average cell-type composition of clones within each C2V clonocluster. **(J)** Mesenchymal composition of clone2vec-defined clonoclusters. Adv=adventitial, Caps= capsular, FB = fibroblasts, HSC = hepatic stellate cells, HSPC = hematopoietic stem and progenitor cells, Ven = venous. See also Figure S11.

Clonal coupling analysis ^29^ of the entire liver dataset revealed strong coupling patterns between multiple mesenchymal cell types, in particular HSCs and mesothelial cells (**Figure 4F**). Although hematopoietic cells were not significantly clonally coupled to any cell types due to their relative over-abundance in the dataset, clonal coupling correlation corroborated that all hematopoietic lineages were linked to one another, and further demonstrated strong correlations for HSC subtypes including M9 HSC/mural cells (**Figure 4G**). Of note, M0 and M12 mesothelial cells exhibited preferential coupling to different mesenchymal subsets: M0 mesothelial cells were strongly coupled to HSCs while M12 mesothelial cells were more strongly coupled to fibroblast subtypes, and were less strongly coupled to HSC subtypes (**Figure 4F**), indicating that mesothelial heterogeneity contributes to lineage diversification. Furthermore, while M17 pericytes and M10 VSMCs were strongly coupled to one another (**Figure 4F**), with correlated clonal architectures (**Figure 4G**), M16 VSMCs were not strongly coupled to either M10 or M17, and instead M16 VSMCs were strongly associated with HSCs (**Figure 4F,G**), suggesting mural cells arise through two differentiation pathways during liver development. These conclusions were corroborated by clone2vec analyses, demonstrating that the predominant clone types included mesothelial cells and/or HSCs (**Figures 4H-J, S11F-H**), and that M0 and M12 mesothelial cells exhibited biased contribution to clones, with M0 mesothelial cells more abundant in HSC-related clones, and M12 mesothelial cells more associated with fibroblast- or mural cell-containing clones.

Together, these findings identify distinct mesothelial populations as lineage-biased sources of liver mesenchymal diversity and reveal that vascular smooth muscle cells arise through separable pericyte-associated and stellate-cell-associated developmental routes.

### Exocoelomic cavity injection and high-resolution spatial transcriptomics resolve cardiac cellular diversity

The heart arises largely from mesodermal progenitors in the first and second heart fields and the proepicardium, with neural crest contributions to the outflow tract, valves, and great vessels. Cardiac development begins around E6.5, when mesodermal progenitors migrate through the primitive streak, forming the bilateral cardiac crescent by E8.25. The first heart field has been described to contribute to the left ventricle and parts of the atria, whereas the second heart field is thought to give rise to the right ventricle, outflow tract, and additional atrial regions. The proepicardium, a mesenchymal structure arising near the septum transversum mesenchyme, generates the epicardium, from which mesenchymal cells emerge and migrate into the heart. Although these cells are proposed to contribute to fibroblasts, smooth muscle cells, and other intracardiac mesenchymal populations, including valve mesenchyme and possibly cardiomyocytes, their exact lineage contributions remain unresolved. Here, we directly address cardiac lineages using *in utero* single-cell lineage tracing to define the developmental fate of the first and second heart fields, as well as proepicardial and neural crest derivatives.

We injected the exocoelomic cavity at E7.5 with the barcode lentivirus, and collected transduced cells from four E16.5 hearts (**Figures 5A,B, S12, S13A-C**). Clusters were annotated using published human and mouse cardiac scRNA-seq and spatial atlases ^24,30–33^. To validate cell type annotations and localize defined and emerging populations within the tissue context, we performed Visium HD spatial transcriptomic analysis of E16.5 hearts (**Figures 5A,C, S14**). Using the scRNA-seq as a reference, spatial deconvolution resolved spatially restricted atrial and ventricular cardiomyocyte subsets alongside fibroblasts, smooth muscle cells, pericytes and epicardium (**Figures 5D**, **S15**).

**Figure 5.**
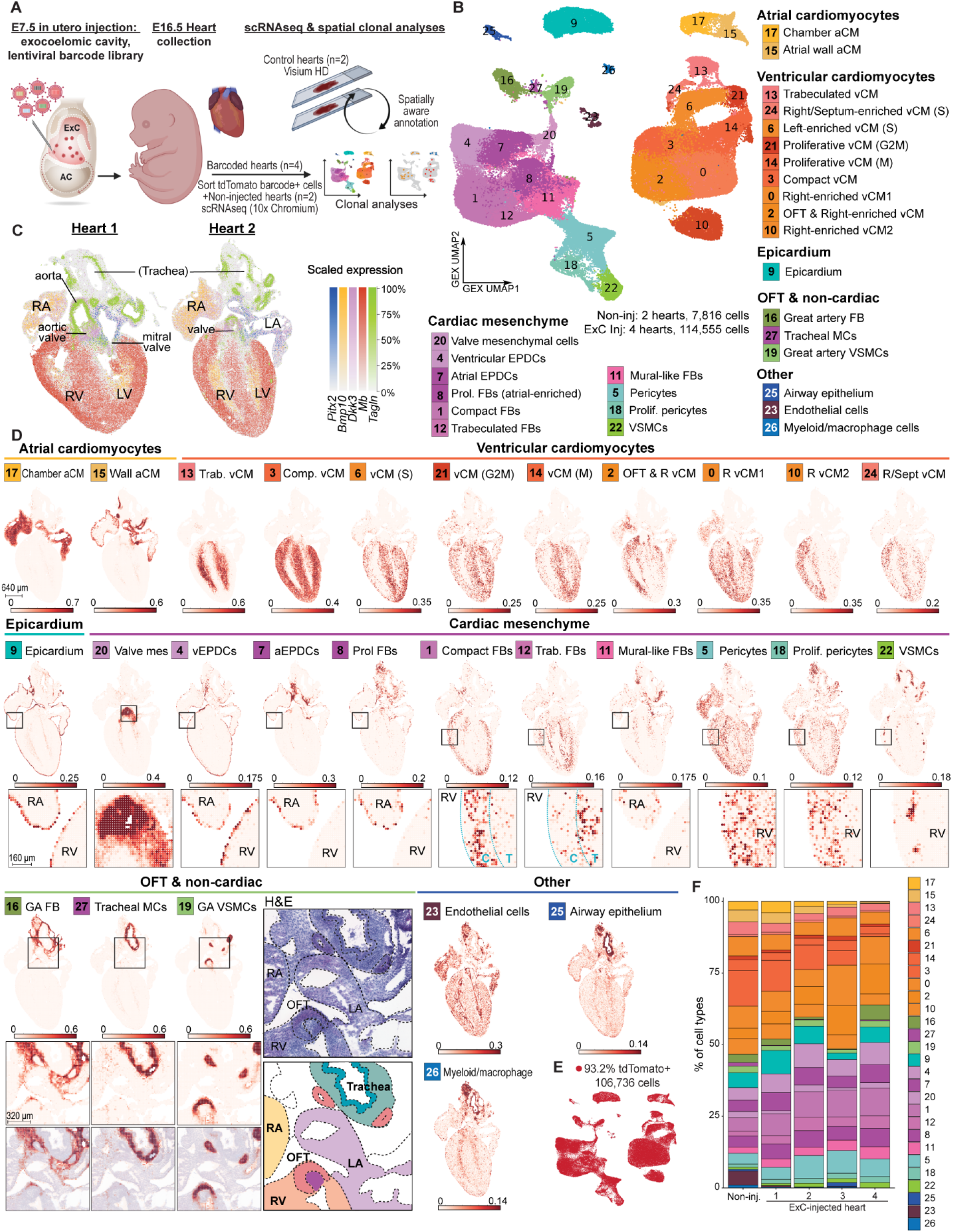
Single-cell and spatial transcriptomics resolve cardiac cell diversity following E7.5 exocoelomic-cavity labelling. **(A)** Experimental workflow. The exocoelomic cavity was injected at E7.5 with barcode lentivirus, and tdTomato⁺ cells were isolated from E16.5 hearts for scRNA-seq and lineage tracing. Visium HD spatial transcriptomics was performed on E16.5 hearts to localize the identified populations. **(B)** UMAP representation of 122,371 cells from four injected and two non-injected E16.5 hearts, colored by annotated cell population. **(C)** Tissue morphology and spatial domains resolved by Visium HD analysis of two E16.5 hearts, with key markers resolving major anatomical hallmarks. **(D)** Spatial deconvolution using the scRNA-seq dataset as a reference, showing the localization of all populations. **(E)** Proportion and distribution of sorted cells expressing tdTomato RNA following exocoelomic-cavity injection. **(F)** Numbers and relative proportions of tdTomato⁺ cells within each population, shown for individual biological replicates. A = atrial, C = compact region, CM = cardiomyocyte, EPDC= epicardium-derived cells, ExC = exocoelomic cavity, G2M = G2M phase of cell cycle, GA = great artery, LA = left atrium, LV = left ventricle, MCs = mesenchymal cells, mes = mesenchyme, OFT = outflow tract, RA = right atrium, RV = right ventricle, S = S phase of cell cycle, Sept = septal, T, trab = trabecular region, v = ventricular, VSMCs = vascular smooth muscle cells. See also Figure S12-S15.

*Tbx5*^+^ *Sln*^+^ *Myl7*^+^ atrial cardiomyocytes comprised *Myom2*^+^ *Nppa*+ atrial working chamber cardiomyocytes (cl.17), and *Cacna1d*^+^ *Nid2*^+^ conductive atrial wall cardiomyocytes (cl.15) lining the atrial surface. *Myl2*^+^ *Myh7*^+^ ventricular cardiomyocyte subtypes included cl.3 *Kcne1*+ *Cited4*+ compact cardiomyocytes and cl.13 *Scn5a*^+^ *Irx3*^+^ trabecular cardiomyocytes. There were four right-side enriched populations: *Myom2*⁺ *Nppb*⁺ *Lmod2*⁺ cardiomyocytes localized to the right ventricle and venous pole (cl.2); *Kcnq1*⁺ *Myh7b*⁺ cardiomyocytes showed a weaker right-ventricular enrichment (cl.0); and two *Ddc*⁺ populations comprised strongly right-ventricle-enriched cardiomyocytes (cl.10) and *Mki67*⁺ *Plk1*⁺ cycling cardiomyocytes enriched in the right ventricle and interventricular septum (cl.24). Three additional cycling populations represented successive cell-cycle states, including S phase, G2/M and mitosis (cl.6, cl.21 and cl.14, respectively; **Figure S13B**), with cl.6 preferentially localized to the left ventricle.

*Msln*^+^ *Rspo1*^+^ *Upk3b*^+^ epicardial cells (cl.9) formed the expected outer lining of the heart, beneath which ten spatially organized mesenchymal populations were resolved. These included cl.20 *Lef1*^+^ *Meox1*^+^ *Scx*^+^ valve mesenchyme, and three sub-epicardial fibroblast populations: ventricle-enriched *Wt1*^+^ *Tcf21*^+^ *Lrat*^+^ *Dpep1*^+^ epicardium-derived cells (EPDCs; cl.4), atrium-enriched *Wt1*^+^ *Tcf21*^+^ *Moxd1*^+^ *Tnc*^+^ EPDCs (cl.7), and proliferating *Mki67*^+^ *Gria4*^+^ fibroblasts concentrated in the atrial subepicardium (cl.8). Two *Col15a1*+ fibroblast populations segregated between ventricular myocardial compartments, with *Kcnn3*^+^ *Scara5*^+^ fibroblasts (cl.1) associated with compact myocardium and *Lrrc1*^+^ *Gria4*^+^ (cl.12) with trabecular myocardium. Four mural-like populations were also distinguished: *Col24a1*+ *Egr3*+ *Rgs5*^+^ *Pdgfrb*^+^ *Tagln*^+^ mural-like fibroblasts (cl.11) localized predominantly to the atria and ventricular trabecular myocardium; *Rgs5*^+^ *Pdgfrb*^+^ pericytes (cl.5) were distributed throughout the heart; proliferating *Mki67*^+^ *Rgs5*^+^ *Pdgfrb*^+^ pericytes (cl.18) were enriched in compact myocardium; and mature *Rgs5*^+^ *Tagln*^+^ *Myh11*^+^ VSMCs (cl.22) encircled coronary arterial vessels.

Notably, labeled cells included two spatially confined great-artery populations: *Mfap5*^+^ *Col12a1*^+^ great artery fibroblasts (cl.16) and *Tagln*^+^ *Lmod1*^+^ great artery smooth muscle cells (cl.19), which originate from neural crest^34^ (**Figures 5B-D, S15**). This labelling is consistent with the timing of neural crest migration into the heart and demonstrates that exocoelomic cavity injection accesses migratory mesoderm- and neural crest-derived mesenchymal progenitors before their final tissue integration. The dataset also included adjacent *Wif1*^+^ *Tbx4*^+^ tracheal mesenchymal cells (cl.27), and *Epcam*^+^ *Krt18*^+^ tracheal/airway epithelium (cl.25), as well as *Kdr*^+^ *Cdh5*^+^ endothelial cells (cl.23) and *Adgre1*^+^ *Csf1r*^+^ myeloid cells (cl.26). Overall, 93.2% of sorted cells expressed tdTomato (**Figure 5E**), and targeting was highly reproducible across replicates (**Figure 5F**). We therefore proceeded to reconstruct the clonal development of cardiac cell types.

### Clonal architecture reveals early myocardial segregation and epicardial diversification

There were 4,918 multicellular clones encompassing 66,799 cells (**Figures 6A,B, S13D-F)**, of five main clonal architectures (**Figures 6C, S13H-J**) including, in order of clonal abundance: ventricular cardiomyocyte-related clones, mesoderm/epicardium-related clones, atrial cardiomyocyte-related clones, neural crest-related clones and myeloid-related clones. These data corroborate findings that atrial and ventricular cardiomyocytes arise from distinct fate-biased progenitors, present already at E7.5^19,35^.

**Figure 6.**
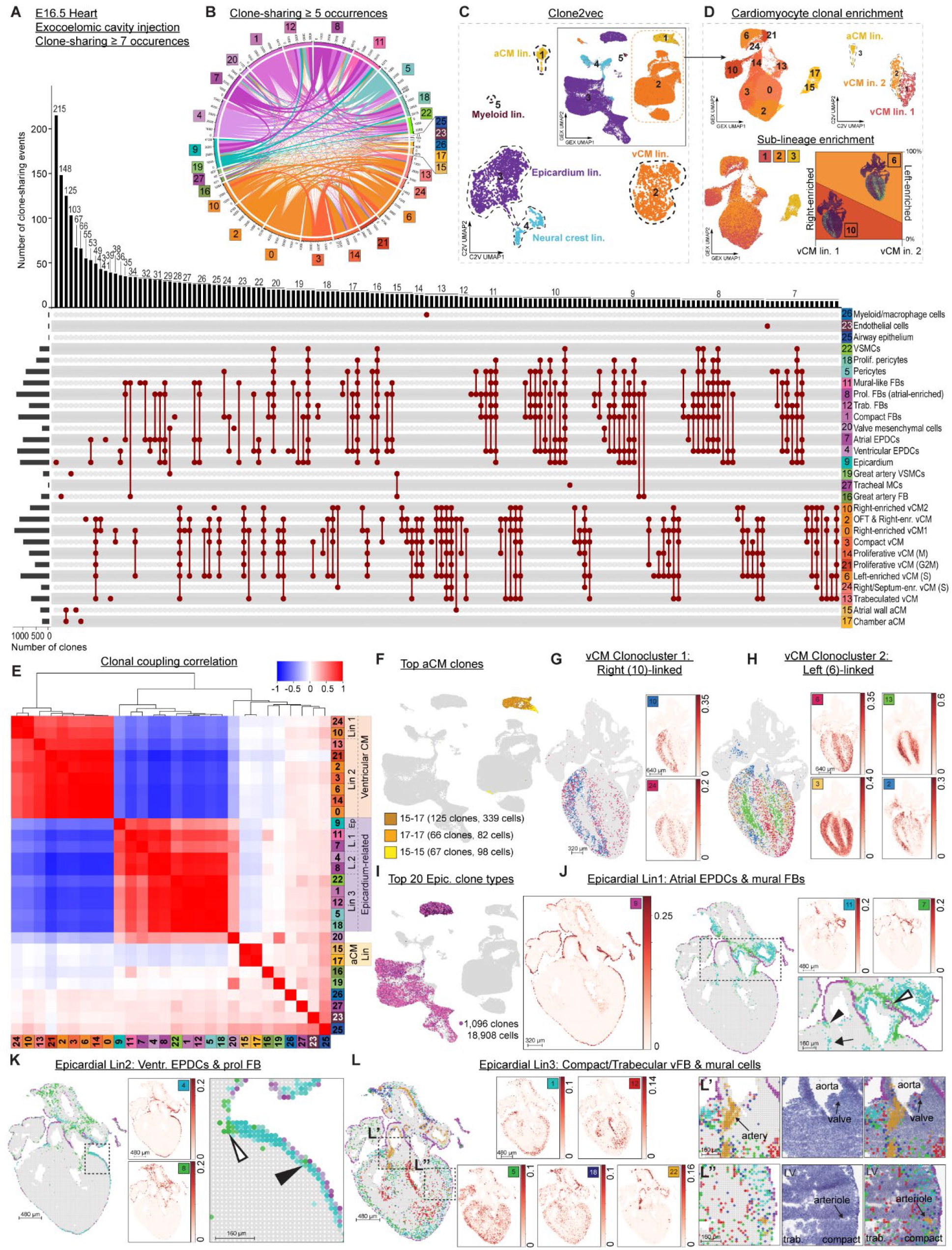
Clonal architecture reveals early myocardial segregation and spatially organized epicardial diversification. **(A)** UpSet plot depicting clonal relationships among cell populations in E16.5 hearts following E7.5 exocoelomic-cavity barcode labelling, occurring ≥7 times. In total, 4,918 multicellular clones encompassed 66,799 cells. Connections indicate multicellular clones shared between populations. **(B)** Circos plot of clonal sharing for all populations. **(C)** Clone2vec (C2V) embedding of cardiac clones. The clone2vec embedding is projected onto the parental gene expression (GEX) UMAP, identifying overarching clonally linked populations (boxed). **(D)** Clone2vec analysis after subsetting cardiomyocyte-containing clones. Top left = Subsetting and re-embedding of cardiomyocyte clusters defined in Figure 5B, labeled with their original annotation. Bottom left = C2V embedding of cardiomyocytes. Top right = projection of C2V clonoclusters onto the GEX UMAP. Bottom right = two ventricular clonoclusters were resolved: clonocluster 1 was enriched for right-biased cl.10 cardiomyocytes, whereas clonocluster 2 was enriched for left-biased cl.6 cardiomyocytes. **(E)** Correlation of population-level clonal-coupling profiles. Underlying clonal coupling analysis can be found in Figure S14G. **(F)** Clonal composition of most prevalent atrial cardiomyocyte clones. **(G)** Focused clonal relationships of vCM Clonocluster 1 from (D), corresponding to Lineage 1 in (E). **(H)** Focused clonal relationships of vCM Clonocluster 2 from (D), corresponding to Lineage 2 in (E). **(I)** Clonal relationships of the top 20 most prevalent clone types linking epicardium to intracardiac fibroblast and mural populations. **(J)** Atrial epicardial-derived trajectory comprising cl.7 atrial EPDCs and cl.11 mural-like fibroblasts. **(K)** Ventricular epicardial trajectory comprising cl.4 ventricular EPDCs and cl.8 proliferating fibroblasts. **(L)** Intramyocardial fibroblast-mural trajectory linking compact- and trabecular-region fibroblasts (cl.1 and cl.12) with pericyte and VSMC populations (cl.5, cl.18 and cl.22). Valve mesenchyme (cl.20) lacked a dominant clonal association and showed weaker coupling to both epicardium-related populations and great-artery VSMCs (cl.19). a = atrial, C2V = clone2vec umap (clones), CM =cardiomyocytes, FB = fibroblasts, GEX = gene expression umap (cells), MCs = mesenchymal cells, v = ventricular. VSMCs = vascular smooth muscle cells. See also Figure S16.

Ventricular cardiomyocyte clones formed a coherent group in the clone2vec embedding, suggesting that E7.5 progenitors span a continuum of competence across ventricular lineages rather than exhibiting a strict left-right fate restriction. To resolve potential biases within this continuum, we subset and re-embedded all cardiomyocytes and repeated clone2vec analysis at a moderately increased clustering resolution. We used the most strongly left- and right-enriched ventricular cardiomyocyte populations (cl.6 and cl.10, respectively) as spatial proxies for assessing clonal laterality (**Figure 6D**). This analysis resolved two clonoclusters: clonocluster 1 was enriched for right-biased cl.10 cardiomyocytes, whereas clonocluster 2 was enriched for left-biased cl.6 cardiomyocytes. Thus, ventricular progenitors exhibit a subtle but detectable left-right clonal bias embedded within a broader continuum of ventricular competence.

Clonal coupling correlations further supported a moderate lineage bias between cl.6 and cl.10, while confirming the pronounced segregation of atrial and ventricular cardiomyocyte lineages (**Figures 6E,F, S13G**). Cluster 10 right-ventricular cardiomyocytes were most strongly coupled to Cluster 24 proliferating right-enriched ventricle cardiomyocytes (**Figure 6E,G**). In contrast, Cluster 6 left-ventricular cardiomyocytes were clonally coupled to all other ventricular cardiomyocytes, including cl.2 outflow tract and right-ventricle cardiomyocytes, cl.13 trabecular ventricular cardiomyocytes and cl.3 compact ventricular cardiomyocytes (**Figure 6E,H**). Some clonal coupling between atrial and ventricular cardiomyocytes was present (**Figure 6**, 13 clones consisting of 32 cells connect Cluster 15 to Cluster 2), indicating that progenitors contributing to atrial cardiomyocytes may also generate OFT ventricular cardiomyocytes.

Clonal coupling linked all intracardiac mesenchymal populations, except valve mesenchyme, to the epicardium and resolved three spatially organized sub-lineages (**Figure 6E,I**): cl.7 atrial EPDCs with cl.11 mural-like fibroblasts (**Figure 6E,J**); cl.4 ventricular EPDCs with cl.8 proliferating fibroblasts (**Figure 6E,K)**; and compact- and trabecular-region fibroblasts (cl.1 and cl.12) with mural populations cl.5, cl.18 and cl.22 (**Figure 6E,L**). These findings establish the epicardium as a major source of intracardiac fibroblast and mural-cell diversity and reveal spatially restricted developmental trajectories within its mesenchymal derivatives. By contrast, valve mesenchyme (cl.20) showed no dominant clonal association, instead exhibiting weak coupling to both epicardium-related populations and cl.19 great-artery VSMCs, consistent with contributions from mesodermal and neural crest lineages (**Figure 6E**). We therefore next examined the clonal architecture of valve mesenchyme at higher resolution.

### Epicardium contributes multiple valve-mesenchymal states that converge with neural crest-derived populations

Valve mesenchyme is assembled from endocardial, epicardial and neural crest-derived progenitors, but their relative contributions remain contested, in part because Cre drivers report marker expression rather than ancestry. Marker-independent clonal tracing is therefore well suited to this problem. Consistent with a multi-lineage origin, cl.20 valve mesenchyme showed dual clonal coupling to neural crest- and mesoderm-related populations (GA VSMC and epicardium-related cells, respectively; **Figure 6E**). Exocoelomic cavity injection only sparsely targeted endocardial cells, and these showed no significant linkage to valve mesenchyme (**Figure S16**, **Table S11, S15**). Thus, the endothelial-to-mesenchymal (EndoMT)-competent endocardial progenitors that contribute to valve mesenchyme were likely not targeted, consistent with E9.5 and E10.5 data (**Figures 1 and 2, and Figure S1B**). We therefore set out to resolve the neural crest- and epicardium-derived arms of valve mesenchyme at clonal resolution.

To enrich for neural crest-linked valve mesenchyme, we repeated the in utero lentiviral injection labelling approach, using amniotic cavity injections at E7.5 to target the neural plate, a, and collected 6 targeted hearts for spatially aware annotation and clonal analyses (**Figure S17,S18, Tables S16, S17**). There were >5-fold more neural-crest related cells in the amniotic cavity-targeted hearts compared to the exocoelomic cavity-targeted hearts, as well as a population corresponding to neurons/glia (cl. 22, **Figure S17A-C**), confirming that this route enriches for neural crest labelling. As in the exocoelomic dataset, the population annotated as valve mesenchyme (cl.16), mapped specifically to the valve region (**Figure S17,S18**), and exhibited dual clonal lineage linkage to both mesodermal and neural crest lineages (**Figure S19**).

Integrating the amniotic and exocoelomic datasets and subsetting valve mesenchyme for lineage-informed subclustering resolved 16 subclusters, designated V1-V16 (**Figures 7A-C; Table S18**): four linked to neural crest derivatives, nine to mesodermal derivatives, and three proliferative populations without dominant lineage association (**Figures 7D,F,G and S20, S21**). Integration preserved the UMAP architecture observed in each dataset independently (**Figures 5B, 7A and S17A**).

**Figure 7.**
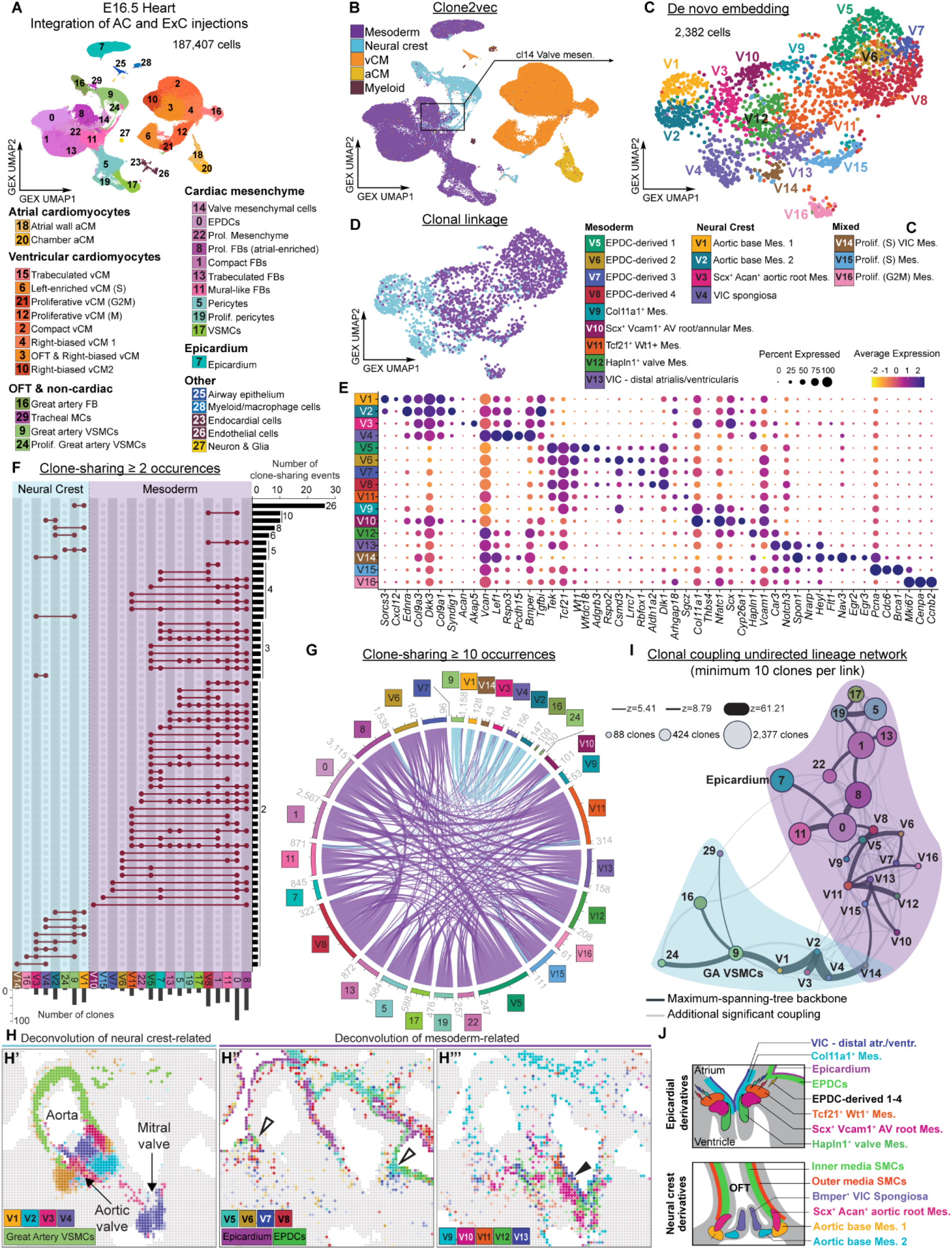
Integrated clonal and spatial analysis resolves convergent development of cardiac valve mesenchyme. **(A)** UMAP representation of the integrated E16.5 heart scRNA-seq dataset, comprising 187,407 cells from 12 hearts following E7.5 amniotic cavity (n=6) or exocoelomic cavity (n=4) targeting, and non-injected controls (n=2), colored by annotated population. **(B)** Clone2vec-inferred mesodermal (epicardium-related, purple) and neural crest (blue) clonal compartments projected onto the integrated gene-expression UMAP. Lineages related to mesoderm/ventricular cardiomyocytes, or mesoderm/atrial cardiomyocytes, are indicated in orange and yellow, respectively. The parental valve mesenchyme population (cl.14) selected for further analysis is indicated. **(C)** De novo embedding and sub-clustering of valve mesenchymal cells. **(D)** Mesodermal or neural crest clonal linkage of cells within cl.14. **(E)** Expression of markers used to annotate valve mesenchymal subclusters in (C). Dot size indicates the proportion of cells expressing each gene and color indicates scaled average expression. **(F)** UpSet plot showing clone sharing between valve mesenchymal subclusters and mesodermal or neural crest-related reference populations; combinations represented by at least two shared clones are shown, with bars indicating the number of shared clones. **(G)** Circos representation of clone sharing among valve mesenchymal subclusters and associated cardiac populations, showing relationships supported by at least ten shared clones. Link width indicates the extent of clone sharing and link color denotes mesodermal or neural crest association. **(H)** Spatial deconvolution of the integrated valve mesenchymal subclusters onto E16.5 heart Visium HD data at 16-µm resolution. **(H’)** Neural crest-linked aortic-base, aortic-root and spongiosa-like populations (V1–V4), together with great-artery VSMCs. **(H’’)** Epicardium, EPDCs and EPDC-derived intermediates V5–V8, showing their spatial progression from the epicardial surface towards the developing valves. White arrowheads show regions of putative inward migration. **(H’’’)** More differentiated epicardium/mesoderm-linked valve mesenchymal states V9–V13. Black arrowhead denotes flanking valve mesenchyme, localized in atrial region. **(I)** Undirected clonal-coupling lineage network connecting valve mesenchymal subclusters to epicardial, intracardiac mesenchymal and neural crest-related populations. Nodes represent transcriptomic populations and are scaled by cell number; edge width represents the clonal-coupling Z-score. Only positive associations supported by at least ten clones and significant after embryo-aware permutation testing and Benjamini–Hochberg correction (Z > 0; FDR ≤ 0.05) are shown. Black edges denote the maximum-spanning-tree backbone and grey edges denote additional significant associations; shaded regions indicate mesodermal and neural crest-related compartments, anchored by the Epicardium (cl.7) and great artery VSMCs (cl.9), respectively. **(J)** Model summarizing the spatially organized contributions of epicardial/mesodermal and neural crest lineages to cardiac valve and outflow-tract mesenchyme. AC, amniotic cavity; AV, atrioventricular; ExC, exocoelomic cavity; EPDC, epicardium-derived cell; FB, fibroblast; GA, great artery; Mes., mesenchyme; OFT, outflow tract; VIC, valve interstitial cell; VSMC, vascular smooth muscle cell. See also Figure S17-S22.

The four neural crest-linked populations were predominantly localized in the outflow tract (**Figure 7H, S20**) and comprised two aortic base-enriched mesenchyme populations, one including outer media smooth muscle cells (*Sorcs3*^+^ *Cxcl12*^+^ V1), and one restricted to the base, adjacent to the V1 cells (*Ednra*^+^ *Col9a3*^+^ V2 clls). Two further populations included n aortic root chondrogenic mesenchyme (Acan+ Akap5+, V3) and a spongiosa-like ventricular interstitial population (Vcan+ Rspo3+ Bmper+, V4). Complete marker sets for all subclusters are given in Tables S19 and S20.

Of the mesoderm-derived subclusters, four Wt1+ Tcf21+ populations (V5-V8) were strongly coupled to cl.0 EPDCs and are hereafter termed EPDC-derived cells (**Figure 7C-G, S20, Table S19, S20**). These localized along a spatial trajectory extending from the epicardial surface towards the developing valves, consistent with inward migration (**Figure 7H’’**) and were distinguished by *Wfdc18*, *Adgrb3* (V5); *Rspo2*, *Csmd3*, *Lrrc7* (V6); *Rbfox1* (V7); or *Aldh1a2* and *Dlk1* (V8). The EPDC-derived cells were, in turn, strongly coupled to five more differentiated clusters. These comprised two *Col11a1*⁺ *Nfatc1*⁺ populations, including *Thbs4*⁺ mesenchyme (V9) and atrioventricular valve-root/annular mesenchyme (*Scx*⁺, V10); *Tcf21*⁺ *Wt1*⁺ *Arhgap18*⁺ *Sgcz*⁺ mesenchyme (V11); mitral valve core/base mesenchyme (*Hapln1^+^*, V12); and atrialis/ventricularis-like VICs localized distally within the mitral valve on the atrial side (*Car3*⁺ *Notch3*⁺, V13) (**Figure 7H’’’**).

Together, clonal relationships, spatial distribution and transcriptional profiles supported a differentiation trajectory from epicardium through EPDC intermediates to valve mesenchyme. To represent these relationships without imposing a hierarchy, we adapted the clonal-coupling framework of Bandler et al.^36^ to build an undirected network in which clusters are connected by significant positive clonal coupling.

Because clonal coupling is symmetric and therefore carries no inherent direction, we oriented the network using the spatial gradient and the degree of transcriptional maturation rather than inferring direction from the lineage data themselves. The resulting continuum linked epicardium to EPDCs, EPDC-derived intermediates and Tcf21+ Wt1+ mesenchyme, and thereafter to more differentiated Hapln1+ spongiosa-like and distal atrialis/ventricularis-like VIC states (**Figure 7I, Table S19, S21**). Valve mesenchyme is therefore assembled by convergent development, in which transcriptionally similar states arise from mesodermal/epicardial and from neural crest progenitors that occupy distinct anatomical territories (**Figure 7J**). Critically, analysing the amniotic and exocoelomic datasets separately reproduced this architecture both transcriptionally and clonally (**Figure S22**), indicating that the convergent organisation is not a product of integration; the separate analyses also resolved the outer medial smooth muscle cells as neural crest-linked. The assignment of outer medial smooth muscle cells to a neural crest lineage was unexpected, as these cells have previously been attributed to the anterior/second heart field (see Discussion).

Three proliferative subclusters (V14-V16) showed no dominant lineage association, which we ascribe to convergence on a proliferative transcriptional state that masks cell-type signatures. V14 nonetheless displayed a specialized Egr2+ Egr3+ programme and localized to atrialis/ventricularis-like regions of both valve types (**Figure S20**); its mixed clonal associations suggest that V14 comprises transcriptionally convergent populations of distinct developmental origin at the two locations, which cannot be separated at the current resolution despite the high read depth.

These data directly establish shared clonal ancestry among epicardium, EPDCs and defined valve-mesenchymal states. Their ordered spatial distribution and progressive transcriptional differentiation further support an epicardium-to-valve developmental route, and suggest epicardium contributes more valve mesenchyme populations than previously reported.

## DISCUSSION

In utero lentiviral delivery provides a rapid and scalable complement to transgenic approaches for manipulating embryonic tissues ^2,3,12,14^. Whereas Cre-based lineage tracing depends on the specificity and timing of predefined molecular drivers, heritable barcode delivery labels progenitors independently of marker identity and enables clonal ancestry to be recovered alongside transcriptional state ^4,22,23^. Building on previous targeting of ectoderm and its derivatives, here we establish the exocoelomic cavity as an access route to mesodermal progenitors during gastrulation. By combining this approach with single-cell lineage tracing and spatial transcriptomics, we reconstructed clonal relationships across more than 590,000 cells and 31,000 multicellular clones and resolved developmentally distinct populations that converge transcriptionally within organ mesenchyme. Although barcode sharing establishes common ancestry rather than direct parent-daughter relationships, its scale and independence from marker selection enable quantitative comparison of lineage coupling across entire tissues. Exocoelomic and amniotic cavity injections therefore provide complementary access to embryonic compartments and establish a versatile platform for reconstructing and ultimately perturbing^37^ mammalian developmental lineages *in vivo*.

Genetic fate mapping has established a model in which septum transversum-derived mesothelium contributes to hepatic stellate cells, fibroblasts and perivascular mesenchymal populations during liver development ^8,9,38^, but the relationships among these derivatives and the extent of mesothelial heterogeneity have remained unresolved^39^. Our clonal analyses (**Figure 4**) strongly linked capsular mesothelium to hepatic stellate cells, providing marker-independent support for their shared developmental ancestry. However, mesothelial sub-populations exhibited distinct spatial and clonal biases: capsular M0 mesothelial cells were preferentially coupled to stellate-cell lineages, whereas portal-associated transitional M12 mesothelial cells were more strongly associated with fibroblast- and mural-cell-containing clones. Fibroblast and vascular smooth muscle populations also displayed more restricted coupling to mesothelium, indicating that liver mesenchyme does not arise through a single homogeneous mesothelium-to-mesenchyme trajectory. Moreover, the separation of pericyte-associated and stellate-cell-associated VSMC clonal architectures suggests that mural cells can be generated through distinct developmental routes. These findings extend the transcriptional heterogeneity identified in developing liver atlases ^40^ by linking mesenchymal states to spatially organized clonal histories. Together, they support a model in which fetal liver mesothelium comprises lineage-biased progenitor states that differentially contribute to stellate, fibroblast and mural-cell diversity, rather than functioning as a uniform multipotent source.

Our clonal analyses indicate that cardiac fate restriction is already evident by E7.5 (**Figure 6A-H**), but operates at different strengths across myocardial lineages. The pronounced separation of atrial and ventricular clones supports previous evidence that cardiac progenitors are spatially, temporally and molecularly patterned during gastrulation ^20,35,41^. This segregation was not absolute, as rare clones connected atrial cardiomyocytes to outflow tract-associated ventricular cardiomyocytes, consistent with residual developmental competence or a shared progenitor state at the time of labelling. Within the ventricular lineage, left- and right-enriched cardiomyocytes exhibited moderate clonal biases but remained embedded within a broader continuum of ventricular competence, extending earlier clonal models of myocardial regionalization^42^. These findings are also consistent with live-imaging evidence that atrial/inflow and left ventricular-atrioventricular canal progenitors emerge as distinct populations during gastrulation ^19^. Together, our data support a hierarchical model of cardiac specification in which atrial and ventricular identities segregate early, whereas regional ventricular fates remain comparatively plastic and become progressively biased rather than being separated by an absolute left-right lineage boundary.

The epicardium is an established source of cardiac fibroblasts and coronary mural cells, although the extent of its developmental potential and the organization of its derivatives remain debated ^10,11,43–45^. Our clonal analyses linked epicardium to fibroblast, pericyte and vascular smooth muscle populations and resolved three spatially organized trajectories (**Figure 6A,B,E,I-L**): an atrial EPDC-mural-like fibroblast lineage, a ventricular EPDC-proliferating fibroblast lineage, and a ventricular intramyocardial fibroblast-mural lineage spanning compact and trabecular regions. The coupling of pericytes to mature coronary VSMCs is consistent with evidence that pericytes act as intermediates in coronary arterial smooth muscle differentiation ^46^, while the segregation of these trajectories indicates that epicardial mesenchymal diversification is regionally patterned rather than generated through a single broadly multipotent state. By contrast, epicardium showed no substantial clonal coupling to cardiomyocytes. Earlier *Tbx18*- and *Wt1*-based fate-mapping studies suggested a proepicardial contribution to myocardium ^47,48^, but interpretation of these experiments has been complicated by expression of the respective drivers outside the epicardium ^49^. Our marker-independent clonal data therefore support a model in which early proepicardial progenitors generate spatially restricted fibroblast and mural-cell lineages but make little or no direct contribution to the cardiomyocyte compartment during normal development.

Cardiac valve mesenchyme is assembled from several embryonic sources, including endocardial, epicardial and neural crest-derived progenitors whose relative contributions vary between valve regions^34,50^. Endocardium was not meaningfully labeled with the injection routes used here, precluding comprehensive reconstruction of the endocardial-linked contribution. In contrast, epicardium and neural crest were well-labeled, leading to high resolution single cell lineage tracing of their linked populations. We identified strong clonal coupling between epicardium and multiple valve-mesenchymal populations and, unexpectedly, between neural crest derivatives and outer medial SMCs of the ascending aorta. This contrasts with Sawada et al., who assigned outer medial SMCs to the anterior/second heart field using *Mef2c*-AHF-Cre and found *Wnt1*-Cre2 labelling confined to the inner media ^51^. However, the specificity and completeness of *Wnt1*-Cre2 labelling remain uncertain: Gandhi et al. reported ectopic recombination in non-neural-crest tissues in a preprint, whereas its coverage of neural crest derivatives has not been established^52^. Although outer medial SMCs express endogenous *Mef2c*, whether the specific AHF enhancer used in this driver remains active in differentiated SMCs is unknown. In our data, these cells were strongly coupled to neural crest lineages but showed no detectable coupling to sampled anterior/second heart-field populations, including outflow-tract cardiomyocytes. Together, these findings support a substantial neural crest contribution to outer medial SMCs of the embryonic ascending aorta, challenging a strict laminar model in which neural crest and second heart-field derivatives are confined to the inner and outer media, respectively.

Although *Tie2*-Cre fate mapping supports an endocardial contribution to valve mesenchyme ^53–56^, the detection of *Tek* (Tie2) in epicardial-related mesenchymal populations (our data and Feng et al^31^) raises the possibility that the cellular specificity of individual *Tie2*-Cre models should be considered when interpreting these experiments. *Tie2*-Cre exhibits variable cell labelling efficiency^56^ and appears to drive sporadic labelling of cells in the epicardial region^53,57^. Comparative *Tie2*-Cre and *Wt1*-Cre fate mapping revealed leaflet-specific contributions of endocardial- and epicardial-derived mesenchyme to the atrioventricular valves^50^. This was corroborated by endothelial *Cdh5-CreER-*tracing^58^, as well as an elegant dual *Cdh5*-Nigri-nox and *Tbx18*-Cre-loxP recombination system^59^, similarly demonstrating unequal endothelial contributions among valve leaflets. The exocoelomic and amniotic cavity injection routes employed here did not appreciably label endocardium (**Figure S1B**, **Figure S16,S18**), and the convergent-development model presented in Figure 7 therefore focuses on the epicardial and neural-crest arms of this multi-lineage system. Together, our findings resolve the epicardial and neural crest arms of convergent valve development and provide a framework for incorporating the endocardial contribution into a complete lineage-resolved model of leaflet formation. Future work to identify in utero targeting approaches able to access endocardium would provide a Cre-independent approach to map endocardial derivatives at high resolution, and efforts towards 3D spatial barcode lineage tracing would also transform our understanding of the development of this complex structure.

By integrating cell state, clonal ancestry and anatomical location at scale, our approach reveals lineage restriction as a progressive and context-dependent process, encompassing both early fate segregation and continued diversification within developing organs. Extending this framework across developmental stages, genetic perturbations and models of injury or disease should enable direct investigation of how lineage history constrains cellular plasticity and shapes tissue function. More broadly, adaptable in utero delivery routes provide a versatile platform for reconstructing and experimentally manipulating mammalian development without dependence on predefined lineage markers.

### Limitations of the study

Several features of the approach define the scope of its interpretation. Barcode sharing identifies common ancestry among cells labeled around E7.5, but does not by itself establish a direct parent-daughter relationship between populations recovered at E16.5. Accordingly, the resolved architectures are most appropriately interpreted as clonal relations rather than linear differentiation paths. Clonal coupling is also influenced by population abundance and cell recovery, potentially limiting detection of relationships involving rare or inefficiently captured cell types. Nevertheless, the reproducibility across biological replicates, agreement among complementary clonal analyses, and concordance with established lineage relationships support the robustness of the major conclusions. Integration with spatial transcriptomics further provided the spatial context required to distinguish transcriptionally related populations. Although all terminal populations were sampled at E16.5, the coexistence of epicardial, EPDC-like, intermediate and differentiated states enabled reconstruction of a spatially and transcriptionally ordered developmental continuum. Finally, because lentiviral integration occurs over a finite interval following injection, E7.5 should be viewed as the developmental window during which progenitors were labeled rather than an instantaneous fate-mapping time point. These considerations do not alter the principal lineage relationships identified here, but define the resolution at which this framework reconstructs mesodermal development.

## RESOURCE AVAILABILITY

### Sequencing data, biological samples and barcodes

All processed sequence files (e.g. BAM alignment files) and derived data (e.g. normalised read count matrices) for the E9.5/E10.5 whole embryos, E16.5 heart, and E16.5 liver scRNA-sequencing, and corresponding VisiumHD, are deposited in Annotare and loaded to the ArrayExpress database, and will be publicly available as of the date of publication (accession numbers listed in Resources table).

All original code has been deposited at Github, and will be publicly available, as of the date of publication.

## ACKNOWLEDGMENTS

We thank our colleagues Sarantis Giatrellis, Paulina Zydowicz-Machtel, Becky Wagner and Linda Roessler for helping with experiments, and we thank Claudio Cortes Rodriguez, Leonie Von Berlin and Stefaan Verhulst for scientific discussions. We acknowledge the support from Isabel Espinosa-Medina for this project. We acknowledge the Biomedicum Imaging Core Facility (BIC) at Karolinska Institutet, financed by the Infrastructure Board, for providing advanced microscopy services and instrumental expertise. We acknowledge the Histological Core Facility (Histocore) and Biomedicum Flow Cytometry Core Facility (BFC) at Karolinska Institutet for access to histology and flow cytometry instrumentation. We acknowledge the National Genomics Infrastructure in Stockholm, funded by Science for Life Laboratory, the Knut and Alice Wallenberg Foundation and the Swedish Research Council, for sequencing services; National Bioinformatics Infrastructure Sweden (NBIS) for bioinformatics assistance (WABI), SNIC/Uppsala Multidisciplinary Centre and PDC, KTH Royal Institute of Technology for Advanced Computational Science for assistance with massively parallel sequencing and access to the UPPMAX and Dardel computational infrastructure. We acknowledge Mattias Karlen and BioRender for help with figure schematics.

This work was funded by the European Union under the Horizon Europe grants (divENSify; 101045026 and LIMITLESS; 101171156). As set out in the Grant Agreement, beneficiaries must ensure that at the latest at the time of publication, open access is provided via a trusted repository to the published version or the final peer-reviewed manuscript accepted for publication under the latest available version of the Creative Commons Attribution International Public License (CC BY) or a license with equivalent rights.

Knut and Alice Wallenberg project grant (ERA, JF, IA; 2020.0109)

Karolinska Institutet Forskarassistent, Förlängning, Consolidator grants (ERA; 2-560/2015-280, 2-2110/2019-7, 2-195/2021)

Karolinska Institutet StratRegen Senior Grant (ERA)

Swedish Research Council MH 3R Project Grant (ERA, 2017-01054, 2021-03537 and 2024-03113)

Swedish Research Council MH Project Grant (ERA, 2019-01350)

Wallenberg Bioinformatics Support (ERA, LB)

ERC LIMITLESS (ERA, 101171156)

ERC divENSify (UM, 101045026)

Hjärnfonden (ERA, FO2025-0271)

Göran Gustafsson prize in Medicine (ERA, GG2026-0003)

Leo Foundation (ERA, LF-OC-25-001992)

N.C.H. was supported by a Wellcome Trust Senior Research Fellowship in Clinical Science (ref. 219542/Z/19/Z)

## AUTHOR CONTRIBUTIONS

Conceptualization: JH, ERA

Methodology: JH, JLo, AAC, SI, RM, SdH, LB, JS, NVH, MR, EL, ERA

Investigation: JH, JLo, AAC, SI, RM, SdH, LB, JS, BS, NVH, RT, JW, MM, EL

Visualization: JH, JLo, AAC, SI, RM, SdH, LB, EL

Funding acquisition: JLu, ERA

Project administration: ERA

Supervision: ERA, JF, IA, JLu

Writing - original draft: JH, ERA

Writing - review and editing: JH, JLo, AAC, SI, RM, SdH, LB, JS, BS, NVH, RT, JW, UM, MM, NCH, JF, MR, PK, IA, EL, JLu, ERA

## DECLARATION OF GENERATIVE AI AND AI-ASSISTED TECHNOLOGIES IN THE WRITING PROCESS

During the preparation of this work, the author(s) used ChatGPT (OpenAI), Claude (Anthropic) and Codon (a Karolinska Institutet-internal AI co-scientist developed by Sten Linnarsson and Roni Henareh) in order to refine language of the manuscript text, and as coding assistance. After using any tool or service, the author(s) reviewed and edited the content as needed and take(s) full responsibility for the content of the publication.

## SUPPLEMENTAL INFORMATION

Supplementary Figures S1–S22.

## STAR★METHODS

### KEY RESOURCES TABLE

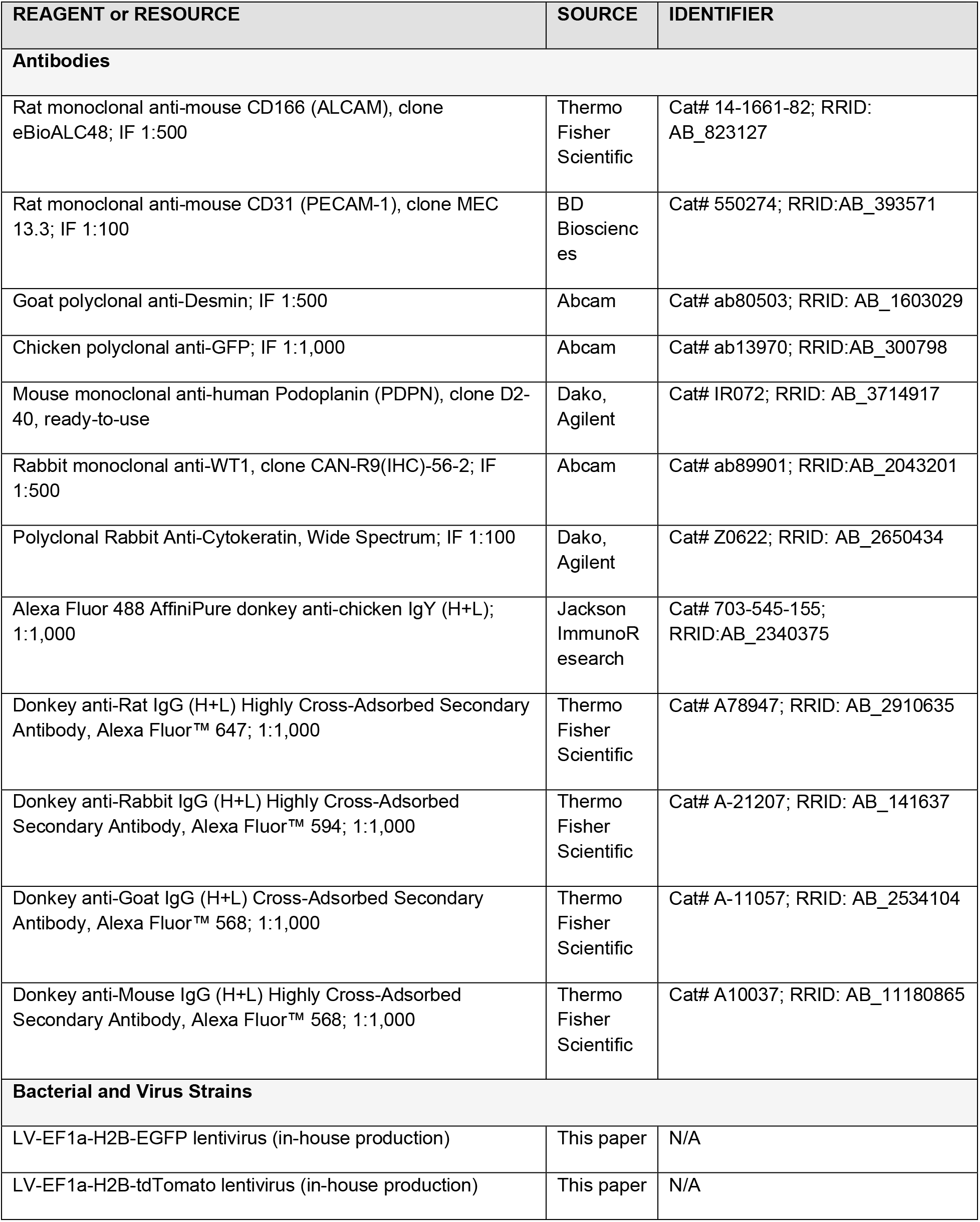
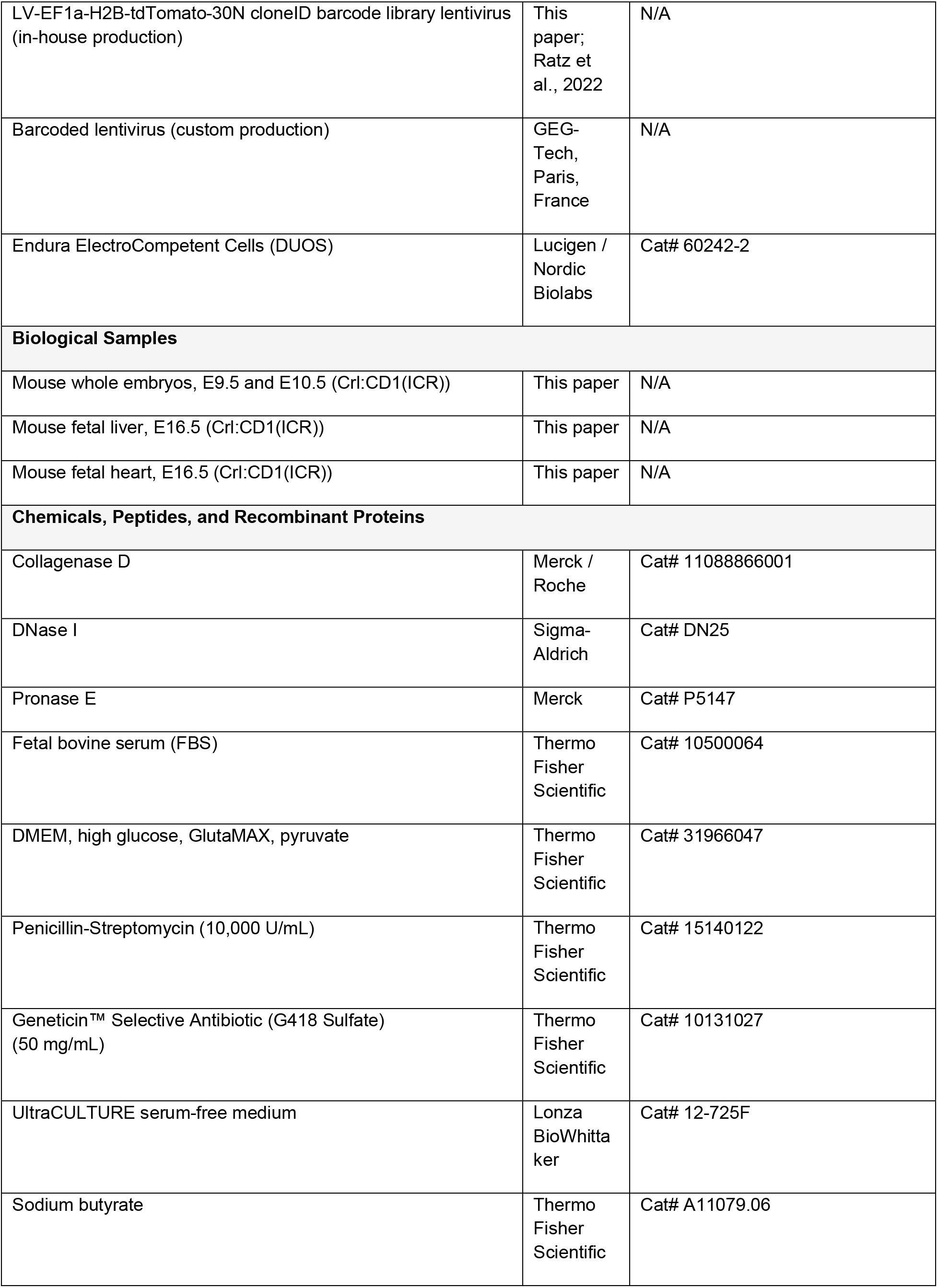
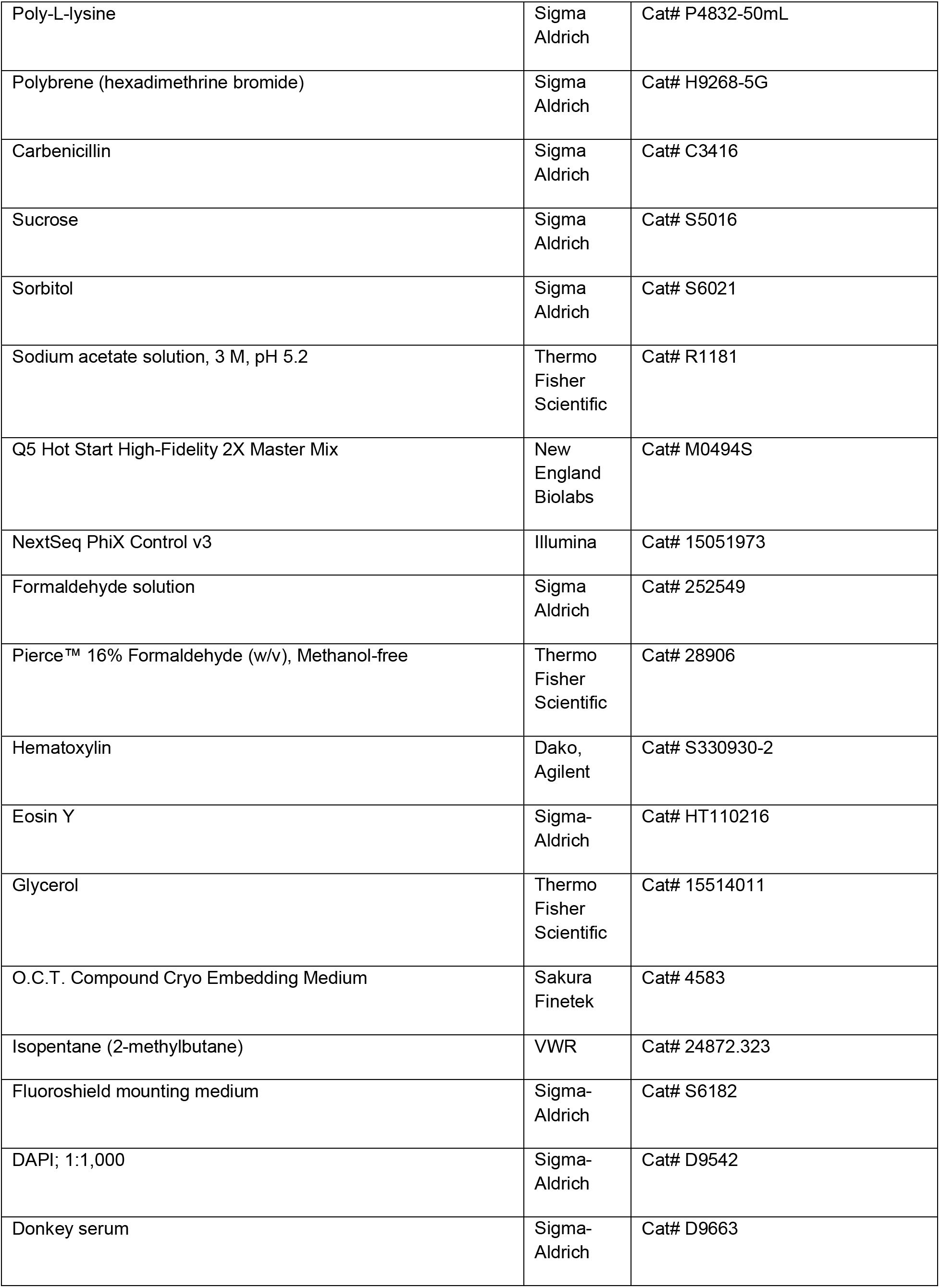
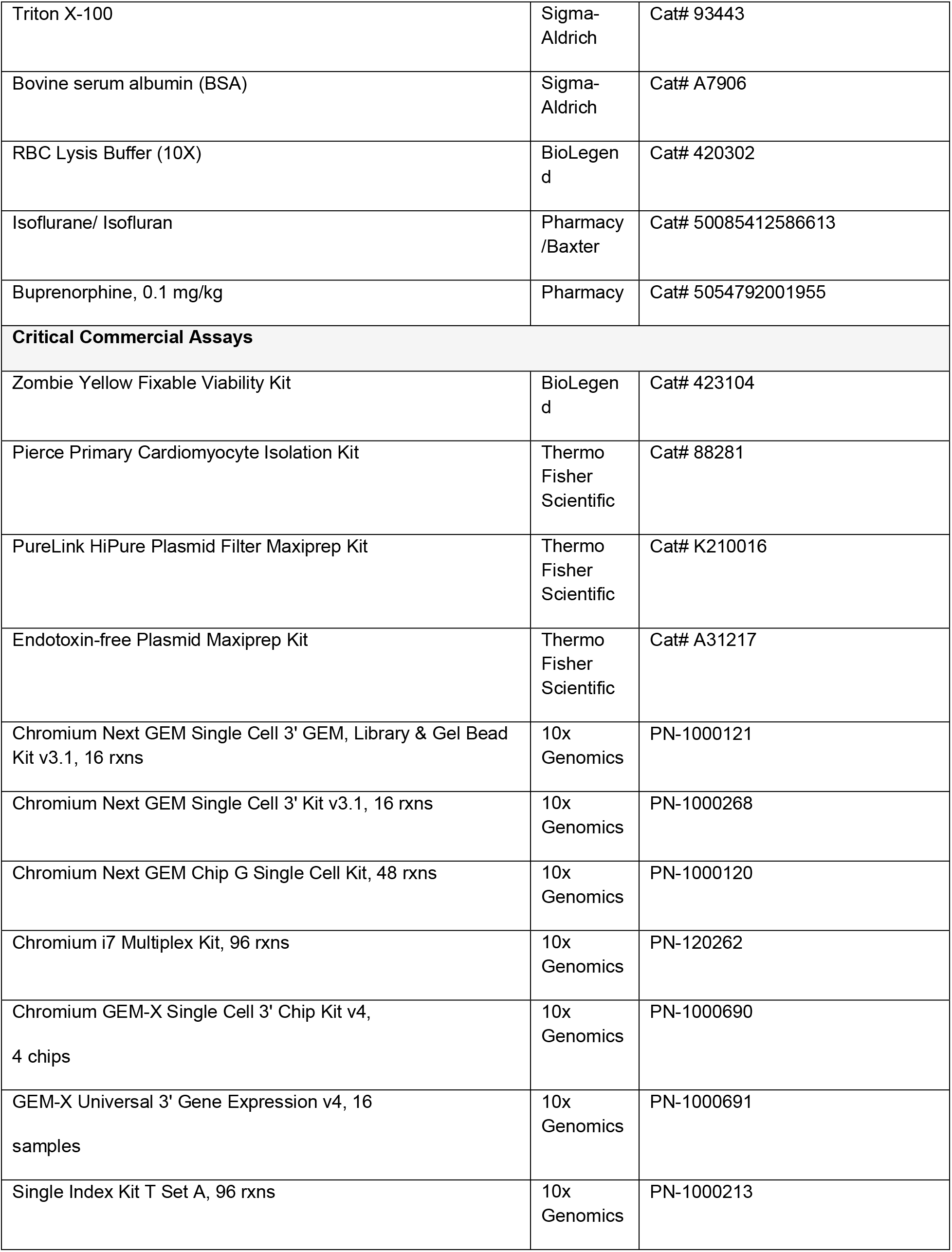
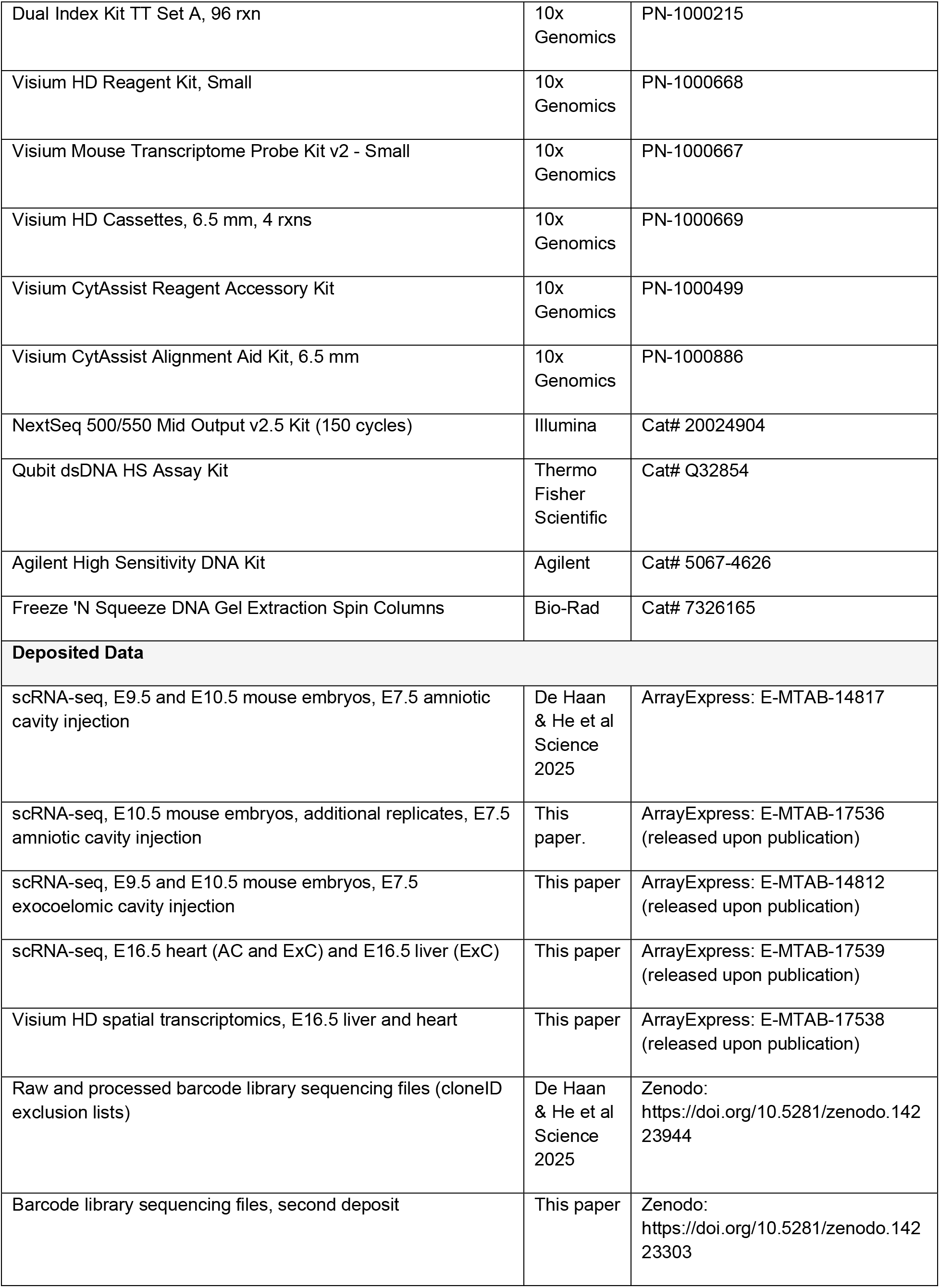
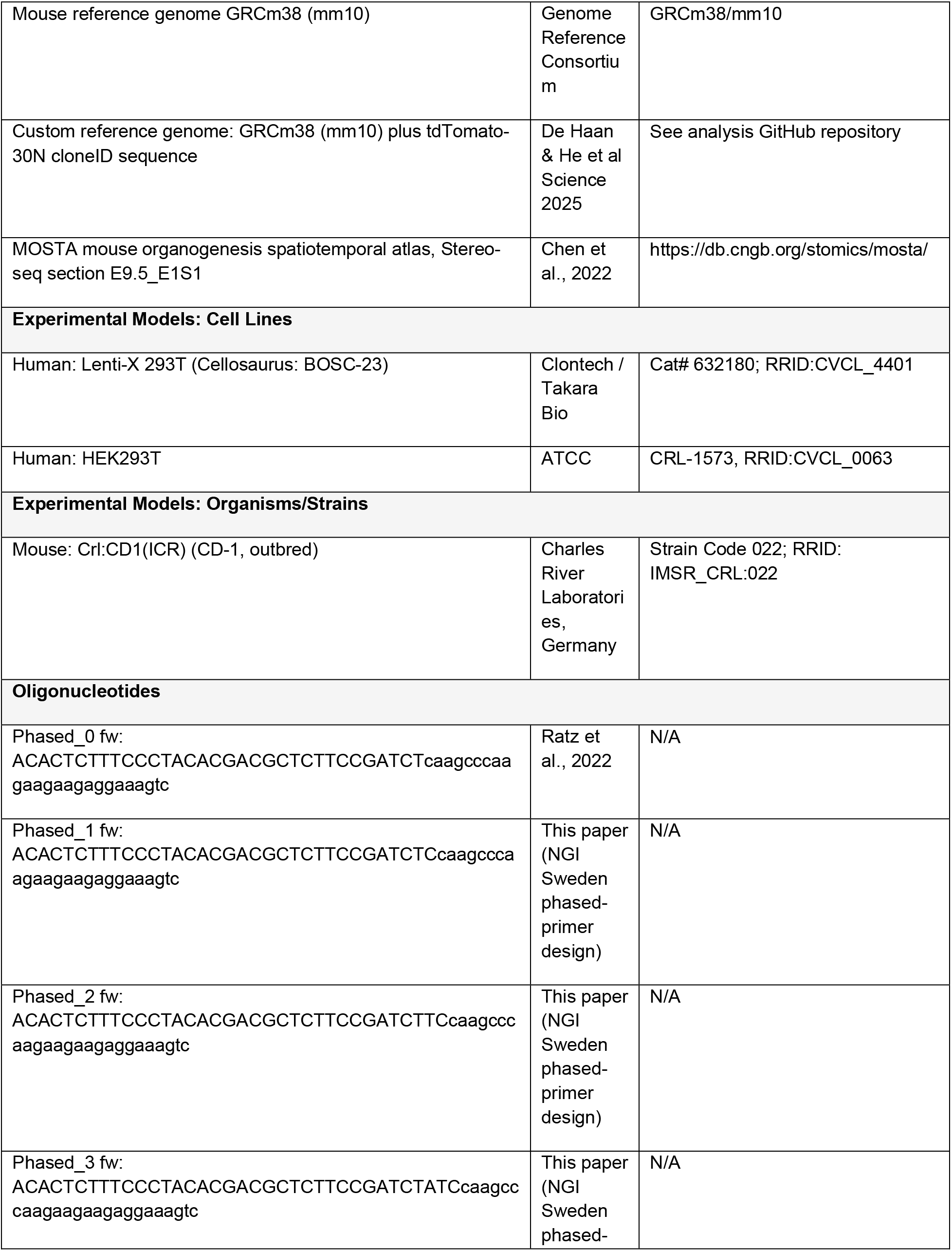
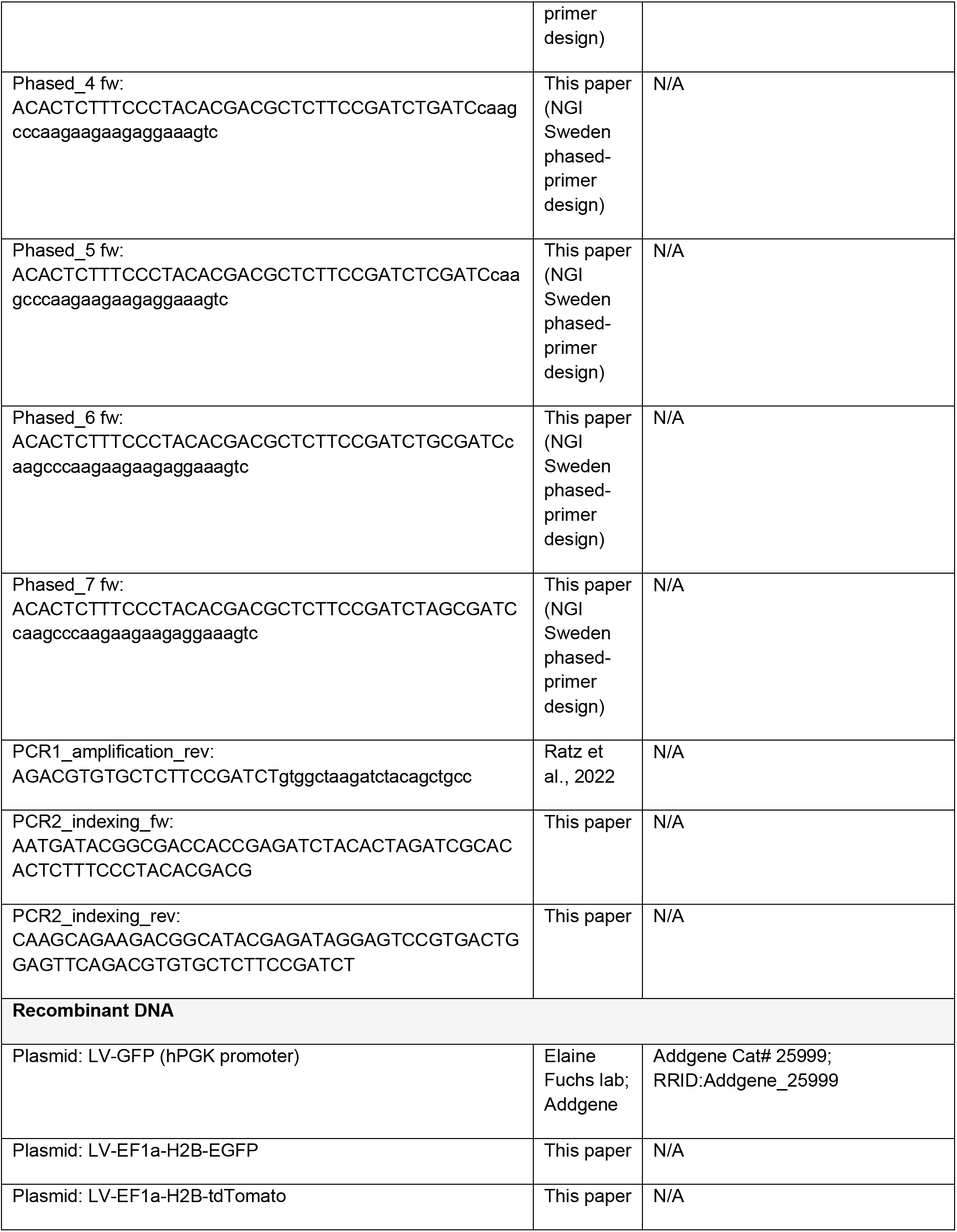
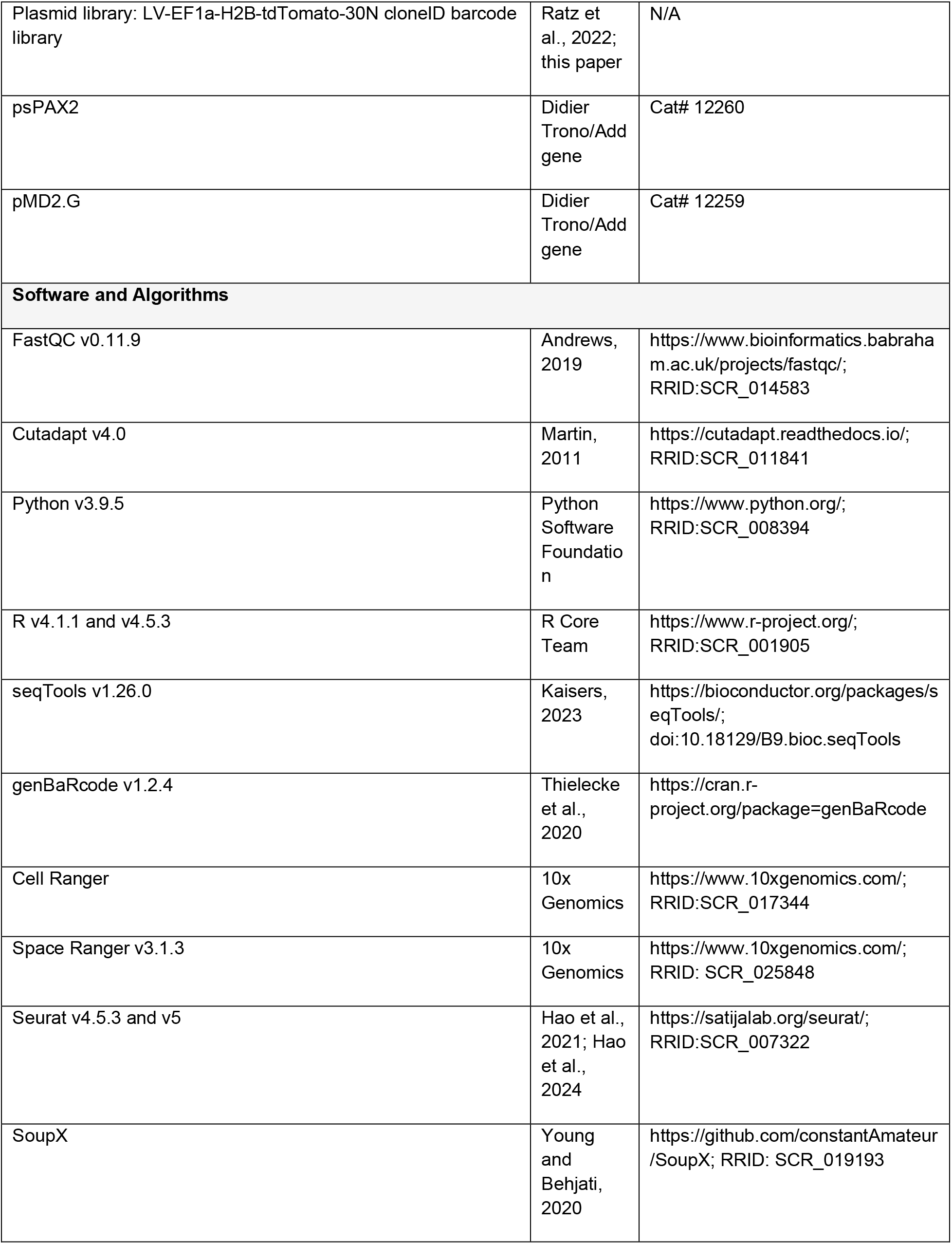
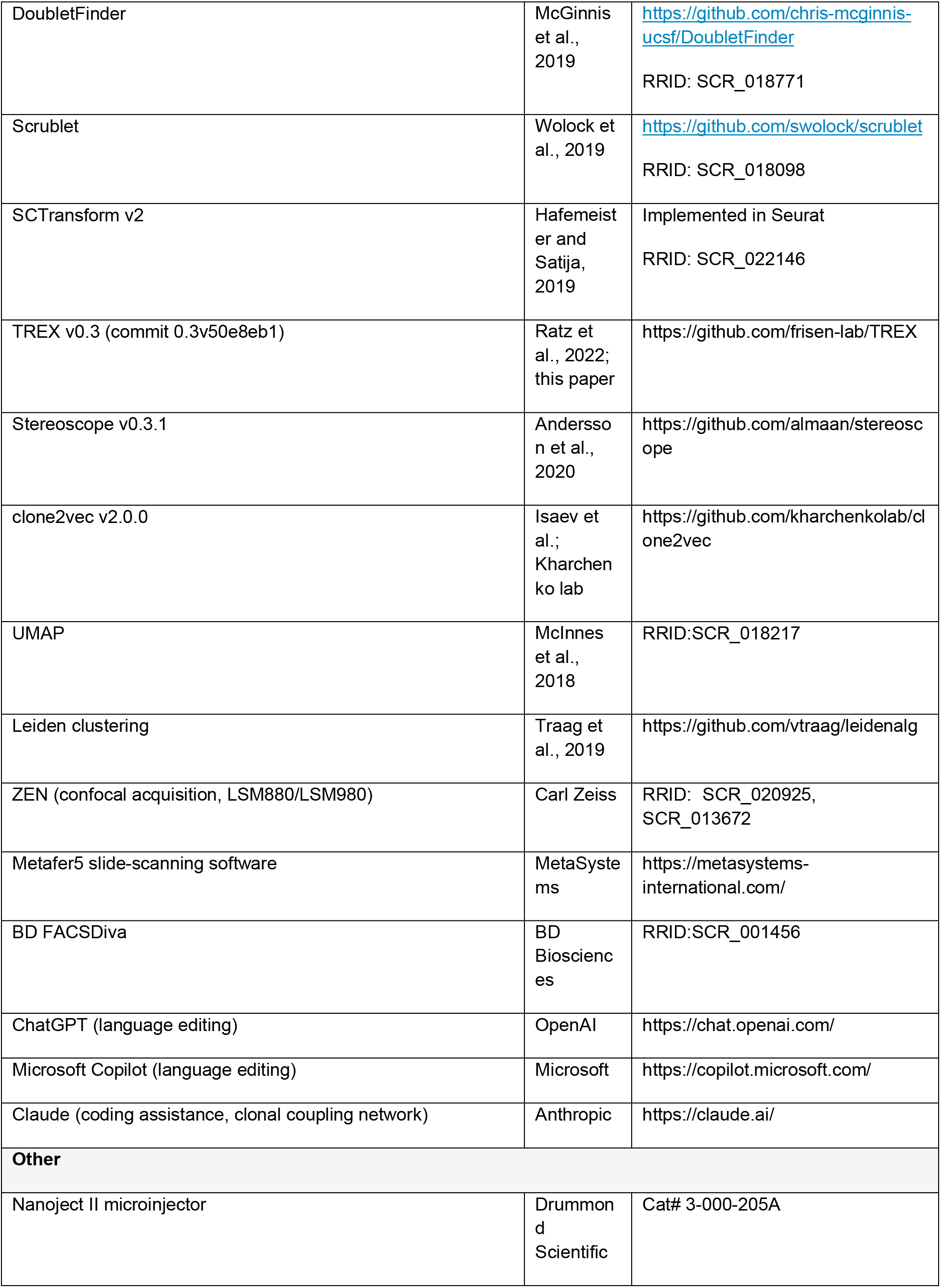
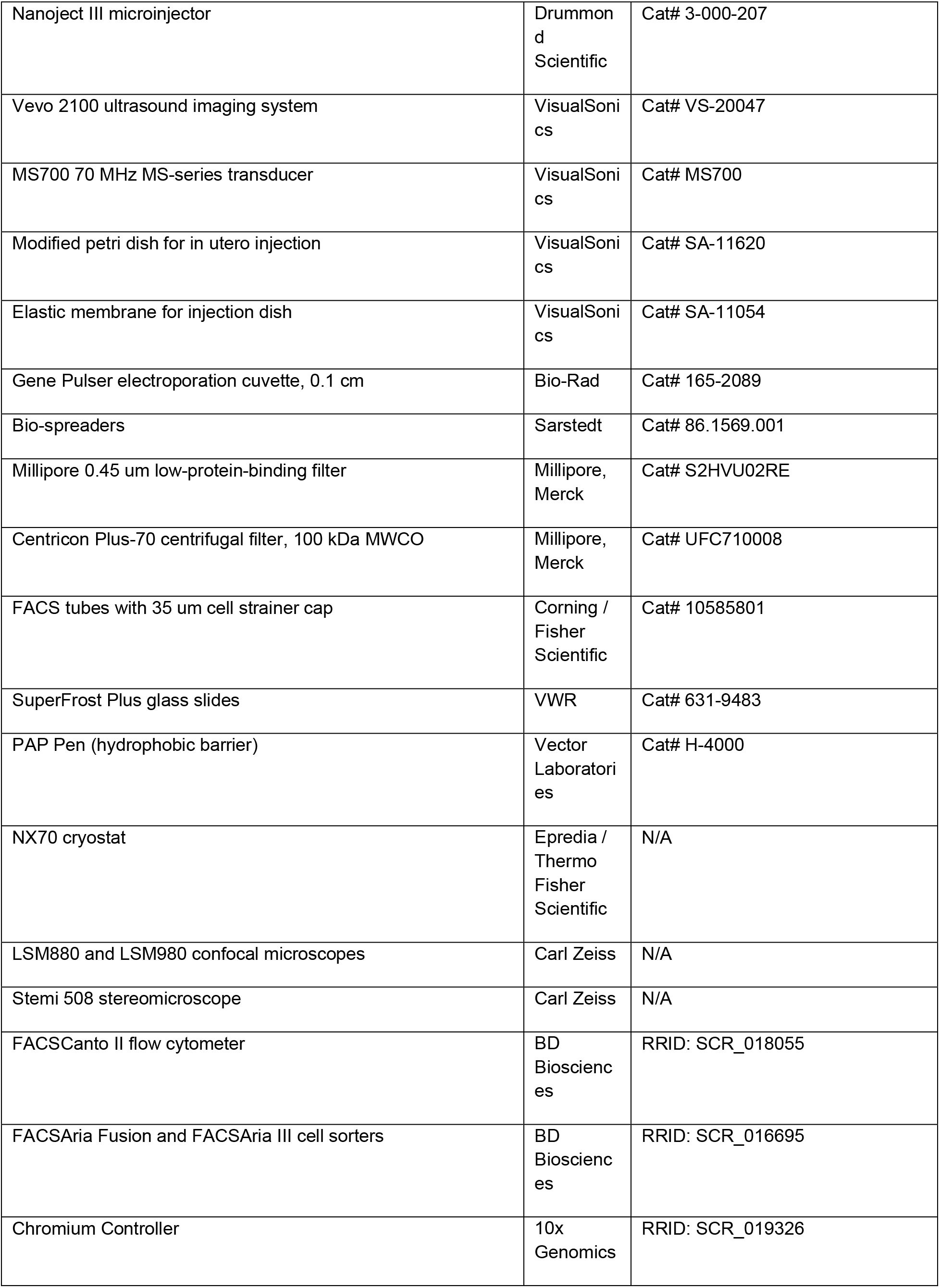
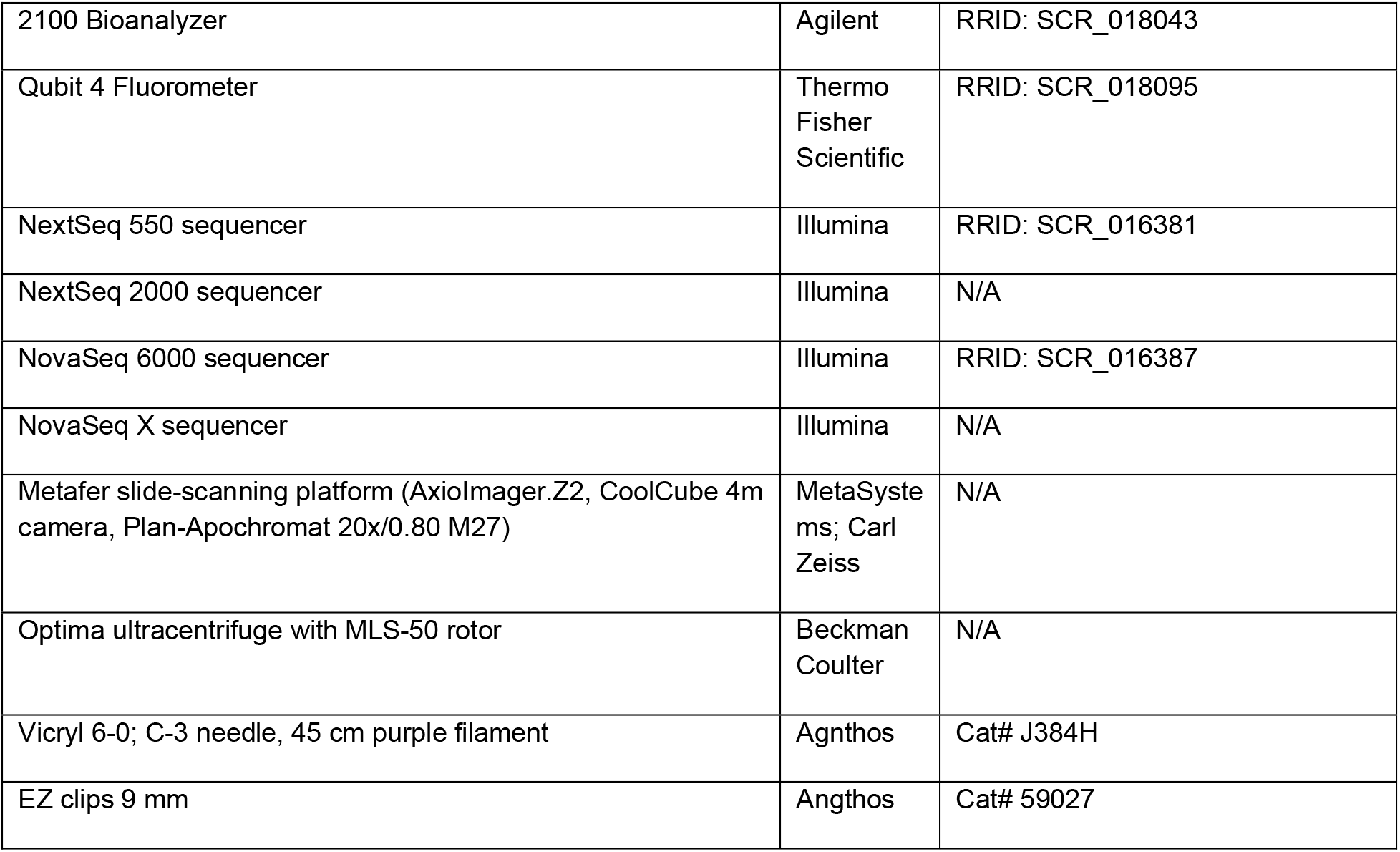

### EXPERIMENTAL MODEL AND STUDY PARTICIPANT DETAILS

#### Mice

CD-1 mice (Crl:CD1(ICR)) were purchased from Charles River Laboratories (Germany). Mice were housed with standard light cycles and following European regulations. Females (8-20 weeks old) were used for estrus check and mated with males overnight. Females were checked for vaginal plugs the following morning, with noon on the day of plug detection designated embryonic day 0.5 (E0.5). While in utero injections were performed at circa E7.5, embryos were staged for *in utero* injection using Theiler staging based on ultrasound imaging. The sex of embryos was not included in analyses. All experiments were performed according to the ethical approval granted by the Swedish Board of Agriculture (Jordbruksverket, N59/14, 8188-2017, 2987-2020, 5237-23; 1613-24).

#### Cells and culture conditions

Lenti-XTM 293T cells (Clontech) were used for lentivirus production and cultured at 37°C/ 5% CO2 in DMEM complete medium (DMEM, high glucose, GlutaMAX, pyruvate, supplemented with 10% fetal bovine serum (FBS), 1% penicillin/streptomycin and 1% geneticin). HEK293T cell line was used for virus titration and cultured at 37°C/ 5% CO2 in DMEM complete medium (DMEM, high glucose, GlutaMAX, pyruvate, supplemented with 10% fetal bovine serum (FBS), 1% penicillin/streptomycin).

### METHOD DETAILS

#### Plasmid construction

The plasmid construction was done as previously described ^60^. LV-EF1α-H2B-EGFP/tdTomato was constructed by replacing the hPGK promoter from LV-GFP (#25999, Elaine Fuchs lab, Addgene) with the EF1α promoter.

#### Plasmid library amplification

To amplify the barcode library, 100ng of plasmid DNA (pDNA) was added to 50uL of thawed electrocompetent cells (Endura ElectroCompetent Cells (DUOS), Nordic biolabs, Cat# 60242-2). Plasmid and cells were transferred into electroporation cuvette (Bio-rad, Cat#165-2089). Immediately after electroporation, 2x 750 uL recovery medium was added. The medium was transferred to a round-bottom 14 mL tube. After the recovery by shaking at 220 rpm at 37°C incubator for 1 h, all cells were combined and distributed onto four 500 cm2 LB agar plate with 50 ug/mL carbenicillin using biospreaders (Starsted, Cat #86.1569.001). To get an accurate colony count, diluted samples were prepared as follows: 10 uL of combined culture was mixed with 990 uL of LB medium, and a final dilution of 1:1,000,000 was achieved by sequential dilutions. The dilutions 1:10,000 (#1) and 1:1,000,000 (#2) were used for colony counting. All colonies in the 500 cm2 plates were collected and centrifuged at 3,000 g at 4℃ for 30min to pellet the cells, which were then used for plasmid maxiprep.

#### Barcode library sequencing

To analyze the barcode library after plasmid expansion, we performed DNA sequencing of the amplified barcode region (**Materials and Methods Fig**). We used a phased primer strategy for the sequencing library preparation to mitigate low complexity amplicon issues for sequencing libraries on the selected Illumina platform. Phased primers were designed using a tool developed by the NGI Sweden facility (**Key Resources table**) to amplify the 30N barcode and its flanking regions (107 bp). The sequencing library was prepared in two rounds of PCR with Q5 Hot Start High-Fidelity 2X Master Mix (NEB, M0494S) as previously described ^60^, with 20 ng of plasmid library as starting material, in addition to sequencing adaptors and indexes, which yielded a 243 – 250 bp product, due to the phased primers. The final PCR product was purified from 1.2% agarose gel using Freeze ’N Squeeze DNA gel extraction spin columns (Bio-Rad, cat. no. 7326165) and precipitated with 3M Sodium Acetate Solution, pH 5.2 (Thermo Fisher Scientific, cat. no. R1181) and isopropanol. The sequencing library was diluted to 2 nM, denatured, spiked with 20% NextSeq™ PhiX Control (Illumina, cat. no. 15051973) and sequenced on Illumina NextSeq550 instrument using NextSeq500/550 Mid-Output v2.5 Kit (150 cycles) (cat. no. 20024904). Read 1 was set to 100 nt, read 2 to 50 nt. Only read 1 was considered for further analysis.

Quality control of raw sequencing files (deposited at 10.5281/zenodo.14223944) was performed using FastQC ^61^ version 0.11.9. The following software packages were used to filter fastq reads and extract and process 30N barcodes: Cutadapt ^62^(version 4.0) with Python 3.9.5, seqTools ^63^ (version 1.26.0) and genBaRcode ^64^ (version1.2.4) in R 4.1.1.

#### Lentivirus production

All viruses were generated using plasmids prepared with Endotoxin-free Maxiprep kits (Thermo Fisher Scientific, #A31217) or PureLink™ HiPure Plasmid Filter Maxiprep Kit (ThermoFisher Scientific, #K210016). H2B-GFP lentivirus was produced in-house. Barcoded lentivirus was produced in-house or purchased from GEG-Tech (Paris, France). For the in-house lentivirus production, we followed the previously described protocol ^65^. In brief: LentiX-293T cells were seeded in one T-75 flask and split into four T-225 flasks when reaching ∼ 80% confluency. After culturing for ∼3 days, cells reached 80-90% confluence and were split 1:1 to four 500 cm^2^ plates coated with poly-L-lysine. Two plates of cells were used for one batch of virus production. The next day, cells were transfected with the calcium transfection method. 12-14 h after transfection, medium was replaced with fresh DMEM culture medium (DMEM + 10% v/v FBS + 1% v/v Pen-strep/L-glut mix + 1% v/v 100mM Sodium Pyruvate + 1% v/v 7.5% sodium bicarbonate). 16 h after transfection, medium was replaced with viral production medium (VPM), (Ultraculture (Lonza BioWhittaker 12-725F) supplemented with 1% v/v Pen-strep/L-glut mix + 1% v/v 100mM Sodium Pyruvate + 1% v/v 7.5% sodium bicarbonate and 5 mM sodium butyrate). To obtain high titer virus, supernatants were collected at both 46 hours and ∼65 hours after transfection, filtered through a 0.45 mM Millipore low-protein binding filter (#S2HVU02RE, Millipore, Merck), and concentrated with a two-step centrifugation method, after first-step centrifugation at 3,300 g through 100 kDa MW cutoff Millipore Centricon 70 Plus cartridges, yielding 4mL of virus supernatant. Next, a 20% sucrose gradient ultracentrifugation was performed at 45,000 rpm (MLS 50 Rotor) for 1 h 30 min. The final lentiviral particles were resuspended in 30-50 µL viral resuspension buffer (VRB) (20 mM Tris pH 8.0, 250 mM NaCl, 10 mM MgCl2 and 5% sorbitol) and aliquoted into vials (5 µL per vial), snap frozen on dry ice, and stored at -80°C.

#### Virus titration

The titers of virus encoding fluorophores produced in house were determined using HEK293T cells. 24 h prior to transduction, cells were seeded in a 6-well plate at a density of 800K cells/well in DMEM complete medium (DMEM containing 10% FBS and 1% pen/strep). On the day of transduction, 1 µL of virus was diluted in 2 mL Infection medium (DMEM +10% FBS+ 10 µg/mL polybrene) and 5, 50, and 500 µL of diluted virus were added to four individual wells. One well was used for cell counting to determine the cell number when transducing. One well was kept as a non-transduced control. Plates were then centrifuged for 30 min at 1100 g at 37°C. The infection medium was discarded and replaced with DMEM complete medium. After 48 h, cells were collected, fixed with 4% PFA for 15 min at room temperature and transferred to FACS tubes with 35 µm strainer (Corning, FisherScientific, #10585801). Quantification of fluorophore+ cells was performed using a Canto II flow cytometer (BD). The functional titer was then calculated in infectious units (IFU) per mL.

#### Ultrasound-guided *in utero* nano-injection

A detailed description of E7.5 amniotic cavity injection was published previously ^65,66^. In this study, all embryos were injected either into the amniotic cavity (AC) or exocoelomic cavity (ExC) approximately E7.5, corresponding to Theiler stages 11b–11d (TS11b–TS11d). In brief, a pregnant mouse was anesthetized with isoflurane and placed on a heating table in a supine position. 0.1 mg/kg Buprenorphine painkiller was injected subcutaneously. A 1–2 cm vertical midline incision was made in the lower abdomen. The uterus was gently exposed and drawn through the bottom of a commercial modified petri dish (Visual Sonics, # SA-11620) with an elastic membrane (Visual Sonics, # SA-11054) in the middle.

The petri dish was then filled with PBS to allow imaging of the embryos within the uterus. With the real-time guidance of ultrasound (Visual Sonics, Vevo 2100, # VS-20047, with 70MHz MS Series transducer, Visual Sonics, # MS700), a specified volume of lentivirus was injected using a capillary needle attached to a Nanoject II (Drummond, #3-000-205A) or Nanoject III (Drummond, #3-000-207), and a micromanipulator. After the injection, the uterus was gently put back into the abdomen. The muscular layer was sutured using 6-0 prolene and the skin was closed with EZ-clips. The whole process was performed within 30 mins. After the surgery, the female was placed into a pre-warmed cage on a 40°C heating plate and carefully monitored until fully recovered.

#### Tissue collection and processing for immunofluorescence

Pregnant females were sacrificed with CO_2_ and E16.5 embryos were sacrificed by decapitation. Embryos/ organs were collected for analysis. Injected embryos were confirmed based on the expression of H2B-GFP or tdTomato, with a stereomicroscope using fluorescent illumination and then dissected under a stereomicroscope (Stemi 508, ZEISS). Whole embryos between E9.5 to E11.5 were fixed in 4% PFA in PBS for 2 h and whole embryos at E12.5 were fixed for 4 h. Livers and hearts were fixed 4-6h depending on the developmental stage. After fixation, samples were washed with PBS and dehydrated with 30% sucrose. Larger samples were transferred to 15% and 30% sucrose for gradient dehydration. When the samples had sunk to the bottom, they were embedded in OCT freezing medium (O.C.T. Compound Cryo Embedding Medium, Sakura, #4583) on dry ice and stored at -80°C. 12 µm cryosections were prepared with NX70 Cryostat and collected on SuperFrost plus glass slides (VWR, #631-9483). Sections were stored at -20°C for short-term storage, or -80°C for long-term storage.

#### Immunofluorescent staining

Tissue section slides were equilibrated to room temperature and a hydrophobic barrier was created by drawing a circle around tissues with a PAP Pen (Vector Labs, #H-4000). After rehydration in PBS for 5-10 min, sections were incubated with blocking buffer (5% Donkey Serum in PBS containing 0.3% TritonX-100) for 1 h at room temperature. Primary Antibody was incubated overnight at 4°C. The following day, the slides were washed 3x15 min in PBS. Secondary antibody and DAPI were incubated for 1 h at room temperature. After washing with PBS for 3x15 min, slides were mounted with Fluoroshield mounting medium (Sigma, Cat #S6182) and stored at 4°C. Images were acquired using a LSM880 or LSM980 confocal microscope. Primary Antibody dilutions: ALCAM (1:500, Thermo Fisher Scientific, #14-1661-82), CD31 (1:100, bd Biosciences, #550274), Desmin (1:500, Abcam, #ab80503); GFP (1:1,000, Abcam, #ab13970), PDPN (ready-to-use, Dako, Agilent, #IRO72), WT1 (Abcam, #AB89901, 1:500), pan-keratin (1:100, Dako, Agilent, #Z0622), Secondary Antibodies (Alexa Fluor secondary antibodies from Thermo Fisher, Alexa Fluor 488 AffiniPure Donkey Anti-Chicken IgY (IgG) (H+L), Jackson Immuno Research, #AB_2340375) and DAPI dilutions: 1:1000.

#### Tissue collection and processing for Visium HD

Pregnant females were sacrificed with CO_2_ and E16.5 embryos were sacrificed by decapitation. Livers and hearts were collected. After dissection in cold PBS under a stereomicroscope (Stemi 508, ZEISS), fresh livers and hearts were immediately transferred to OCT compound pre-chilled at 4 ℃ and snap frozen in pre-cooled isopentane (VWR, # #24872.323) with liquid nitrogen to better preserve tissue morphology. Samples were stored in -80 ℃ until cryosectioning.

For each sample, serial 10 µm cryosections were prepared using an NX70 cryostat, and one section was collected at the center of each SuperFrost Plus glass slide (VWR, #631-9483). Sections were collected throughout the entire sample. The slides were stored at −80°C until use.

#### Tissue dissociation for cell sorting

E9.5 and E10.5 whole embryos and E16.5 livers and hearts were collected for scRNA sequencing (**Materials and Methods Table 1**). Embryo dissection was performed in cold PBS under a stereomicroscope. After dissection, E9.5 and E10.5 embryos were incubated in 1mL of PBS with 1mg/mL Collagenase D (Merck, Cat # 11088866001), 0.05mg/ mL DNase I (Sigma, Cat #DN25) and 3% fetal bovine serum (FBS) (Thermo Fisher Scientific, Cat # 10500064) at 37°C for 20-30 min for different developmental stages, with occasional gentle pipetting a few times through 200 uL tips. After dissociation, cells were washed with ice-cold PBS containing 3% FBS. E16.5 livers were dissociated in 2 mL of PBS with 1mg/mL Collagenase D, 0.5 mg/mL Pronase E (Merck, Cat #P5147), 0.05 mg/ mL DNase I and 3% FBS at 37°C for 30min. E16.5 hearts were dissociated using Pierce™ Primary Cardiomyocyte Isolation Kit (ThermoFisher Scientific, #88281), according to the manufacturer’s instructions. Red blood cells were removed by incubation with 1X RBC lysis buffer (Biolegend, #420302) for 7 min at room temperature.

Then cells were washed with PBS with 0.5% BSA and 0.01mg/mL DNaseI and centrifuged at 350 g for 5 min at 4°C. The cell pellet was resuspended in PBS with 0.5% BSA and 0.01 mg/mL DNaseI and passed through a 40 µm strainer. Cells were stained with viability dye (Zombie Yellow™ Fixable Viability Kit, #423104, BioLegend), BV510 for 30 min at 4°C and washed once with cold PBS.

#### Cell sorting and library preparation

Single cell suspensions were prepared as described above and then transferred to round-bottom FACS tubes for cell sorting. Cell sorting was performed on BD FACS Aria Fusion or Aria III with 100 μm nozzle. All tdTomato positive cells were sorted into 1.5 mL Eppendorf tubes containing PBS with 0.5% BSA and 0.01 mg/ml DNaseI. All sorted samples were centrifuged at 350 g for 7 min at 4°C. The supernatant was carefully discarded, and cells were resuspended in ∼30 uL of PBS with 0.5% BSA. Cells were counted with a hemocytometer at the microscope to estimate the cell concentration and total cell number. Library preparation was performed following the manufacturer’s protocol. Samples were processed with 10X Genomics Chromium Single Cell Kit Version 3.1 (with single index) or Version 4 (with dual index). Cells were mixed with 10X Chromium RT mix and loaded in 10X Chromium Next GEM Chip G. Single cell gel beads in emulsion were generated with 10X Chromium Controller. After GEM-RT incubation and post GEM-RT cleanup, cDNA amplification was performed with 12 cycles in a Thermocycler. cDNA and post library quality control were determined on Agilent Bioanalyzer. DNA concentration was measured with Qubit dsDNA HS and BR Assay Kits on Qubit 4 (Thermo Fisher Scientific).

#### Data normalization and cell filtering for scRNA-seq

The pipeline for data processing is shown in **Figures S2, S7, S12**. All cells were sequenced using Nova Seq 6000 or NovaSeq X. Fastq files were provided by the NGI facility in Sweden. A custom reference consisting of the GRCm38(mm10) and an additional sequence representing the tdTomato-N transgene, in which the barcode region was marked with “N”) was used. The fastq files were processed with Cell Ranger, to obtain gene expression matrices. The Seurat V4.5.3 package^67^ in RStudio and Python was used for data analysis.

Each dataset was processed separately. Firstly, SoupX^68^ was used to remove ambient RNA and obtain an adjusted matrix. The adjusted matrix (min.cells = 3, min.features = 200) was loaded in Seurat (v5^69^) as a Seurat object (**Figures S2, S7, S12**, Adjusted object), which was then processed with initial filtering: cells expressing low gene features and low RNA count were filtered out (cells were excluded based on the following criteria: nCount_RNA less than 1000 to 2500 & nFeature_RNA less than 1000 to 3000 depending on the sequencing depth in each sample). Low quality cells were discarded (mitochondrial >5% or 15%, high ribosomal gene expression ratio). Cells with higher than 60,000-100,000 RNA counts (depending on the sequencing depth in each sample) were considered doublets and discarded. We then used DoubletFinder ^70^ to remove estimated doublets based on initial clustering in individual samples. Furthermore, we again used Scrublet package ^71^ in Python and discarded cells with a doublet_score higher than 0.2. The clean objects were saved for further analysis.

#### Unsupervised clustering and annotation

Multiple integration approaches were tested for the E9.5/E10.5 datasets, including mapping both injected datasets onto the non-injected reference dataset. Because the reference dataset was composed of fewer cells than the injected samples, there was a risk of biasing clustering and thereby missing cell types that were not present in the reference dataset. The integration of E9.5 and E10.5 datasets (AC injections, ExC injections, and non-injected controls) were integrated (in Figure 1). Because the ExC condition consists of subsets of populations from whole embryo, the ExC dataset was analysed separately from uninjected controls to avoid integration artefacts caused by unequal cell-type representation across conditions.

E16.5 liver and heart datasets were also integrated with non-injected livers/hearts. All processing and integration were performed in R 4.5.3. We applied the SCTransform V2 normalization method ^72^, and principal components analysis (PCA) for linear dimensionality reduction. The datasets were then integrated using IntegrateLayers function with Reciprocal PCA (RPCA) integration method. The integrated data were subsequently used for clustering and Uniform Manifold Approximation and Projection (UMAP) visualization. Major cell identities were manually assigned to clusters based on canonical markers and with the guidance of the spatial deconvolution with Visium HD samples using the scRNA-seq data as a reference.

#### Extraction of cloneIDs and clone calling for scRNA-seq

We further adapted the previously described TREX pipeline ^60^. Only filtered cells (described above) were used for downstream TREX analysis. A custom reference genome consisting of the GRCm38 (mm10) genome and an additional sequence representing the tdTomato and cloneID was used. The aligned sequencing reads were then analyzed with TREX version 0.3 (0.3v50e8eb1) (https://github.com/frisen-lab/TREX) to reconstruct clones found in each dataset. A detailed description of the algorithm was presented in Ratz et al. ^60^. The version used in this study has new features that improve CloneID retrieval as well as clone assignment. Firstly, reads aligned to the CloneID sequence are filtered out if they had low complexity (e.g. only As), less than 7 read characters or if they appeared in an exclusion list of overrepresented CloneIDs, in order to minimize the risk of barcode collision (list generated from barcodes over-represented in the sequenced plasmid library and barcodes over-represented present across different experimental datasets). After a consensus sequence for each molecule is obtained, error-correction of CloneIDs was run on a per-cell basis (instead of against the whole dataset) using a maximum allowed Hamming distance of 5. Clone assignment was performed using a Jaccard index threshold of 0.6. An extra step of doublet detection was added in which cells were removed from clones and flagged if the graph resulting from their removal results in two or more connected components.

Clones identified with TREX are later added to [Seurat object] for downstream analysis. CloneIDs only identified in one cell with one read were discarded. CloneIDs that were supported by only one UMI and have a high frequency in another cell were also considered to be contamination and were discarded.

#### Clonal coupling scores and correlation analysis

To assess the overrepresentation and underrepresentation of cell types sharing clones, clonal coupling Z-scores were calculated as described in detail in ^73^. The analysis utilizes cell identity and clonal information, comparing the obtained clonal data to a randomized dataset, which we generated by shuffling the dataset 100, 500, 1,000 or 5,000 times, with three independent runs for each number of permutations. The number of shuffles was selected by sequentially increasing it and keeping the value at which differences between matrices stabilized. The E16.5 liver and heart datasets exhibited variability at 100 shufflings, and 5,000 shufflings were stable and used for the analysis.

Z-scores were computed for each pair of cell types, representing the number of standard deviations by which the observed clonal sharing deviated from the expected (randomized) values. Positive Z-scores indicated higher-than-expected clonal sharing, while negative Z-scores indicated lower than expected sharing. Values above 2 or below -2 were considered significant. To cluster cell types with similar clonal coupling profiles, Pearson correlations of Z-scores were computed, grouping cell types based on similarities in their clonal coupling patterns (*12*). In the E16.5 liver dataset **(Figure 4G)**, blood lineages segregated to one major branch, but did not further segregate clearly along erythroid/myeloid sub-branches, possibly due to erythroid noise in the data as a consequence of blood cell lysis and possible barcode contamination, despite extensive filtering and data clean-up.

#### Spatial transcriptomics with Visium HD

10 µm fresh-frozen cryosections were prepared as described above. Two high-quality liver sections and two high-quality heart sections from different biological replicates were selected based on morphology and initial H&E staining assessment.

Sample fixation and H&E staining: The tissue sections were processed according to the RNA Rescue Spatial Transcriptomics protocol ^74^. Slides were transferred from -80 ℃ freezer to a thermocycler pre-heated to 37℃ for 1 min, and then immediately fixed using 4% methanol-free formaldehyde (ThermoFisher, #28906) for 1 min at room temperature. The slides were washed twice in 1 x PBS and incubated with a thermocycler at 37 ℃ for 20 min. After the slides had cooled down, they were stained with Hematoxylin (Dako, #S330930-2) for 3 min and Eosin (Sigma-Aldrich, #HT110216) for 1 min. The slides were briefly washed with 1× PBS, mounted with approximately 100 µL of 85% glycerol (Thermo Fisher, #15514011), covered with coverslips, and imaged at ×20 magnification using the Metafer Slide Scanning platform (microscope: AxioImager.Z2 with ScopeLED Illumination, Zeiss; camera: CoolCube 4 m, MetaSystems; objective: Plan-Apochromat ×20/0.80 M27, Zeiss; software, Metafer5). After assessment of tissue quality and morphology, the coverslips were removed by immersing the slides in a beaker of Milli-Q water.

The probe hybridization and library preparation were performed with the Visium HD Spatial Gene Expression Reagent Kits (10x Genomics), following the manufacturer’s protocol (#CG000685, Rev C). Libraries were sequenced by using the Illumina NextSeq 2000 platform, in which the length of read 1 was 43 bp and the length of read 2 was 50 bp.

Raw sequencing reads were processed using Space Ranger (10x Genomics) to generate spatially barcoded feature-barcode matrices binned at 2, 4, 8, and 16 µm resolutions. Downstream analyses were performed using the 16 µm bin size.

#### Stereoscope

Spatial transcriptomics data were deconvoluted using Stereoscope (version 0.3.1), a probabilistic framework for estimating the spatial proportions of cell populations using annotated scRNA-seq reference data (https://github.com/almaan/stereoscope)^25^. Stereoscope analyses were performed using raw transcript counts, and all deconvolutions were run for 50,000 iterations.

E9.5/E10.5 scRNA-seq data were used as reference to deconvolute whole-embryo Stereo-seq section E9.5_E1S1 from the mouse organogenesis spatiotemporal transcriptomic atlas (MOSTA^24^).

E16.5 spatial analyses were performed on 10x Genomics Visium HD data, processed by Space Ranger v3.1.3, using a bin size of 16 µm. E16.5 liver scRNA-seq following E7.5 exocoelomic cavity injection were deconvolved onto two E16.5 liver Visium HD sections. Owing to the large size of the liver scRNA-seq reference and spatial transcriptomic liver datasets and compute restraint, the reference scRNA-seq dataset was down-sampled to a maximum of 500 cells per cluster by retaining the cells with the highest number of detected features (nFeature). The 5,000 most highly variable genes (HVGs) were used for deconvolution.

For heart spatial analysis, the E16.5 AC and ExC scRNA-seq populations were deconvolved separately onto two E16.5 heart Visium HD sections using all available scRNA-seq cells using a bin size of 16 µm.

To resolve valve mesenchymal subpopulations, ExC and AC scRNA-seq datasets with the valve mesenchyme subclustered (cl.20 for E16.5 ExC, cl.16 for E16.5 AC) were deconvoluted separately on cropped heart section 1 corresponding to the valve regions. Subsequently, the integrated ExC and AC scRNA-seq data were jointly used as a combined Stereoscope reference, allowing for harmonized spatial localization of valve mesenchymal cells. All valve mesenchyme deconvolutions were performed using a bin size of 16 µm.

For visualization, individual cell-population plots displayed the full range of inferred Stereoscope proportions across all spatial spots, whereas combined multi-population plots were restricted to the top 5% of spots by inferred proportion for each population to improve spatial contrast and reduce visual overlap.

#### Clonal embedding analysis using clone2vec

Clonal relationships were analyzed using clone2vec (version 2.0.0) (https://github.com/kharchenkolab/clone2vec^75^). Clone2vec learns low-dimensional representations of individual clones from the local transcriptomic states occupied by clonally related cells, independently of predefined cell-type annotations. Cells were assigned to clones using cloneIDs. Clones containing ≥ 2 cells were retained for E9.5 and E10.5 clonal analyses. Clones containing ≥ 5 cells were retained for E16.5 ExC liver clonal analyses. Clones containing ≥3 cells were retained for E16.5 ExC heart and ≥2 for E16.5 AC heart analyses. Sequencing libraries were additionally assigned to embryo-specific biological replicates using an explicit replicate mapping such that libraries originating from the same embryo shared a common replicate identifier.

A kNN graph was constructed using k = 15 and was used to derive clone-level adjacency. Clone2vec embeddings were trained with a 10-dimensional latent space, batch size of 4,096, learning rate of 0.001, and maximum of 500 epochs. Embeddings were initialized by singular value decomposition, with an early stopping rate after three epochs without a loss improvement of ≥1 x 10^-4^. UMAP (minimum distance = 0.1, spread = 1.0) and Leiden clustering were performed for downstream analyses. Neural crest enrichment fold in the amniotic cavity versus exocoelomic cavity was calculated using clone2vec clone size ≥ 2 (resolution 0.1) (ExC 66,799 cells, 1,874 neural crest clones; AC 37,403 cells, 5,470 clones).

#### Clonal coupling undirected lineage network

Clonal coupling between transcriptomic states was quantified by building on the lineage-coupling framework described by Bander et al., in “*Single-cell delineation of lineage and genetic identity in the mouse brain*” (https://github.com/mayer-lab/Bandler-et-al_lineage^36^), itself based on the approach of Wagner and colleagues^76^ . In this framework, the contribution of each clone to a pair of cell states is normalized by the total size of that clone, and the observed coupling is standardized relative to a permutation-derived null distribution. We adapted this framework to account explicitly for biological replicates and to construct an undirected statistically filtered clonal-coupling lineage network.

For each embryo-specific clone *c* and transcriptomic cluster *x*, we defined

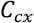

As the number of cells from clone *c* assigned to cluster *x*. The total size of clone *c* was

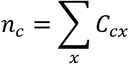

For a pair of transcriptomic clusters *x* and *y*, clone *c* was considered shared only when at least one cell from the clone was present in each cluster

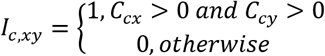

For a shared clone, the number of clone-derived cells occupying either member of the cluster pair was

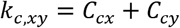

and its contribution to the pairwise coupling was

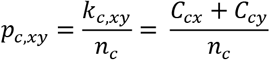

The clonal coupling score between clusters x and y was therefore

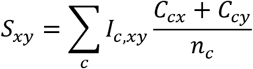

To assess whether the observed clonal coupling between transcriptomic clusters was greater than expected by chance, we created a biological replicate (embryo) -aware permutation null distribution based on the framework of Bandler et al. Cluster labels were permuted independently within each biological replicate (embryo), preserving the number of cloneID-positive cells assigned to each transcriptomic state within each replicate. Transcriptomic cluster labels were randomly permutated among cloneID-positive cells within each replicate while the clone assignment, clone sizes, cluster abundances and total number of cloneID-positive cells remained fixed.

For each cluster pair *x*, *y*, the coupling score was recalculated across 10,000 independent permutations

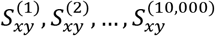

The mean coupling score expected under the null model was then calculated as, where *S_xy_*^(*r*)^ is the coupling score from the *r*-th permutation

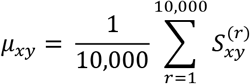

The standard deviation of the null distribution was calculated as

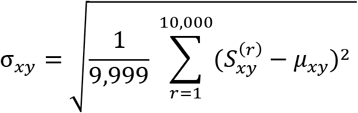

The coupling score was then standardized relative to the permutation-derived null distribution as

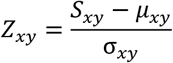

Thus, *Z_xy_* > 0 indicates that the clonal coupling between transcriptomic clusters *x* and *y* was greater than the mean coupling expected under the replicate-specific permutation null model, whereas *Z_xy_* < 0 indicates a coupling below the null model expectation. Values close to zero indicate coupling similar to the expected null model.

In addition to the Z-score framework, we assessed whether coupling was significantly greater than expected under the permutation model. We calculated an empirical enrichment *P* value for each cluster pair, where the +1 correction prevents zero empirical *P* values.

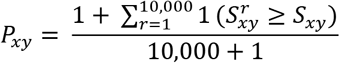

*P* values for all unique transcriptomic cluster pairs were then corrected for multiple testing using Benjamini-Hochberg false discovery rate (FDR). *P* values were ordered from smallest to largest where *m* is the largest rank

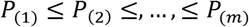

For each ranked *P* value, where *i* denotes the rank index of a particular *P* value in this ordered list an initial FDR-adjusted value *A* was calculated as

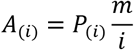

The final FDR-adjusted value *q* is then obtained after monotonic correction where each initial FDR-adjusted value *A*_(*i*)_was replaced by the smallest *A* value at its rank or any rank below

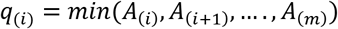

Finally, an edge between two transcriptomic cluster nodes in the network is only plotted if all three requirements are fulfilled:

(1) *Z_xy_* > 0;

(2) The number of clones contributing to that edge ≥ 10;

(3) *q_xy_* ≤ 0.05

### QUANTIFICATION AND STATISTICAL ANALYSIS

Statistical details of individual experiments, the exact test used, the value of *n*, and what n represents, are reported in the corresponding figure legends. Quality-control filtering (see *Data normalization and cell filtering for scRNA-seq* sections) and cloneID extraction (see *Extraction of cloneIDs and clone calling for scRNA-seq*) were performed using deterministic threshold-based criteria. Cluster identities were assigned manually from canonical marker expression and cross-validated against spatial deconvolution data (see above). Cell-group markers were identified using FindAllMarkers (Seurat v4.5.3/v5), applying a two-sided Wilcoxon rank-sum test to SCT-normalized expression values, with *p*-values corrected for multiple comparisons across all tested genes using the Bonferroni method; only positively enriched markers were reported for each group.

Two related, but distinct clonal-coupling analyses were used in this study. Clonal coupling Z-scores (see *Clonal coupling scores and correlation analysis*) were calculated relative to randomized null distribution generated by shuffling clone- and cell-identity data and were considered significant at |Z| > 2, without correction for multiple comparisons across cell-type pairs. The clonal-coupling network (see *Clonal coupling undirected lineage network)* extended this framework using a biological-replicate-stratified permutation null, generated from 10,000 permutations of cluster labels within each of the biological replicates, and Benjamini-Hochberg correction applied across all 990 cluster pairs tested. An edge was included in the network only when Z_xy_ > 0, at least 10 clones contributed to the pair, and the FDR-corrected *q*-values were ≤ 0.05. For both analyses, the center and dispersion of the null distribution were defined as its mean and standard deviation, from which the Z-score was derived.

---

**Materials and Methods Figure 1.**
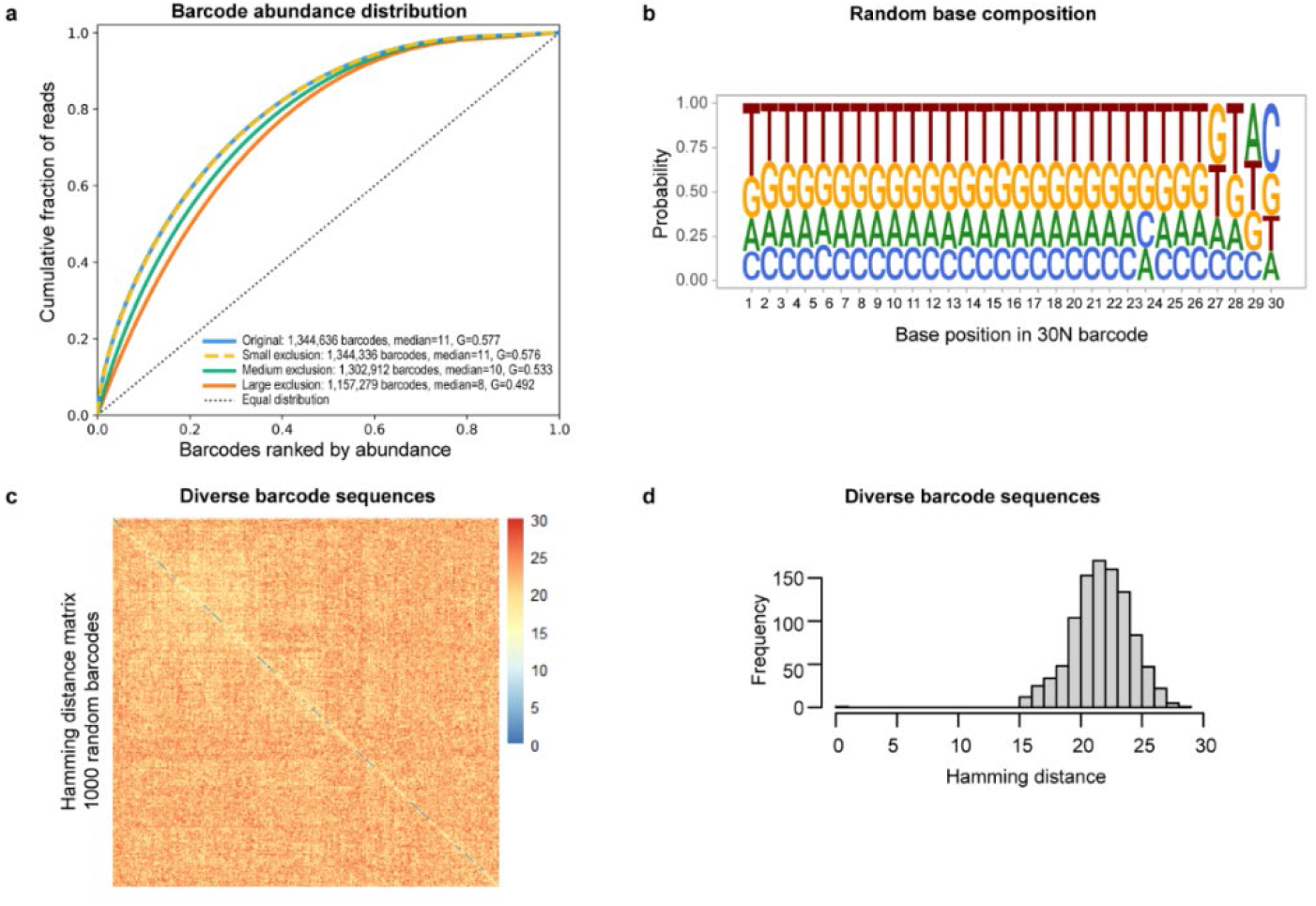
Characterization of plasmid barcode library. **a**, Barcode abundance distributions of the plasmid library before filtering and after filtering using small (barcodes have more than 365 reads), medium (barcodes have more than 73 reads), and large (barcodes have more than 37 reads) exclusion lists. The Lorenz curve shows a possible bias in the sequencing library preparation (due to e.g., library expansion, PCR amplification), as a small proportion of barcodes accumulates a higher share of total reads (e.g., 1.3% of total barcodes have more than 100 reads/barcode and represent around 10% of total reads). **b**, The sequence plot demonstrates no bias in the sequence base composition of the barcodes. The plot was calculated from 100 000 randomly selected barcodes. **c,d**, The diversity of 30N barcode sequences is estimated by Hamming distance. Matrix of Hamming distances calculated for 1000 randomly selected barcodes (**c**) and representative histogram (**d**) of a randomly selected barcode from the matrix above show the sequences are highly diverse, with Hamming distance ≥ 15 bases.

**Materials and Methods Table 1.**
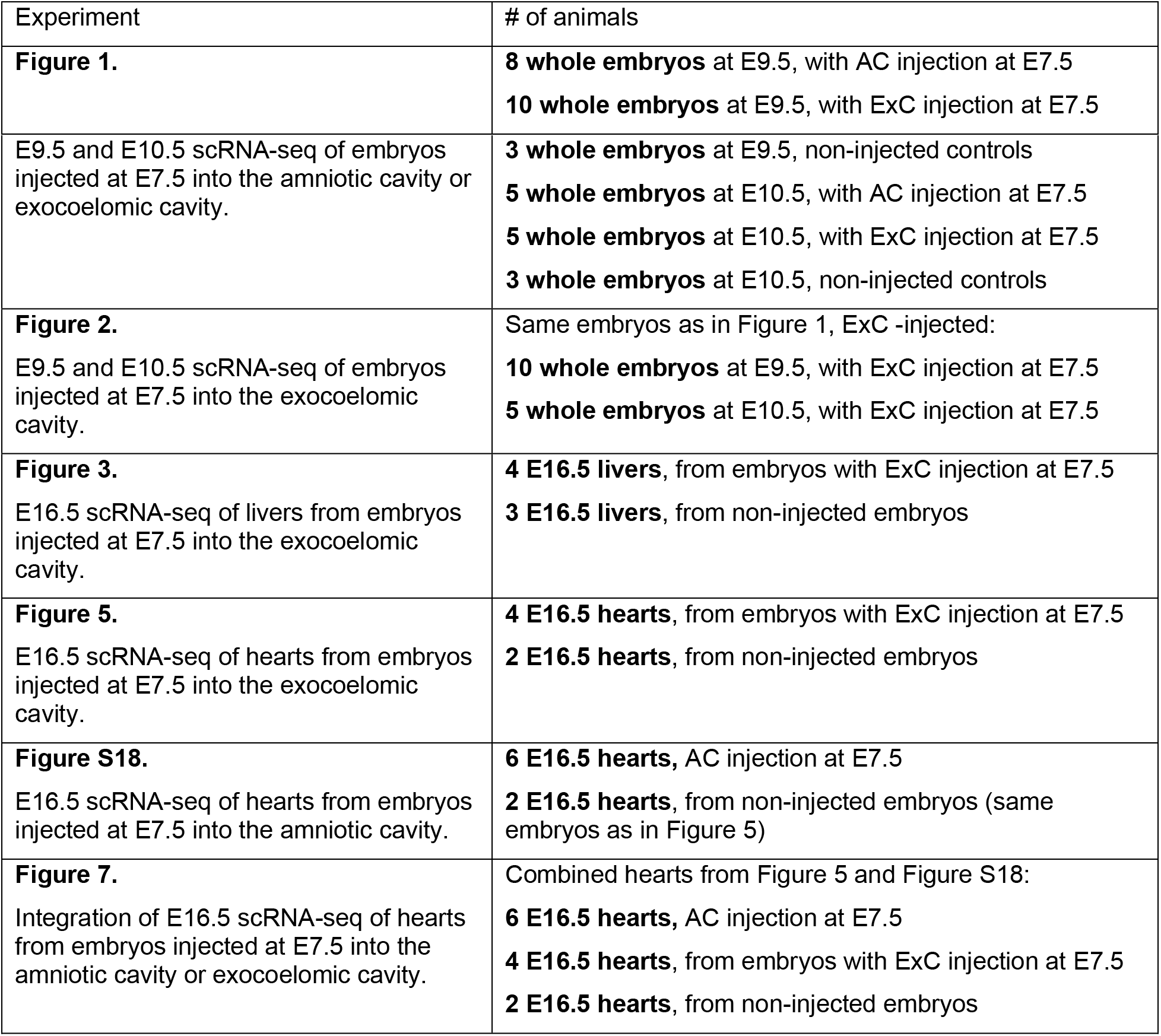
Summary of samples processed for scRNA-seq.

| Experiment | # of animals |
| --- | --- |
| **Figure 1.** | **8 whole embryos** at E9.5, with AC injection at E7.5<br>**10 whole embryos** at E9.5, with ExC injection at E7.5 |
| E9.5 and E10.5 scRNA-seq of embryos injected at E7.5 into the amniotic cavity or exocoelomic cavity. | **3 whole embryos** at E9.5, non-injected controls<br>**5 whole embryos** at E10.5, with AC injection at E7.5<br>**5 whole embryos** at E10.5, with ExC injection at E7.5<br>**3 whole embryos** at E10.5, non-injected controls |
| **Figure 2.**<br>E9.5 and E10.5 scRNA-seq of embryos injected at E7.5 into the exocoelomic cavity. | Same embryos as in Figure 1, ExC -injected:<br>**10 whole embryos** at E9.5, with ExC injection at E7.5<br>**5 whole embryos** at E10.5, with ExC injection at E7.5 |
| **Figure 3.**<br>E16.5 scRNA-seq of livers from embryos injected at E7.5 into the exocoelomic cavity. | **4 E16.5 livers**, from embryos with ExC injection at E7.5<br>**3 E16.5 livers**, from non-injected embryos |
| **Figure 5.**<br>E16.5 scRNA-seq of hearts from embryos injected at E7.5 into the exocoelomic cavity. | **4 E16.5 hearts**, from embryos with ExC injection at E7.5<br>**2 E16.5 hearts**, from non-injected embryos |
| **Figure S18.**<br>E16.5 scRNA-seq of hearts from embryos injected at E7.5 into the amniotic cavity. | **6 E16.5 hearts**, AC injection at E7.5<br>**2 E16.5 hearts**, from non-injected embryos (same embryos as in Figure 5) |
| **Figure 7.**<br>Integration of E16.5 scRNA-seq of hearts from embryos injected at E7.5 into the amniotic cavity or exocoelomic cavity. | Combined hearts from Figure 5 and Figure S18:<br>**6 E16.5 hearts**, AC injection at E7.5<br>**4 E16.5 hearts**, from embryos with ExC injection at E7.5<br>**2 E16.5 hearts**, from non-injected embryos |

