## Supplemental Figures for "In utero barcoding of mouse mesoderm reveals the clonal architecture of organ mesenchyme"

[1] Department of Cell and Molecular Biology, Karolinska Institutet; Stockholm, Sweden.
[2] Janelia Research Campus, Howard Hughes Medical Institute, Ashburn, VA 20147, USA.
[3] Current Affiliation: Department of Physics, Faculty of Exact and Natural Sciences, University of Buenos Aires
[4] Department of Neuroimmunology, Center for Brain Research, Medical University Vienna; Vienna, Austria.
[5] Science for Life Laboratory, KTH Royal Institute of Technology, Stockholm, Sweden
[6] Current affiliation: Department of Human Genetics, Leiden University Medical Center, Leiden, the Netherlands
[7] Department of Comparative Medicine, Karolinska Institutet; Stockholm, Sweden.
[8] Current affiliation: Department of Laboratory Medicine, Karolinska Institutet, and Preclinical Laboratory Facility, Karolinska University Hospital.
[9] Centre for Inflammation Research, Institute for Regeneration and Repair, University of Edinburgh, Edinburgh, UK.
[10] Department of Medical Biochemistry and Biophysics, Karolinska Institutet; Stockholm, Sweden.
[11] Science for Life Laboratory, Stockholm University, Sweden
[12] MRC Human Genetics Unit, Institute of Genetics and Cancer, University of Edinburgh, Edinburgh, UK.
[13] Institute of Molecular Biotechnology, Vienna, Austria
[14] Department of Physiology and Pharmacology, Karolinska Institutet; Stockholm, Sweden.
[15] These authors contributed equally.
[16] Senior author
[17] Lead contact

### Supplementary Figures S1-S22

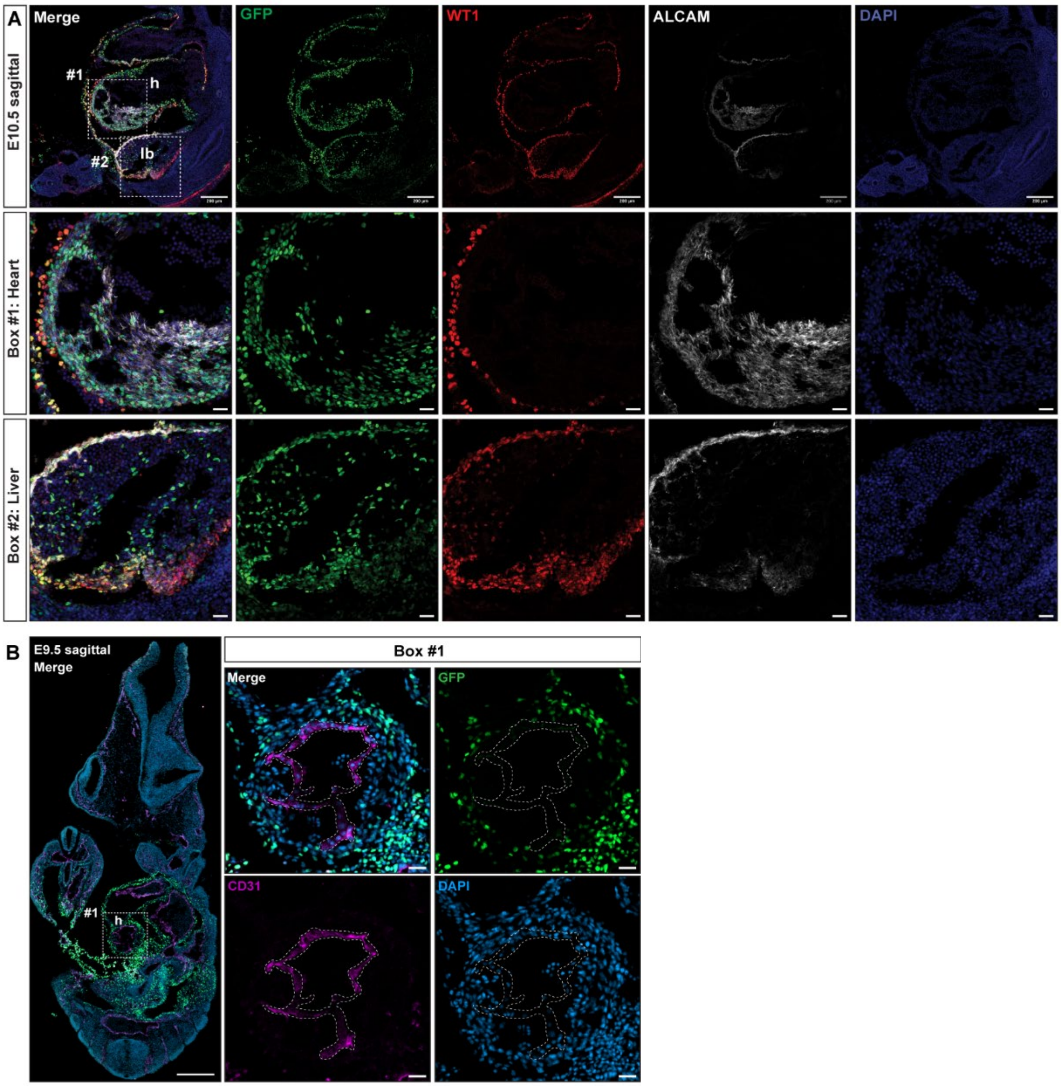


**Figure S1. Exocoelomic cavity injection of lentivirus encoding H2B-GFP at E7.5 efficiently labels the heart and liver in E10.5 mouse embryos, related to Figure 1.**

**(A)** First row, sagittal view of the abdominal region of an E10.5 embryo following E7.5 exocoelomic cavity injection of H2B–GFP lentivirus, stained for WT1, ALCAM and DAPI. Second row, magnified view of Box 1, heart region. Third row, magnified view of Box 2, liver bud region.

**(B)** Left, sagittal view of a whole E9.5 embryo following E7.5 exocoelomic cavity injection of H2B-GFP lentivirus. Right, magnified view of Box 1, heart region, showing H2B-GFP-negative CD31+ endocardium (delineated by dashed lines).

Scale bars: **A**, Row 1 Scalebar = 200 µm. Row 2/3 Box Scalebars = 30 µm . **B,** left image scalebar = 300 µm, right Box 1 scalebar = 30 µm. h = heart; lb =liver bud.

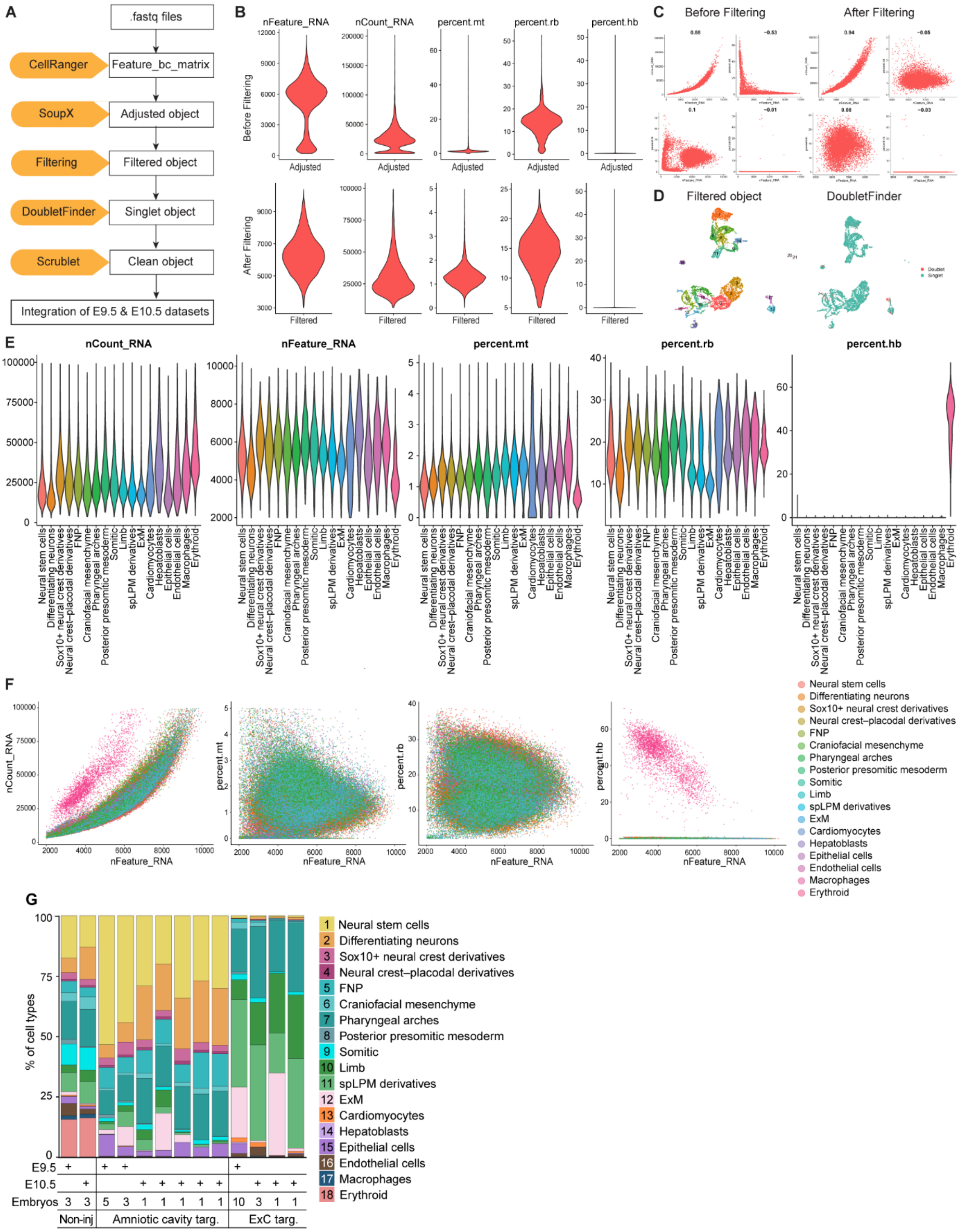


**Figure S2. Quality control for single-cell RNA-sequencing analysis of E9.5 and E10.5 embryos, and cell type abundance, related to Figure 1.**

**(A)** Workflow of the quality-control pipeline.

**(B)** Violin plots showing quality-control metrics (nFeature_RNA, nCount_RNA, the percentage of mitochondrial genes, the percentage of ribosomal genes and hemoglobin genes) for a SoupX-adjusted object, with an E9.5 amniotic cavity-injected sample shown as an example. Metrics are shown before filtering (top) and after filtering (bottom).

**(C)** Scatter plots showing the same metrics before filtering (left) and after filtering (right).

**(D)** UMAP visualization of the filtered object, showing annotated cell types (left) and DoubletFinder-defined singlets and doublets (right). Singlets are shown in blue and doublets in red; doublets were removed from downstream analysis.

**(E,F)** Violin plots (**E**) and scatter plots (**F**) showing quality-control metrics for the integrated object containing all E9.5 and E10.5 embryos, including amniotic cavity-injected, exocoelomic cavity-injected and non-injected control samples.

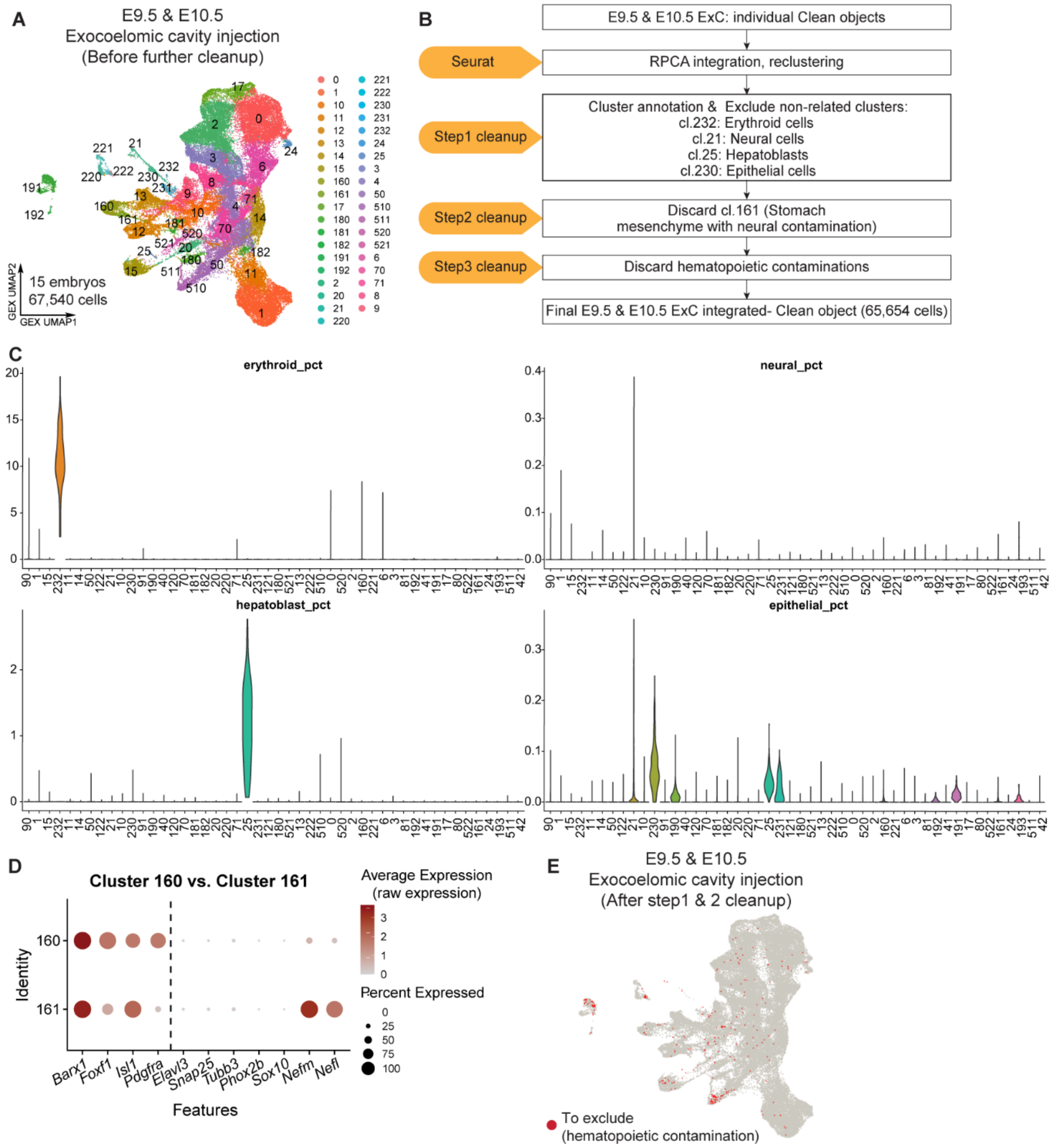


**Figure S3. Quality control for single-cell RNA-sequencing analysis of E9.5 and E10.5 embryos following E7.5 exocoelomic cavity injection, related to Figure 2.**

**(A)** UMAP visualization of the integrated object of E9.5 and E10.5 exocoelomic cavity-injected embryos before further cleanup.

**(B)** Workflow of the quality-control pipeline for E9.5 and E10.5 exocoelomic cavity-injected embryos. "Non-related clusters" exhibit hallmarks of low quality or contaminant cells, including a "straight line" appearance indicative of poor quality (low data content or ambient noise lack distinct biological variance, causing dimensionality reduction algorithms to force them into artificial, geometric linear arrangements).

**(C)** Violin plots showing the percentage of erythroid, neural, hepatoblast and epithelial lineage marker expression across clusters, corresponding to the step 1 cleanup described in **b**.

**(D)** Dot plot showing raw expression of stomach mesenchyme markers (left of the bar) and neural markers (right of the bar) in clusters 160 and 161. **e,**

**(E)** UMAP visualization showing hematopoietic contaminants in red in the integrated object.

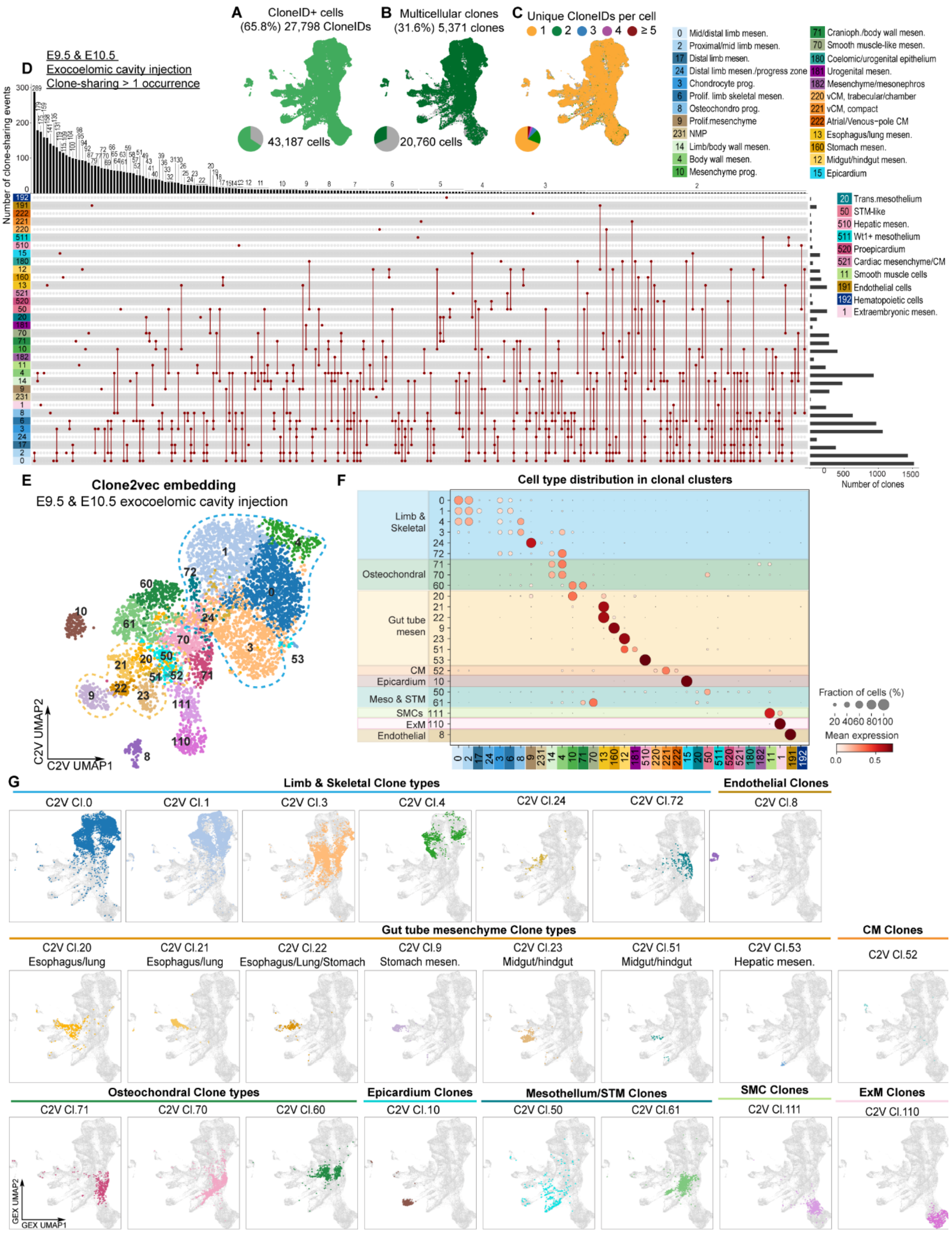


**Figure S4. Single-cell lineage tracing with E7.5 exocoelomic cavity injection resolves early mesodermal clonal relations, related to Figure 1,2.**

**(A-C)** UMAPs and pie charts showing CloneID+ cells (**A**), multicellular CloneID+ clones (clone size ≥2 cells; **B**) and the number of CloneIDs per cell (**C**) among all cells isolated from E9.5 and E10.5 embryos after E7.5 exocoelomic cavity injection.

**(D)** UpSet plot showing clonal relationships across cell types (clone-sharing >1 occurrence) in the integrated E9.5 & E10.5 dataset. The top bars indicate the number of clone-sharing events among cell types or within a single cell type, and the right bars indicate the total clone numbers containing the indicated cell types.

**(E)** clone2vec embedding of the integrated E9.5 & E10.5 dataset.

**(F)** Dot plot showing cell-type distribution across C2V clonal clusters. Dot size represents the fraction of clones containing the indicated cell type. Dot color represents the average cell-type composition of clones within each C2V clonocluster.

**(G)** Gene expression UMAP (GEX UMAP) showing the cell composition of each C2V clonal cluster.

Cranioph = craniopharyngeal, CM = cardiomyocytes, ExM = extraembryonic mesenchyme, mesen = mesenchyme, NMP = neuromesodermal progenitors, Osteochondro = osteochondrocyte, SMC = smooth muscle cells, STM = septum transversum mesenchyme, trans.mesothelium = transitional mesothelium, vCM = ventricular cardiomyocytes.

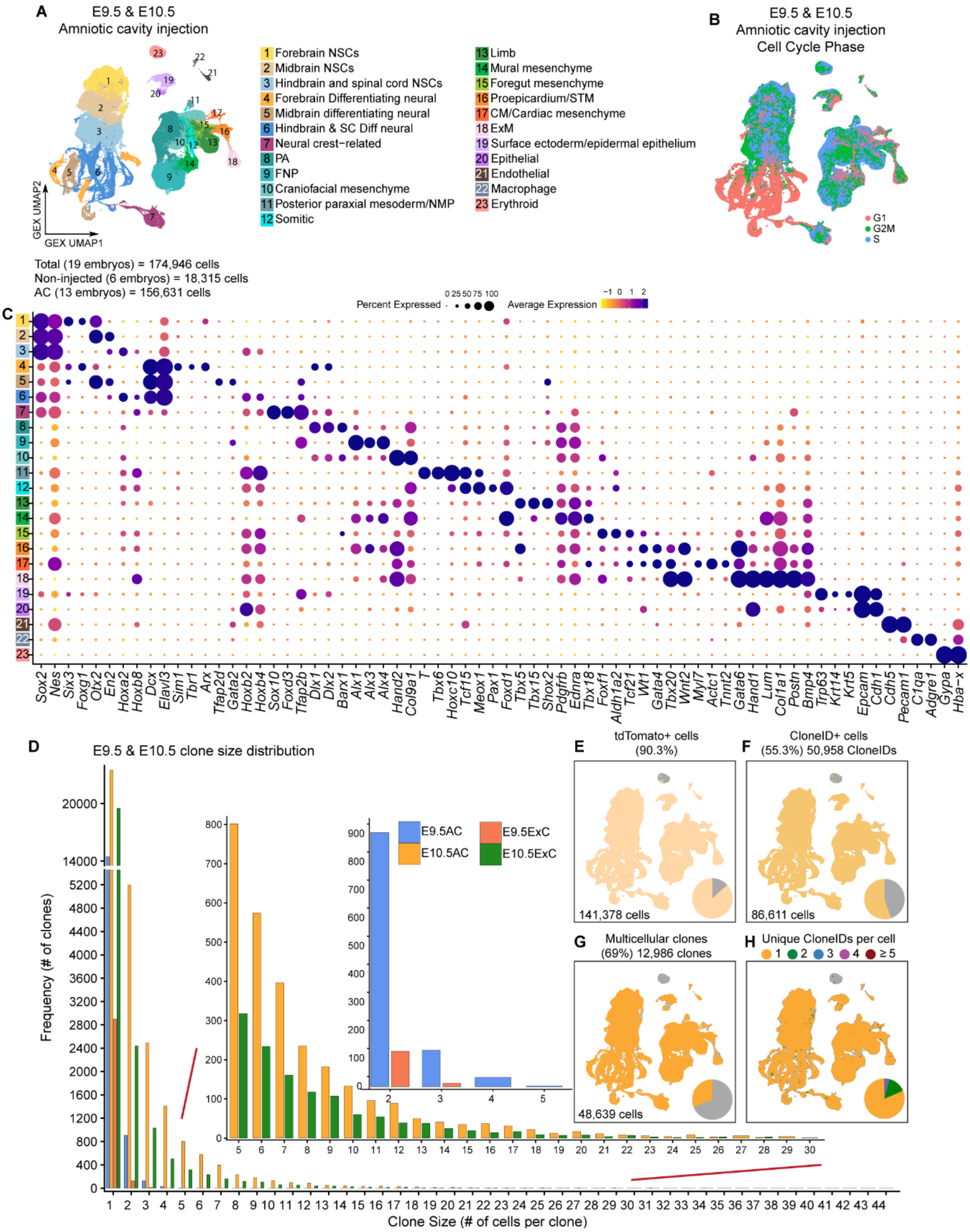


**Figure S5. E7.5 amniotic cavity injection enriches ectodermal and neural crest derivatives, related to Figure 1.**

**(A,B)** UMAPs of 174,946 cells from E9.5 and E10.5 embryos after E7.5 amniotic cavity injection, colored by annotated cell population (**A**) and cell-cycle phase (**B**). **c,**

**(C)** Expression of canonical markers used to annotate ectodermal, neural crest, mesenchymal and mesodermal populations in **a**.

**(D)** Clone-size distribution in integrated E9.5 and E10.5 embryos after E7.5 amniotic or exocoelomic cavity injection, colored by developmental stage and injection route.

**(E-H)** UMAPs and pie charts showing tdTomato+ cells (**E**), CloneID+ cells (**F**), multicellular CloneID+ clones (≥2 cells; **G**) and the number of CloneIDs per cell (**H**) among all cells isolated from E9.5 and E10.5 embryos after E7.5 amniotic cavity injection.

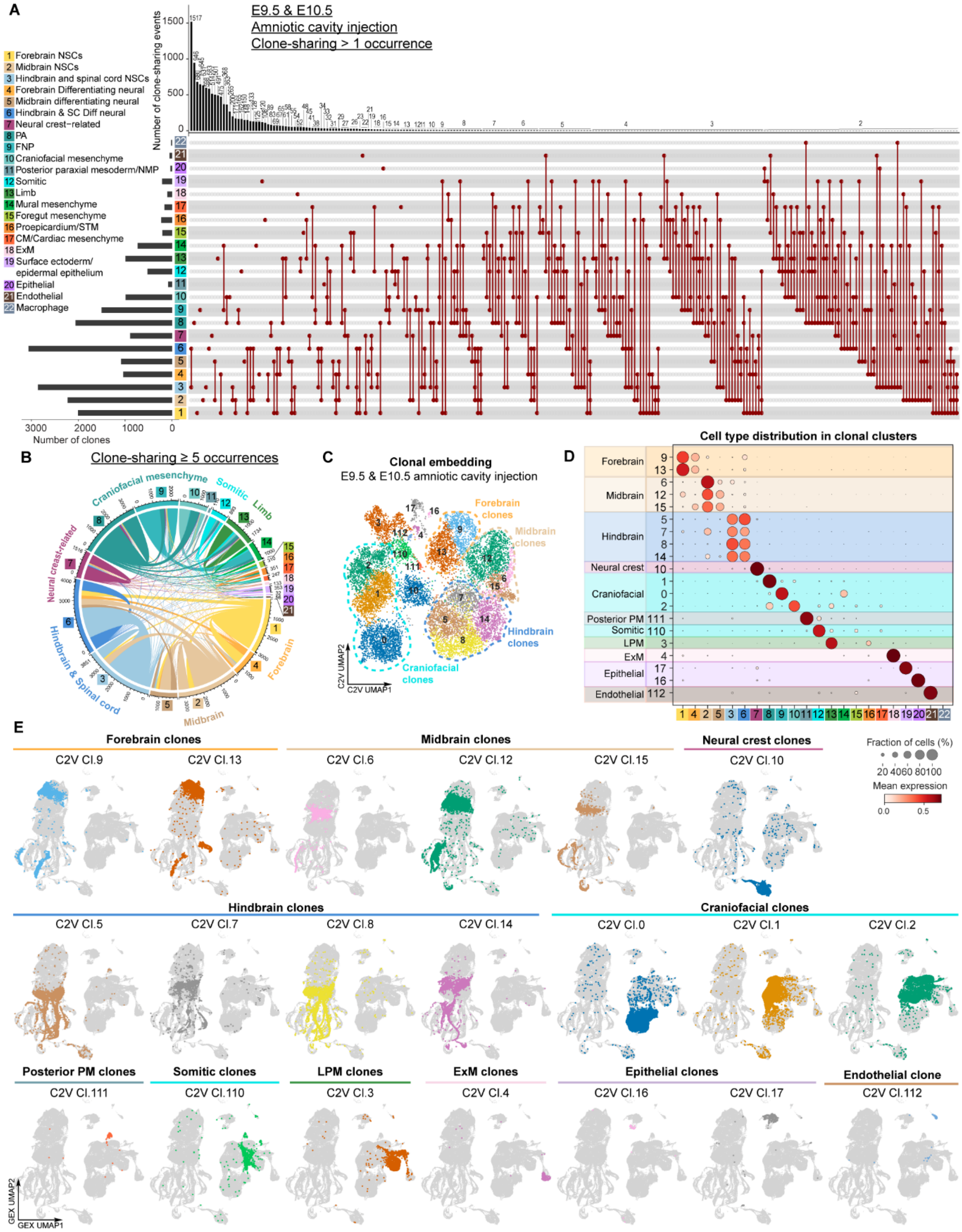
A
E9.5 & E10.5
Amniotic cavity injection
Clone-sharing > 1 occurrence
1 Forebrain NSCs
2 Midbrain NSCs
3 Hindbrain and spinal cord NSCs
4 Forebrain Differentiating neural
5 Midbrain differentiating neural
6 Hindbrain & SC Diff neural
7 Neural crest-related
8 PA
9 FNP
10 Craniofacial mesenchyme
11 Posterior paraxial mesoderm/NMP
12 Somitic
13 Limb
14 Mural mesenchyme
15 Foregut mesenchyme
16 Proepicardium/STM
17 CM/Cardiac mesenchyme
18 ExM
19 Surface ectoderm/ epidermal epithelium
20 Epithelial
21 Endothelial
22 Macrophage
Number of clone-sharing events
Number of clones
B
Clone-sharing ≥ 5 occurrences
Craniofacial mesenchyme
Somitic
Limb
Neural crest-related
Hindbrain & Spinal cord
Midbrain
Forebrain
C
Clonal embedding
E9.5 & E10.5 amniotic cavity injection
Forebrain clones
Midbrain clones
Hindbrain clones
Craniofacial clones
C2V UMAP1
C2V UMAP2
D
Cell type distribution in clonal clusters
Forebrain
Midbrain
Hindbrain
Neural crest
Craniofacial
Posterior PM
Somitic
LPM
ExM
Epithelial
Endothelial
Fraction of cells (%)
20 40 60 80 100
Mean expression
0.0 0.5
E
Forebrain clones
C2V Cl.9
C2V Cl.13
Midbrain clones
C2V Cl.6
C2V Cl.12
C2V Cl.15
Neural crest clones
C2V Cl.10
Hindbrain clones
C2V Cl.5
C2V Cl.7
C2V Cl.8
C2V Cl.14
Craniofacial clones
C2V Cl.0
C2V Cl.1
C2V Cl.2
Posterior PM clones
C2V Cl.111
Somitic clones
C2V Cl.110
LPM clones
C2V Cl.3
ExM clones
C2V Cl.4
Epithelial clones
C2V Cl.16
C2V Cl.17
Endothelial clone
C2V Cl.112
GEX UMAP1
GEX UMAP2

130

**Figure S6. Single-cell lineage tracing with E7.5 amniotic cavity injection resolves early ectodermal, mesodermal and endodermal clonal relations, related to Figure 1.**

**(A)** UpSet plot showing clonal relationships across cell types (clone-sharing >1 occurrence) in the integrated E9.5 & E10.5 dataset after E7.5 amniotic cavity injection. The top bars indicate the number of clone-sharing events among cell types or within a single cell type, and the left bars indicate the total clone numbers containing the indicated cell types.

**(B)** Circos plot showing pairwise co-occurrence of cell types across clones. Link width indicates the number of contributing cell type pairs; link color represents cell type.

**(C)** clone2vec embedding of selected multicellular clones, colored by C2V clonocluster.

**(D)** Dot plot showing cell-type distribution across C2V clonoclusters. Dot size represents the fraction of clones containing the indicated cell type. Dot color represents the average cell-type composition of clones within each C2V clonocluster.

**(E)** Gene-expression UMAP showing the cell composition of each C2V clonocluster.

CM= cardiomyocytes, ExM= extraembryonic mesenchyme, FNP= frontonasal prominence, PA= pharyngeal arch, PM= paraxial mesoderm, NSC= neural stem cells, NMP= neuromesodermal progenitors, LPM= lateral plate mesoderm, SC= spinal cord, STM= septum transversum mesenchyme.

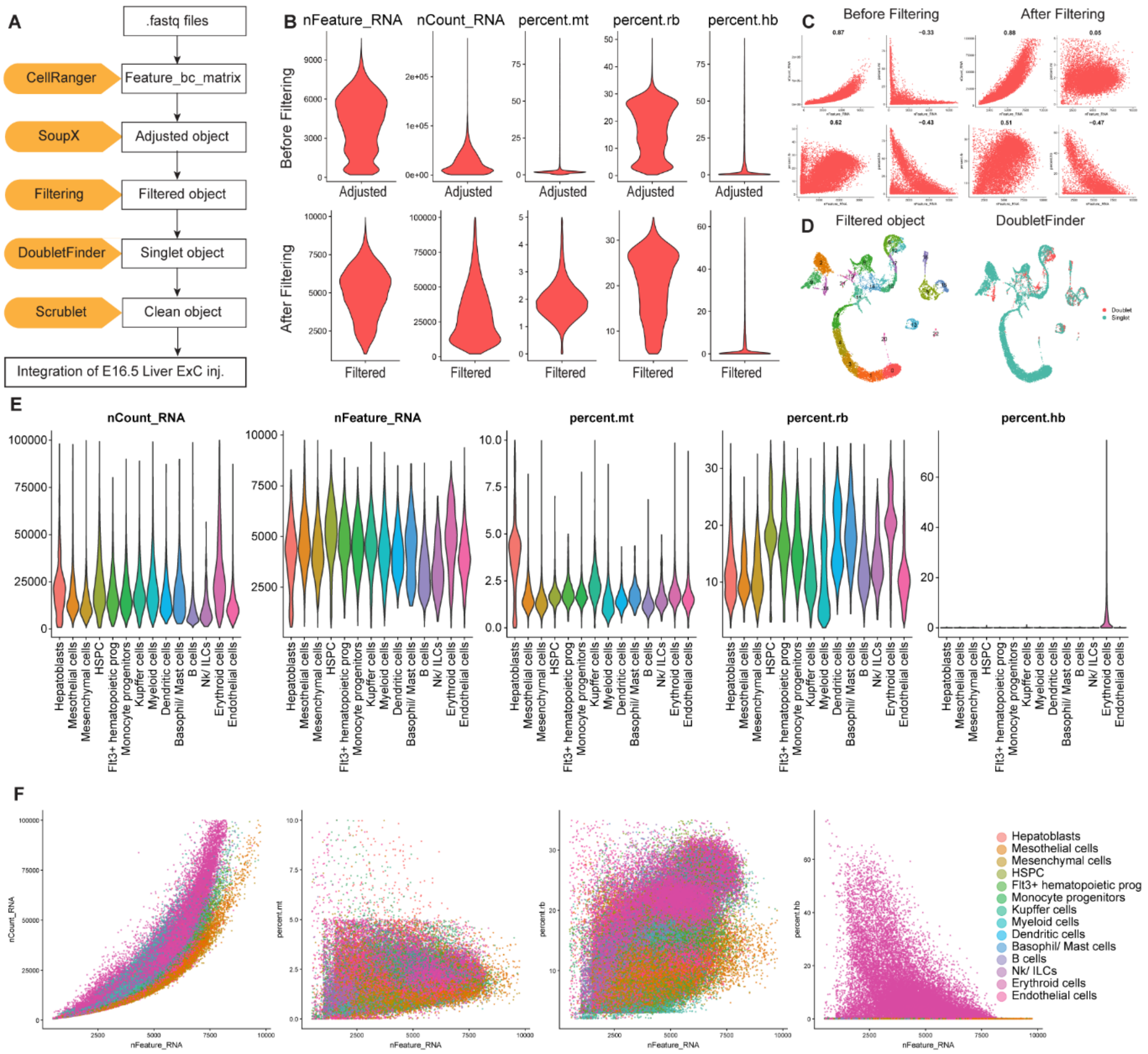


**Figure S7. Quality control for single-cell RNA-sequencing analysis of E16.5 livers following E7.5 exocoelomic cavity injection, related to Figure 3.**

**(A)** Workflow of the quality-control pipeline.

**(B)** Violin plots showing quality-control metrics (nFeature_RNA, nCount_RNA, the percentage of mitochondrial genes, the percentage of ribosomal genes and hemoglobin genes) for a SoupX-adjusted object, with one E16.5 liver sample shown as an example. Metrics are shown before filtering (top) and after filtering (bottom).

**(C)** Scatter plots showing the same metrics before filtering (left) and after filtering (right).

**(D)** UMAP visualization of the filtered object, showing annotated cell types (left) and DoubletFinder-defined singlets and doublets (right). Singlets are shown in blue and doublets in red; doublets were removed from downstream analysis.

**(E,F)** Violin plots (**E**) and scatter plots (**F**) showing quality-control metrics for the integrated object containing all liver samples.

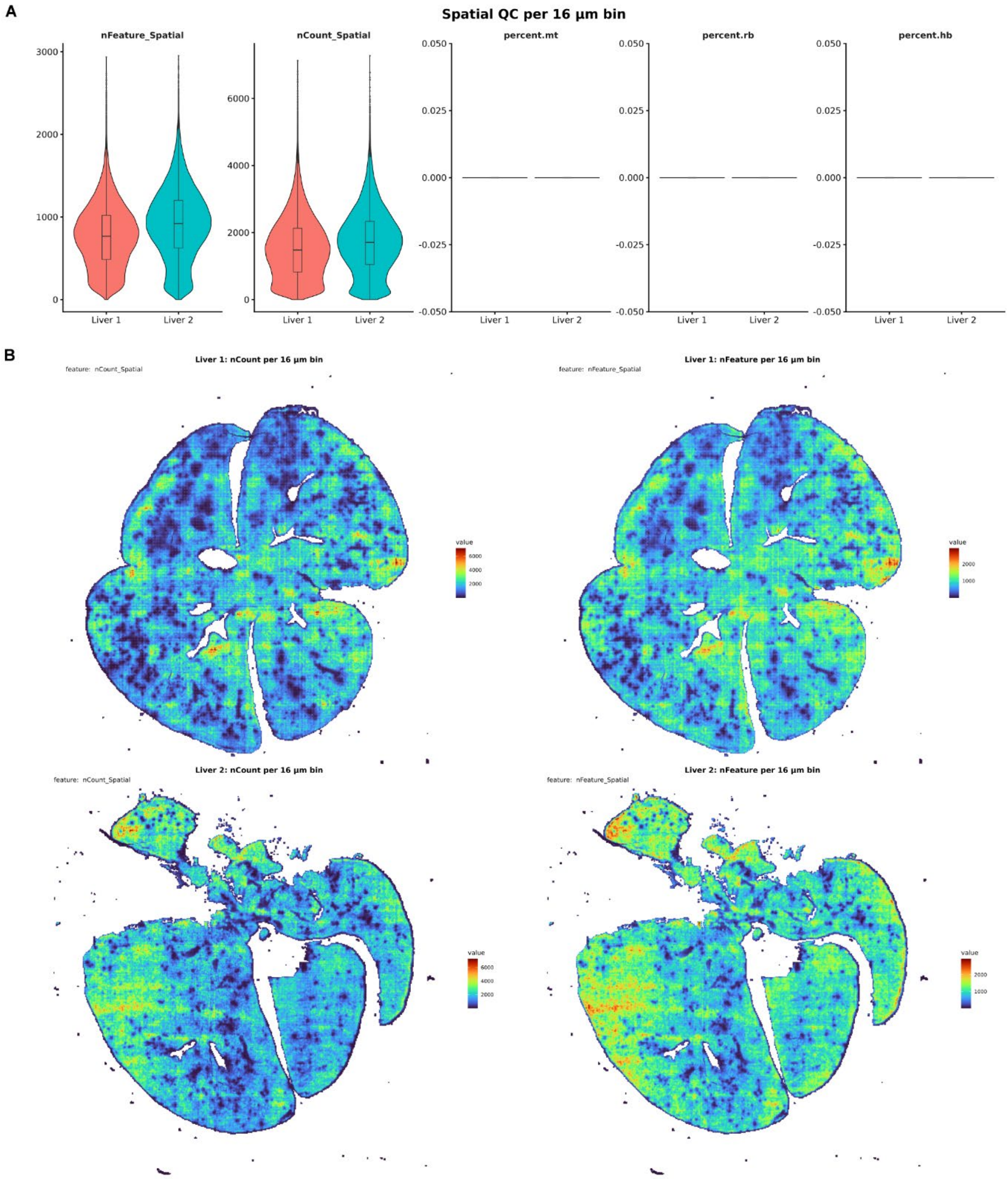


**Figure S8. Quality control for Visium HD analysis of E16.5 livers, related to Figure 3.**

**(A)** Violin plots showing quality-control metrics for E16.5 liver samples analyzed using Visium HD at 16 µm bin resolution, including the number of detected genes per 16 µm bin (nFeature_Spatial), total UMI counts per 16 µm bin (nCount_Spatial), and the percentages of mitochondrial, ribosomal, and hemoglobin gene expression.

**(B)** UMAP embedding of 16 µm bins colored by nCount_RNA and nFeature_RNA.

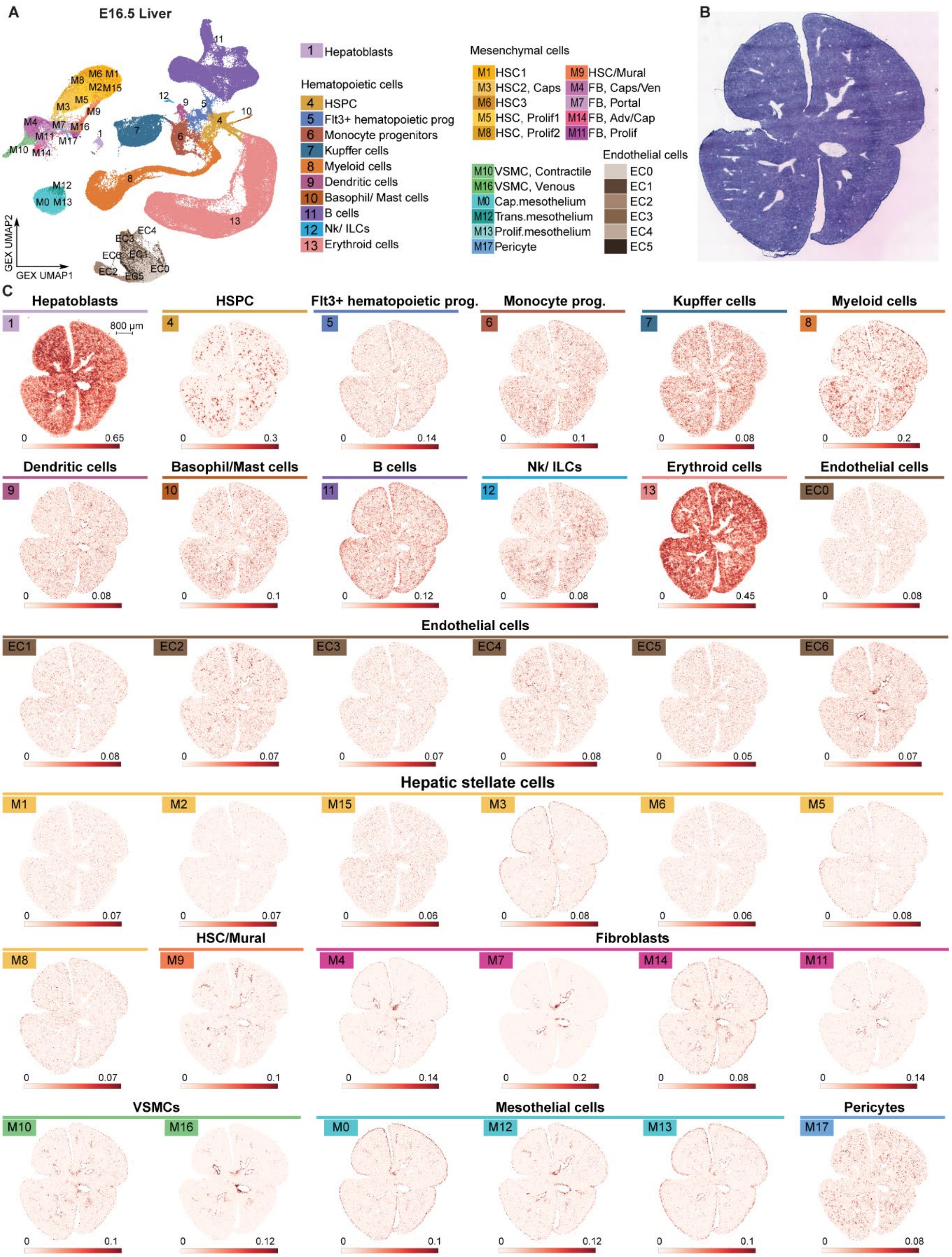
A
E16.5 Liver
GEX UMAP1
GEX UMAP2
1 Hepatoblasts
Hematopoietic cells
4 HSPC
5 Flt3+ hematopoietic prog
6 Monocyte progenitors
7 Kupffer cells
8 Myeloid cells
9 Dendritic cells
10 Basophil/ Mast cells
11 B cells
12 Nk/ ILCs
13 Erythroid cells
Mesenchymal cells
M1 HSC1
M3 HSC2, Caps
M6 HSC3
M5 HSC, Prolif1
M8 HSC, Prolif2
M9 HSC/Mural
M4 FB, Caps/Ven
M7 FB, Portal
M14 FB, Adv/Cap
M11 FB, Prolif
M10 VSMC, Contractile
M16 VSMC, Venous
M0 Cap.mesothelium
M12 Trans.mesothelium
M13 Prolif.mesothelium
M17 Pericyte
Endothelial cells
EC0
EC1
EC2
EC3
EC4
EC5
B
C
Hepatoblasts
HSPC
Flt3+ hematopoietic prog.
Monocyte prog.
Kupffer cells
Myeloid cells
800 μm
Dendritic cells
Basophil/Mast cells
B cells
Nk/ ILCs
Erythroid cells
Endothelial cells
Endothelial cells
Hepatic stellate cells
HSC/Mural
Fibroblasts
VSMCs
Mesothelial cells
Pericytes

**Figure S9. Spatial deconvolution of E16.5 liver sample #1 using liver scRNA-seq reference following E7.5 exocoelomic cavity injection, related to Figure 3.**

**(A)** UMAP visualization of the integrated object used for deconvolution, reproduced from Fig. 3B.

**(B)** Hematoxylin and eosin staining of section used for spatial transcriptomics, as an anatomical reference.

**(C)** Deconvolution of cell types. M1, M2, and M15 were deconvoluted separately but were merged as M1 (**Fig. 3H,I,R**) for clonal analyses because of their highly similar transcriptomic profiles and spatial locations.

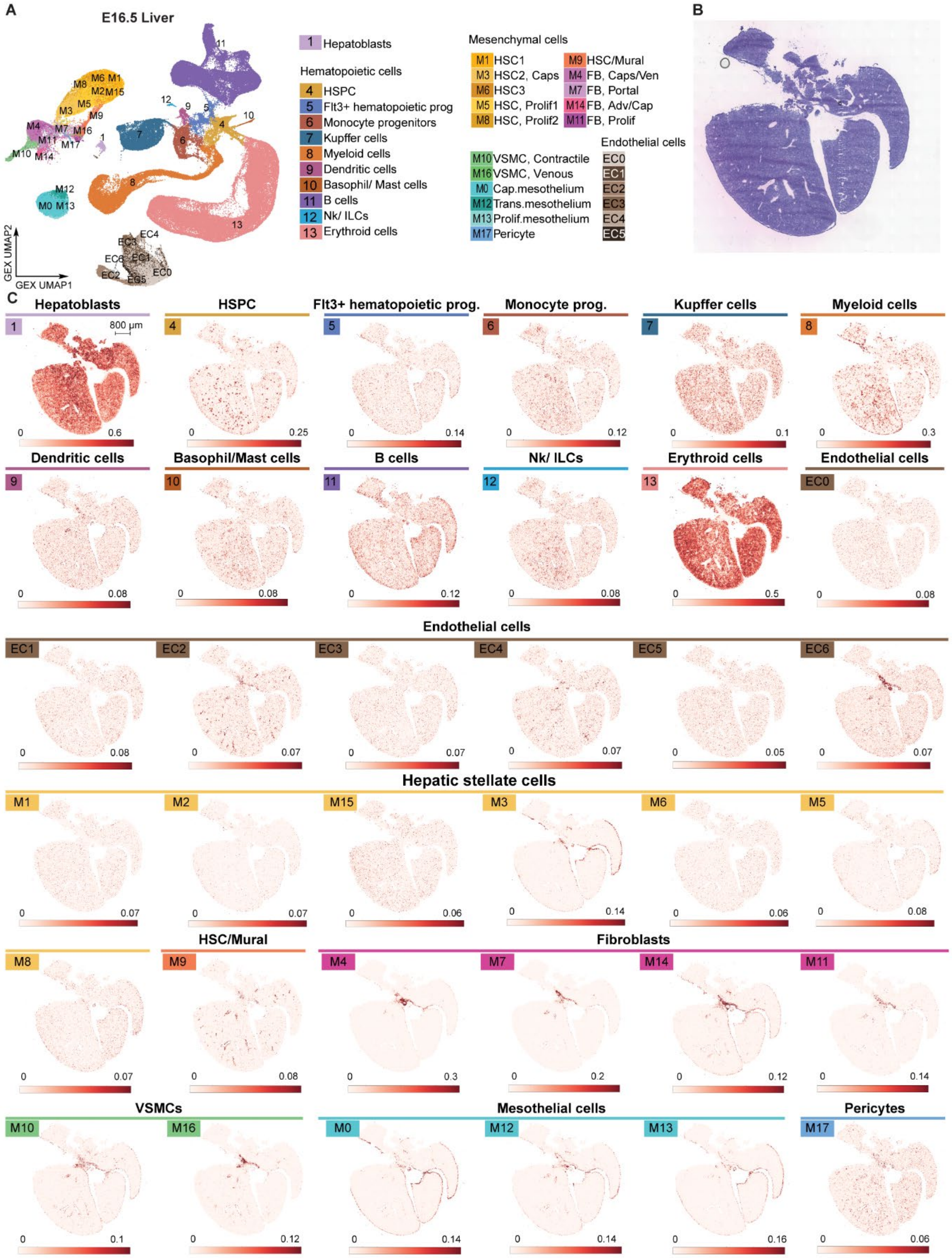

A
E16.5 Liver
GEX UMAP2
GEX UMAP1
1 Hepatoblasts
Hematopoietic cells
4 HSPC
5 Flt3+ hematopoietic prog
6 Monocyte progenitors
7 Kupffer cells
8 Myeloid cells
9 Dendritic cells
10 Basophil/ Mast cells
11 B cells
12 Nk/ ILCs
13 Erythroid cells
Mesenchymal cells
M1 HSC1
M3 HSC2, Caps
M6 HSC3
M5 HSC, Prolif1
M8 HSC, Prolif2
M9 HSC/Mural
M4 FB, Caps/Ven
M7 FB, Portal
M14 FB, Adv/Cap
M11 FB, Prolif
M10 VSMC, Contractile
M16 VSMC, Venous
M0 Cap.mesothelium
M12 Trans.mesothelium
M13 Prolif.mesothelium
M17 Pericyte
Endothelial cells
EC0
EC1
EC2
EC3
EC4
EC5
B
C
Hepatoblasts
800 μm
HSPC
Flt3+ hematopoietic prog.
Monocyte prog.
Kupffer cells
Myeloid cells
Dendritic cells
Basophil/Mast cells
B cells
Nk/ ILCs
Erythroid cells
Endothelial cells
Hepatic stellate cells
HSC/Mural
Fibroblasts
VSMCs
Mesothelial cells
Pericytes

**Figure S10. Spatial deconvolution of E16.5 liver sample #2 using liver scRNA-seq reference following E7.5 exocoelomic cavity injection, related to Figure 3.**

**(A)** UMAP visualization of the integrated object used for deconvolution, reproduced from Fig. 3B.

**(B)** Hematoxylin and eosin staining of section used for spatial transcriptomics, as an anatomical reference.

**(C)** Deconvolution of cell types. M1, M2, and M15 were deconvoluted separately but were merged as M1 (**Fig. 3H,I,R**) for clonal analyses because of their highly similar transcriptomic profiles and spatial locations.

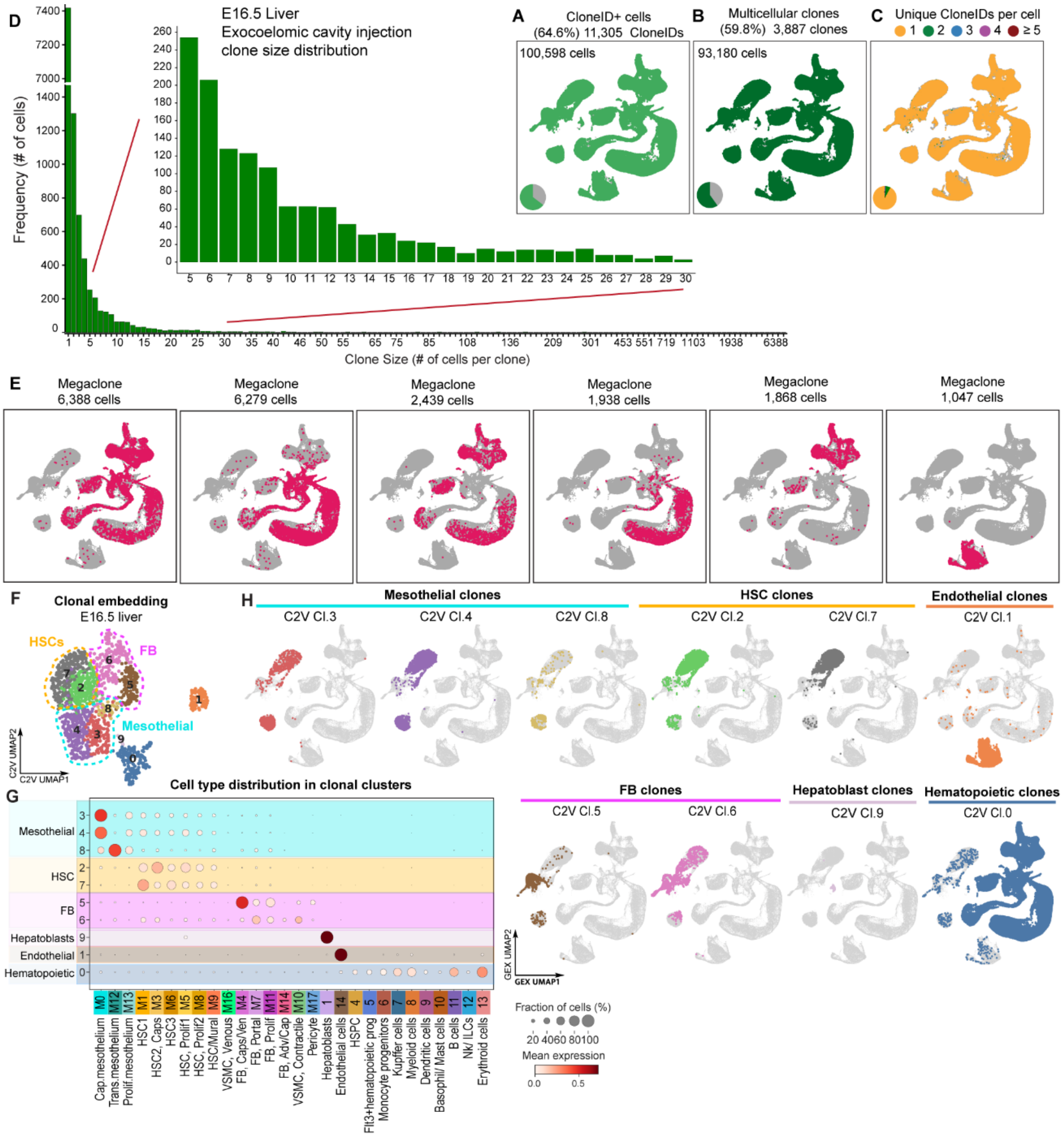


**Figure S11. Clonal characterization and analyses of E16.5 livers following E7.5 exocoelomic cavity injections, related to Figure 4.**

**(A-C)** UMAP visualizations and pie charts of CloneID+ cells (**A**) and multicellular CloneID+ cells (clones including ≥ 2 cells) (**B**), and number of cloneIDs per cell (**C**) out of all isolated cells from E16.5 livers, injected at E7.5 into exocoelomic cavity.

**(D)** Bar plot showing the clone size distribution in integrated E16.5 livers following E7.5 exocoelomic cavity injection.

**(E)** UMAP visualization of megaclones (clone size >1000 cells) in hematopoietic and endothelial populations, CloneID+ cells are in red, negative cells in grey.

**(F)** Clone2vec embedding of E16.5 liver integrated object (clone size ≥ 5) after exocoelomic cavity injection at E7.5.

**(G)** Dot plot showing the cell type distribution in C2V clonal clusters. Dot size shows the fraction of cell types in each C2V cluster and the color scale shows the expression level of each cell type in C2V cluster.

**(H)** Gene expression (GEX) UMAP visualization of the cell composition in each C2V clonoclusters.

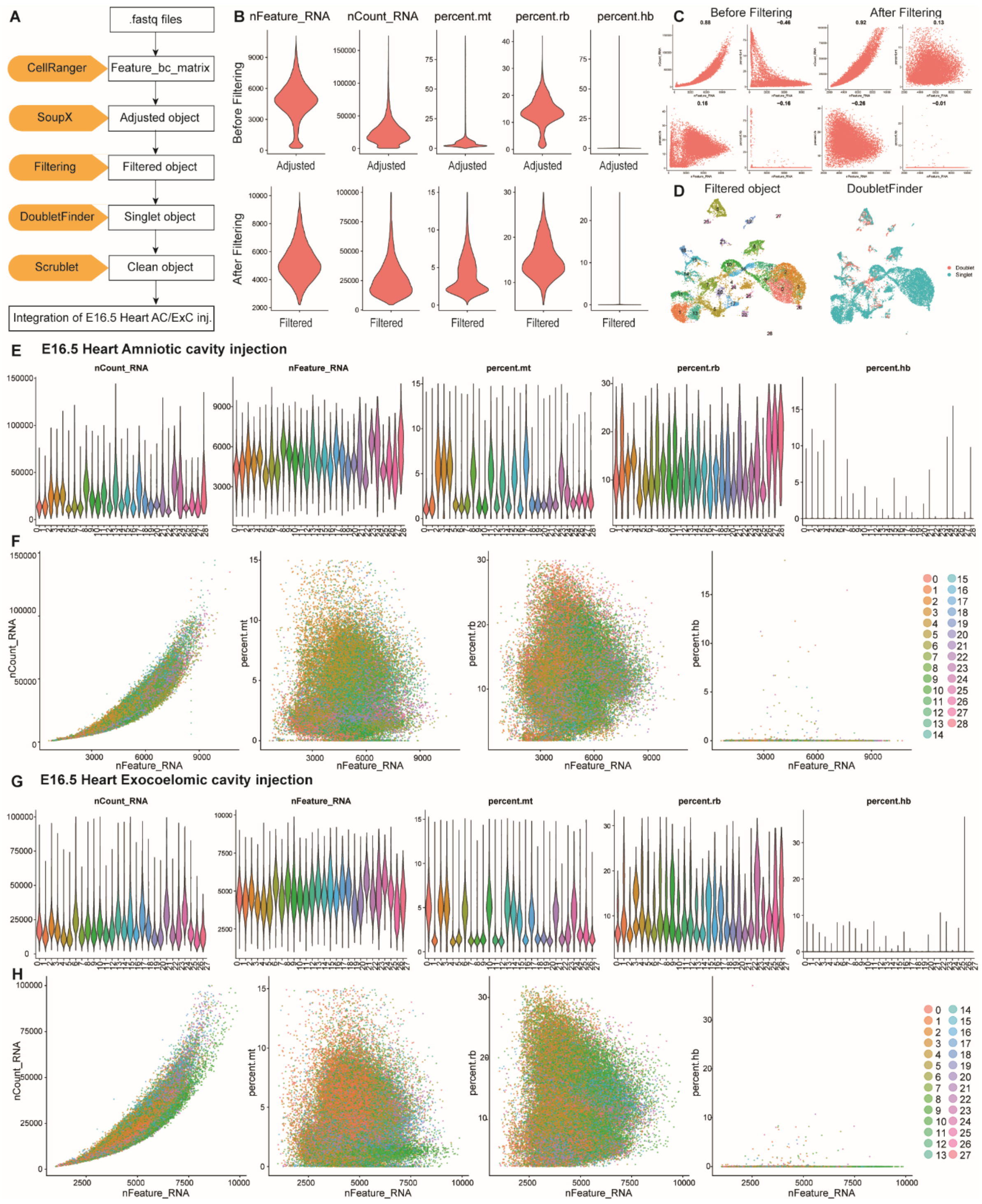


**Figure S12. Quality control for single-cell RNA-sequencing analysis of E16.5 hearts following E7.5 amniotic and exocoelomic cavity injection, related to Figure 5.**

**(A)** Workflow of the quality-control pipeline.

**(B)** Violin plots showing quality-control metrics (nFeature_RNA, nCount_RNA, the percentage of mitochondrial genes, the percentage of ribosomal genes and hemoglobin genes) for a SoupX-adjusted object, with one E16.5 heart sample shown as an example. Metrics are shown before filtering (top) and after filtering (bottom).

**(C)** Scatter plots showing the same metrics before filtering (left) and after filtering (right).

**(D)** UMAP visualization of the filtered object, showing annotated cell types (left) and DoubletFinder-defined singlets and doublets (right). Singlets are shown in blue and doublets in red; doublets were removed from downstream analysis.

**(E,F)** Violin plots (**E**) and scatter plots (**F**) showing quality-control metrics for the E16.5 heart integrated object, including amniotic cavity-injected and non-injected control samples.

**(G,H)** Violin plots (**G**) and scatter plots (**H**) showing quality-control metrics for the E16.5 heart integrated object, including exocoelomic cavity-injected and non-injected control samples.

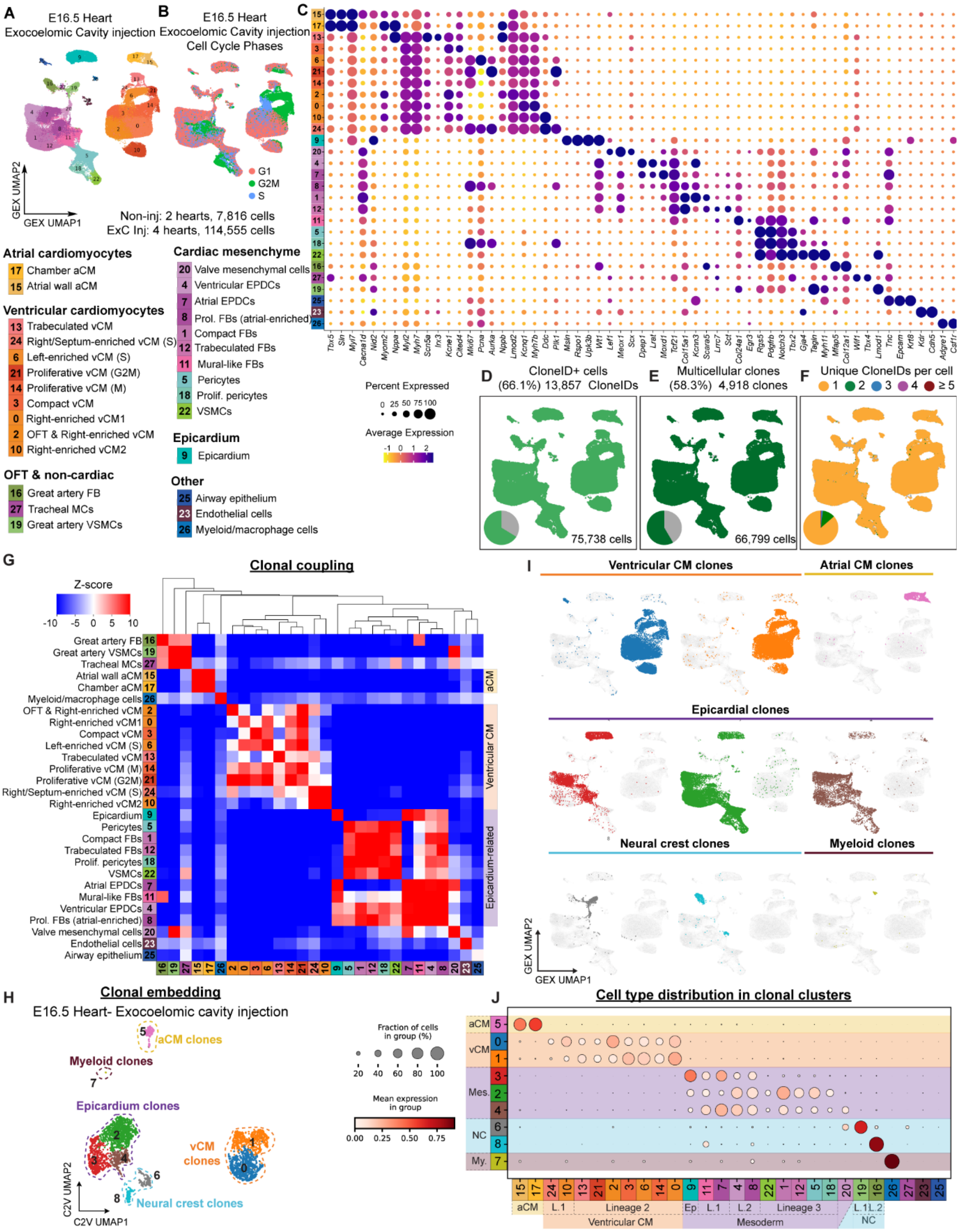

240

**Figure S13. Clonal characterization and analysis of E16.5 hearts following E7.5 exocoelomic cavity injections, related to Figure 6.**

**(A,B)** UMAPs of E16.5 heart integrated object after E7.5 exocoelomic cavity injection, colored by annotated cell population (**A**) and cell-cycle phase (**B**).

(C) Expression of canonical markers used to annotate the populations in **(A)**.

**(D-F)** UMAP visualizations and pie charts of CloneID+ cells (**D**) and multicellular CloneID+ cells (clones including 2 or more cells (**E**), and number of cloneIDs per cell (**F**) out of all isolated cells from E16.5 hearts, injected at E7.5 into exocoelomic cavity.

**(G)** Heatmaps of lineage coupling Z-scores between pairs of all cell types in E16.5 hearts following E7.5 exocoelomic cavity injections, clustered by correlation distance.

**(H)** Clone2vec embedding of E16.5 heart integrated object (clone size $\geq$ 3) after exocoelomic cavity injection at E7.5.

**(I)** Gene expression (GEX) UMAP visualization of the cell composition in each C2V clonoclusters.

**(J)** Dot plot showing the cell type distribution in C2V clonal clusters. Dot size shows the fraction of cell types in each C2V cluster and the color scale shows the expression level of each cell type in the C2V cluster.

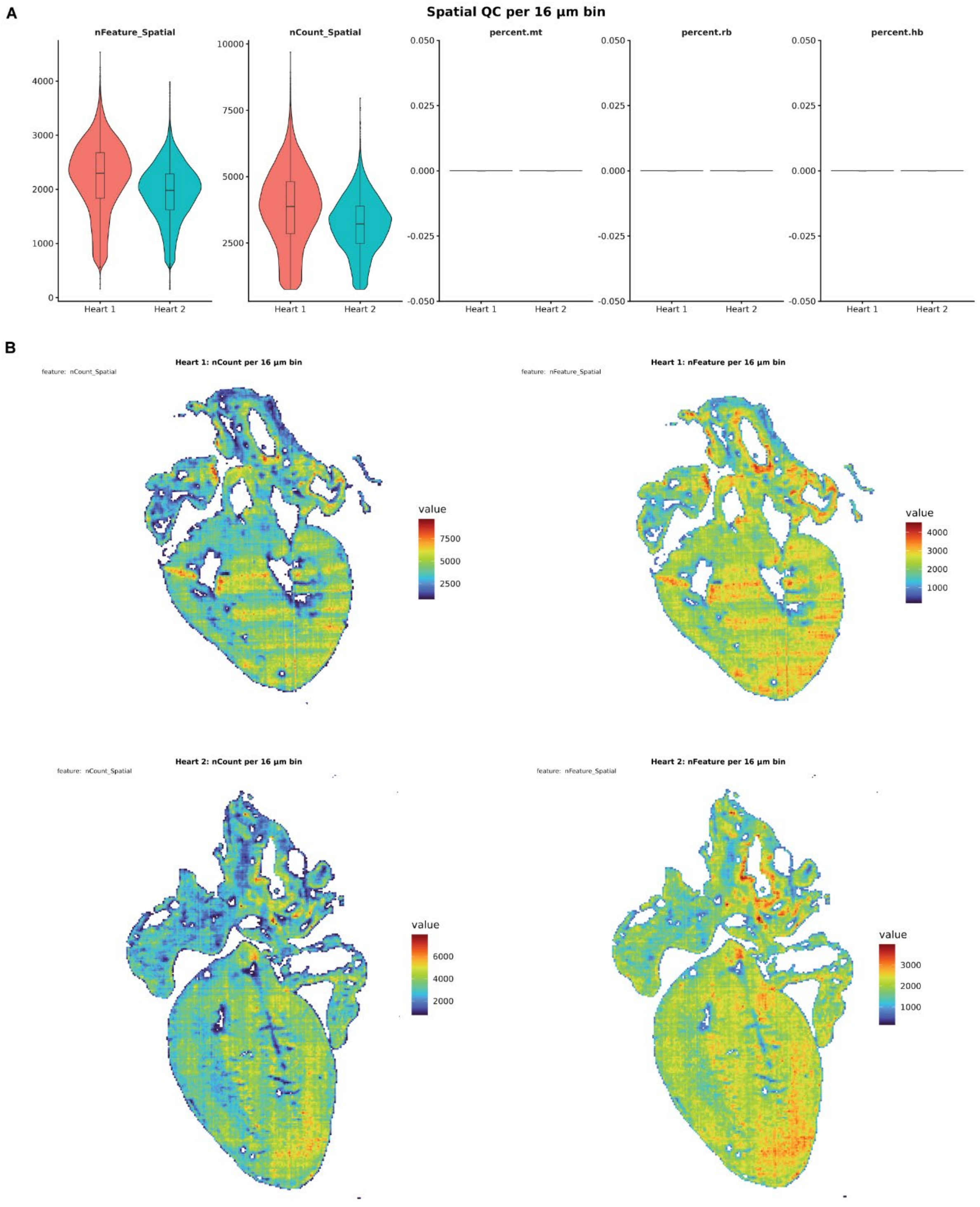


**Figure S14. Quality control for Visium HD analysis of E16.5 hearts following E7.5 exocoelomic cavity injection, related to Figure 5.**

**(A)** Violin plots showing quality-control metrics for E16.5 heart samples analyzed using Visium HD at 16 µm bin resolution, including the number of detected genes per 16 µm bin (nFeature_Spatial), total UMI counts per 16 µm bin (nCount_Spatial), and the percentages of mitochondrial, ribosomal, and hemoglobin gene expression.

**(B)** UMAP embedding of 16 µm bins colored by nCount_RNA and nFeature_RNA.

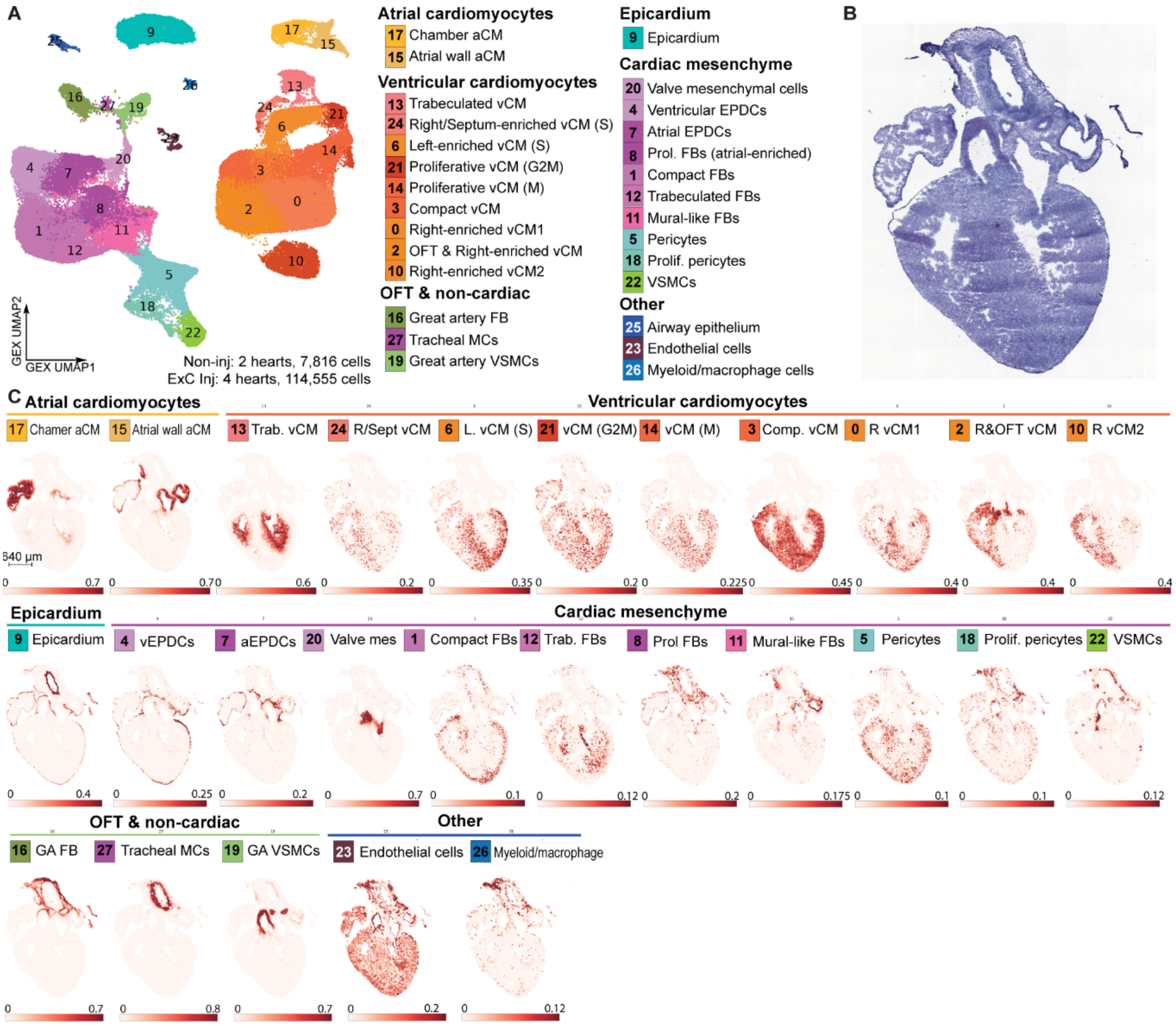


**Figure S15. Spatial deconvolution of E16.5 heart sample #1 using heart scRNA-seq reference following E7.5 exocoelomic cavity injection, related to Figure 5.**

**(A)** UMAP visualization of the integrated object used for deconvolution, reproduced from **Fig. 5B**.

**(B)** Hematoxylin and eosin staining of section used for spatial transcriptomics, as an anatomical reference.

**(C)** Deconvolution of cell types.

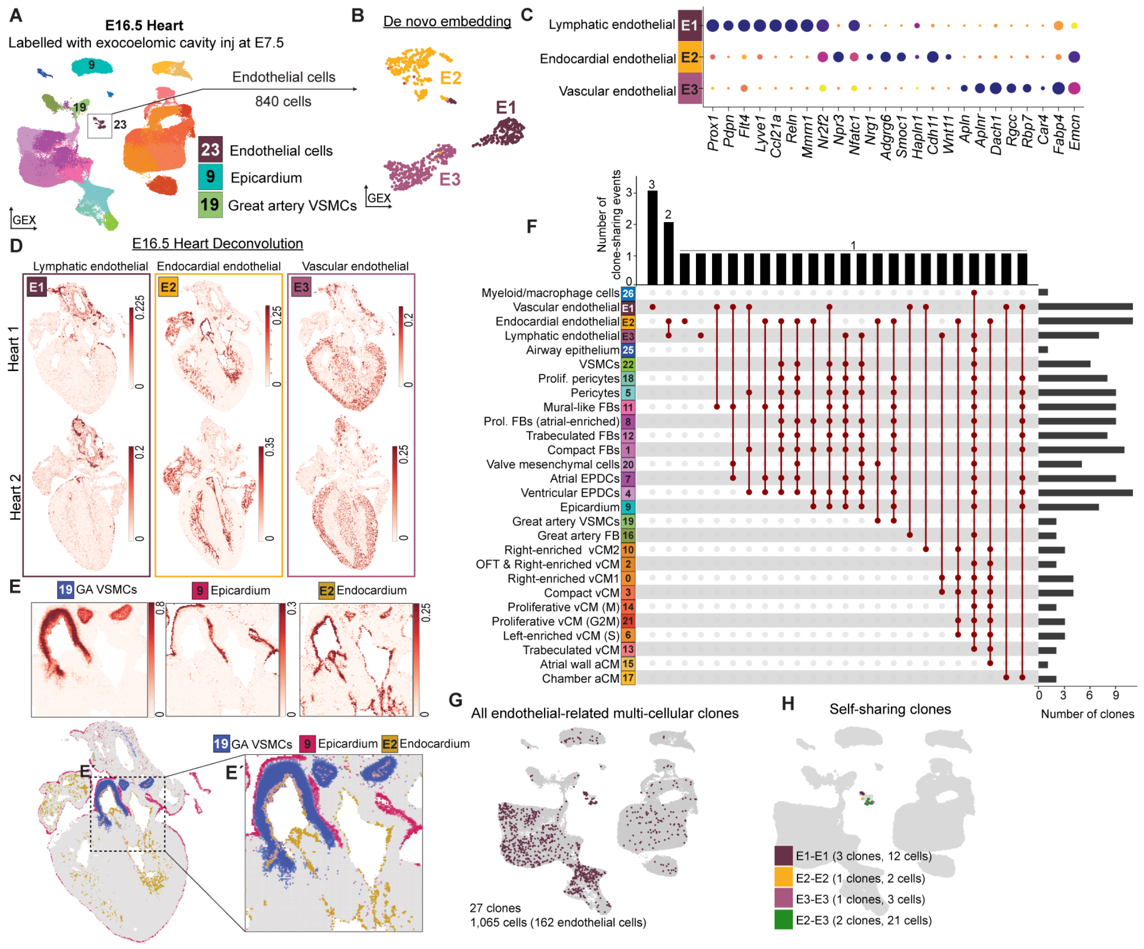


**Figure S16. Clonal characterization and analysis of endothelial cells including endocardium in E16.5 hearts following E7.5 exocoelomic cavity injections, related to Figure 6.**

**(A,B)** UMAP visualization of E16.5 heart following E7.5 exocoelomic cavity injection **(A).** 840 endothelial cells (cl.23, boxed cluster, of which 455 from the ExC condition, 385 from non-injected controls) were re-clustered and re-embedded (E1 211 cells; E2 293 cells; E3 336 cells) **(B).**

**(C)** Expression of canonical markers used to annotate the populations in **(B)**.

**(D)** Spatial localization of re-clustered endothelial cells in **(B)**.

**(E)** Spatial localization of GA VSMCs (cl.19), Epicardium (cl.9) and endocardium (E2).

**(F)** UMAP visualization of self-sharing clones for clusters in **(B).**

**(G)** UpSet plot showing all endothelial-related clones, clonal relationships across cell types (clone-sharing ≥1 occurrence; 146 cells from multicellular clones) in E16.5 heart following E7.5 exocoelomic cavity injection. The top bars indicate the number of clone-sharing events among cell types or within a single cell type, and the right bars indicate the total clone numbers containing the indicated cell types.

**(H)** Circos plot showing pairwise co-occurrence of cell types across endothelial-related clones. Link width indicates the number of contributing cell type pairs; link color represents cell type.

**(I)** Clonal coupling analysis of E16.5 heart, with endothelial cells re-clustered. Clonal coupling Z-scores quantify the enrichment of observed barcode sharing relative to randomized datasets preserving cell-population abundances, with a positive (red) score indicating enriched coupling and a negative (blue) score indicating under-represented coupling.

**(J)** Correlation of clonal coupling profiles across heart cell populations, with endothelial cells re-clustered.

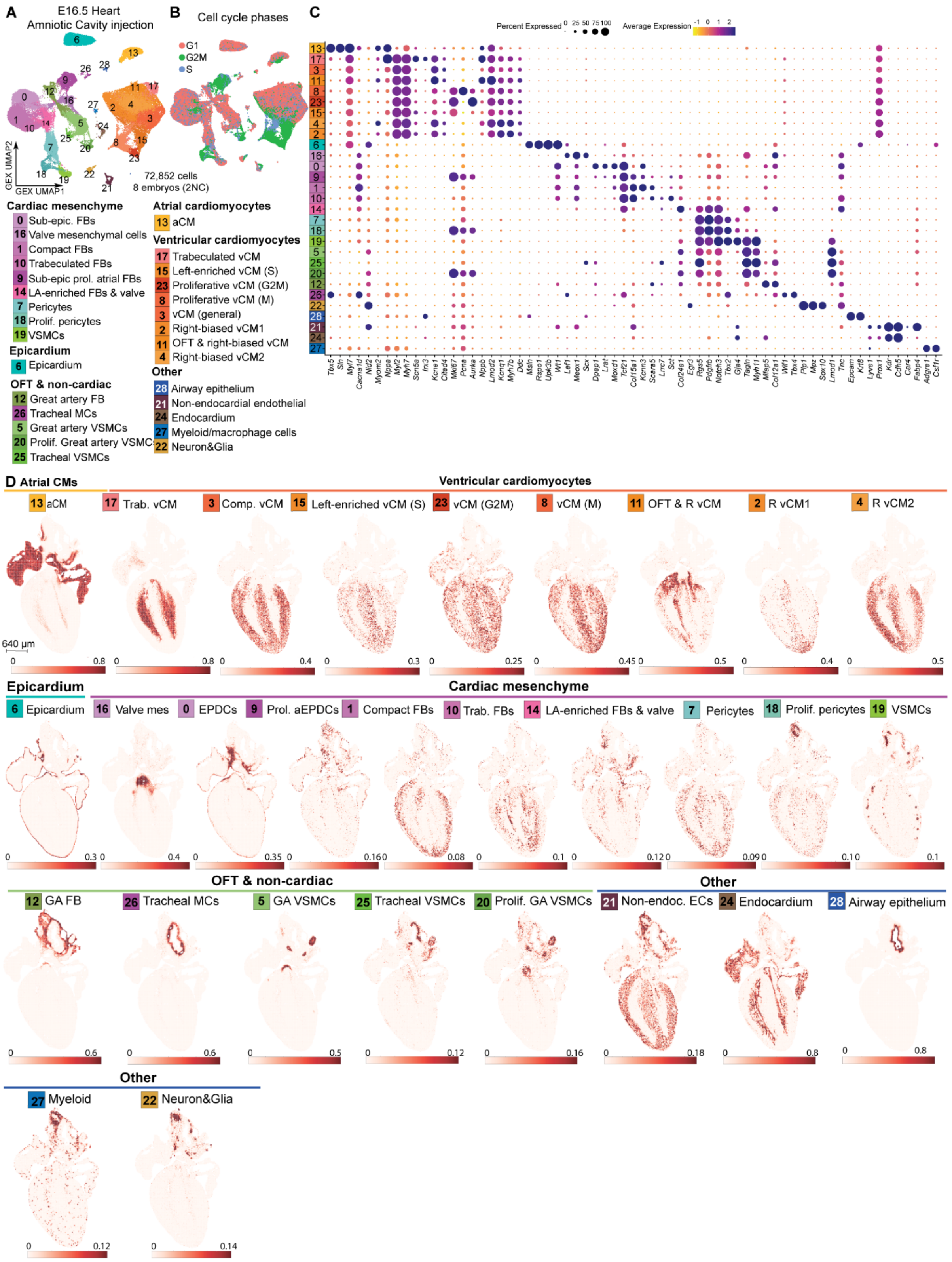


**Figure S17. Spatial deconvolution of E16.5 heart sample #1 using heart scRNA-seq reference following E7.5 amniotic cavity injection, related to Figure 7.**

**(A,B)** UMAP visualization of the integrated object used for deconvolution, grouped by cell types (**A**) and cell cycle phases (**B**).

**(C)** Expression of canonical markers used to annotate the populations in **(A)**.

**(D)** Deconvolution of cell types.

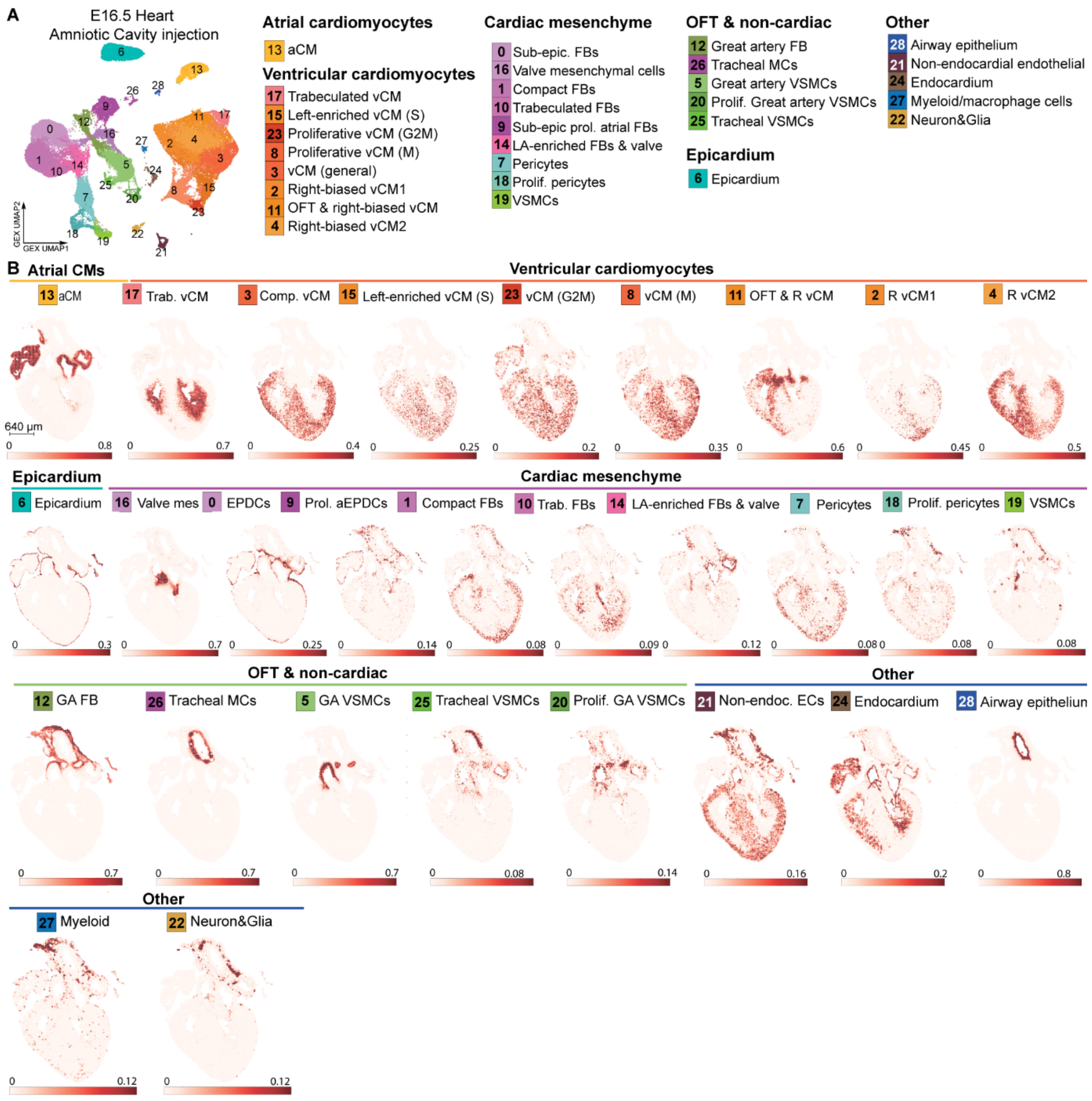


**Figure S18. Spatial deconvolution of E16.5 heart sample #2 using heart scRNA-seq reference following E7.5 amniotic cavity injection, related to Figure 7.**

(A) UMAP visualization of the integrated object used for deconvolution.

(B) Hematoxylin and eosin staining of section used for spatial transcriptomics, as an anatomical reference.

(C) Deconvolution of cell types.

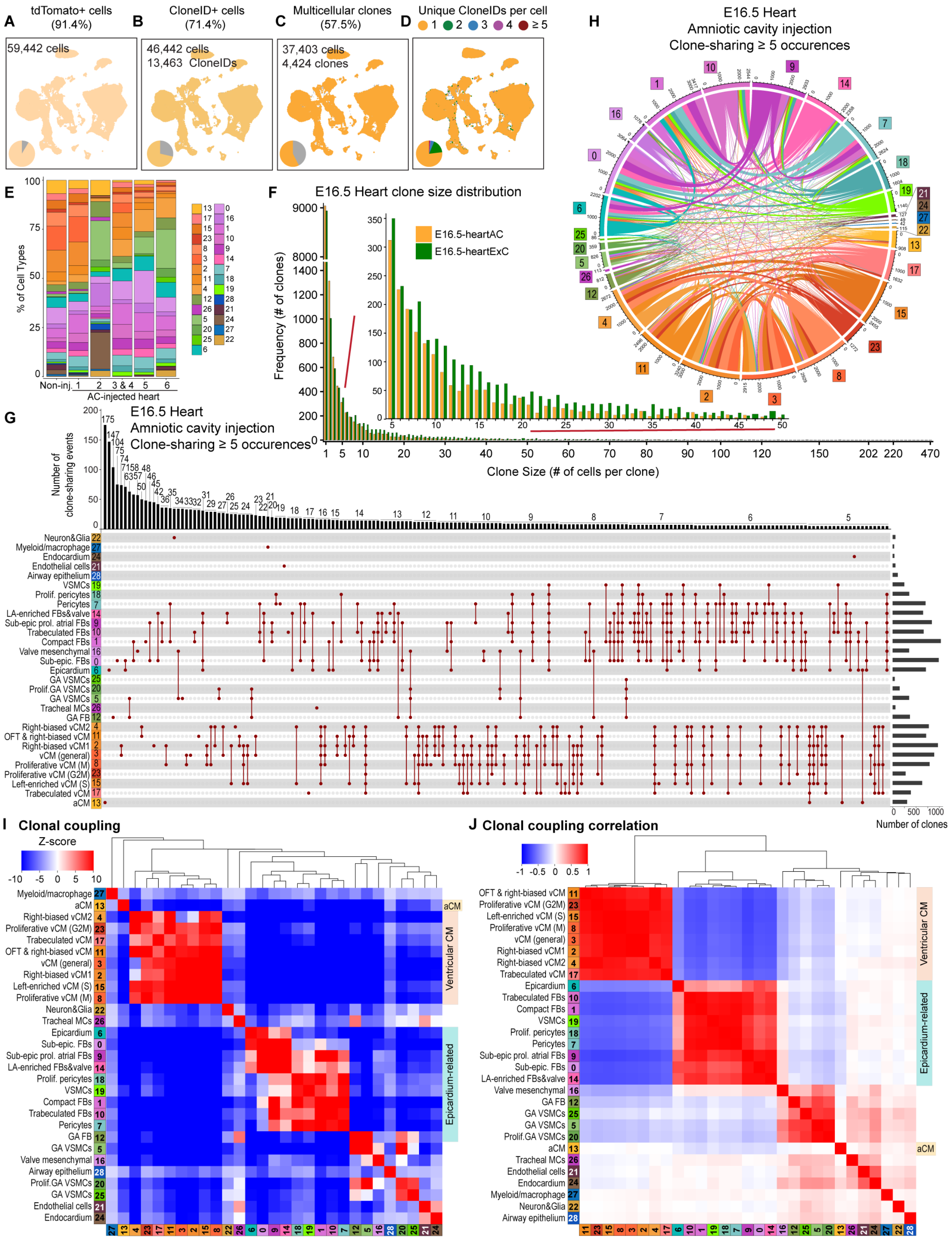


**Figure S19. Clonal characterization and analysis of E16.5 hearts following E7.5 amniotic cavity injections, related to Figure 7.**

**(A-D)** UMAPs and pie charts showing tdTomato+ cells (**A**) CloneID+ cells (**B**), multicellular CloneID+ clones (clone size ≥2 cells; **C**) and the number of CloneIDs per cell (**D**) among all cells isolated from E16.5 hearts after E7.5 amniotic cavity injection.

**(E)** Stacked barplot showing the proportion of different cell types in cells expressing tdTomato RNA.

**(F)** Bar plot showing the clone size distribution in integrated E16.5 hearts following E7.5 amniotic or exocoelomic cavity injection, for comparison. AC injections in yellow, ExC injections in green.

**(G)** UpSet plot showing clonal relationships across cell types (clone-sharing ≥5 occurrences) in the integrated E16.5 hearts after E7.5 amniotic and exocoelomic cavity injection. The top bars indicate the number of clone-sharing events among cell types or within a single cell type, and the right bars indicate the total clone numbers containing the indicated cell types.

**(H)** Circos plot showing pairwise co-occurrence of cell types across cardiac clones. Link width indicates the number of contributing cell type pairs; link color represents cell type.

**(I,J)** Heatmaps of lineage coupling Z-scores **(I)** and correlations **(J)** between pairs of all cell types in E16.5 hearts collected from AC and ExC injections, clustered by correlation distance and linkage.

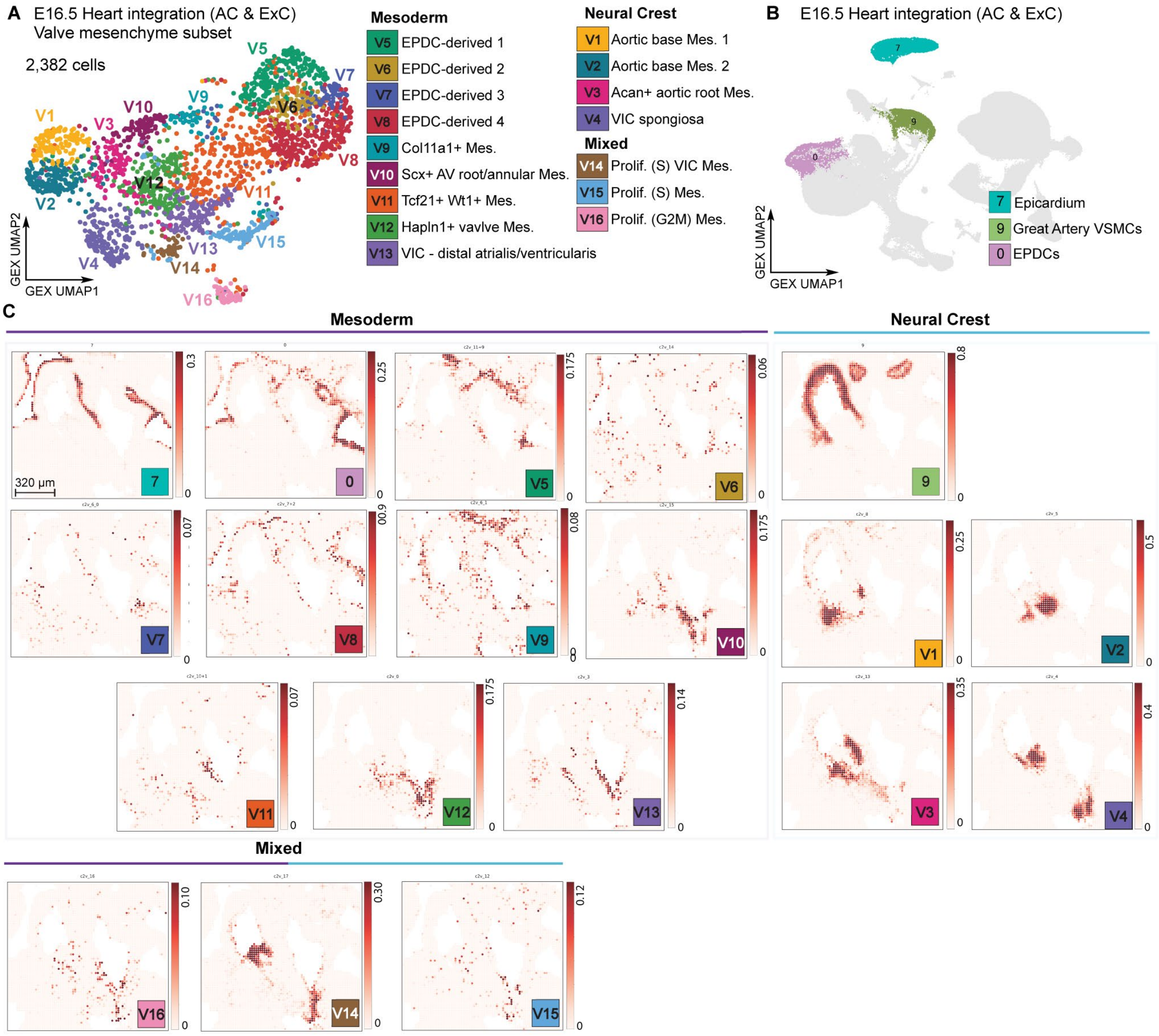


**Figure S20. Spatial deconvolution of integrated valve-mesenchymal subclusters in E16.5 heart sample #2, related to Figure 7.**

**(A)** UMAP representation of 2,382 valve mesenchymal cells subset from the integrated E16.5 heart dataset following E7.5 amniotic- or exocoelomic-cavity labelling, colored by subcluster. Subclusters are grouped according to mesodermal, neural crest or mixed clonal linkage.

**(B)** UMAP representation of the integrated heart dataset highlighting epicardium (cl.7), EPDCs (cl.0) and great-artery VSMCs (cl.9), which served as mesodermal and neural crest-related reference populations.

**(C)** Visium HD spatial deconvolution of E16.5 heart sample #2 at 16-μm resolution, showing mesoderm-linked epicardium, EPDCs and valve mesenchymal subclusters V5–V13; neural crest-linked great-artery VSMCs and subclusters V1–V4; and proliferative subclusters V14–V16 with mixed lineage associations.

AC, amniotic cavity; AV, atrioventricular; ExC, exocoelomic cavity; EPDC, epicardium-derived
cell; Mes., mesenchyme; VIC, valve interstitial cell; VSMC, vascular smooth muscle cell.

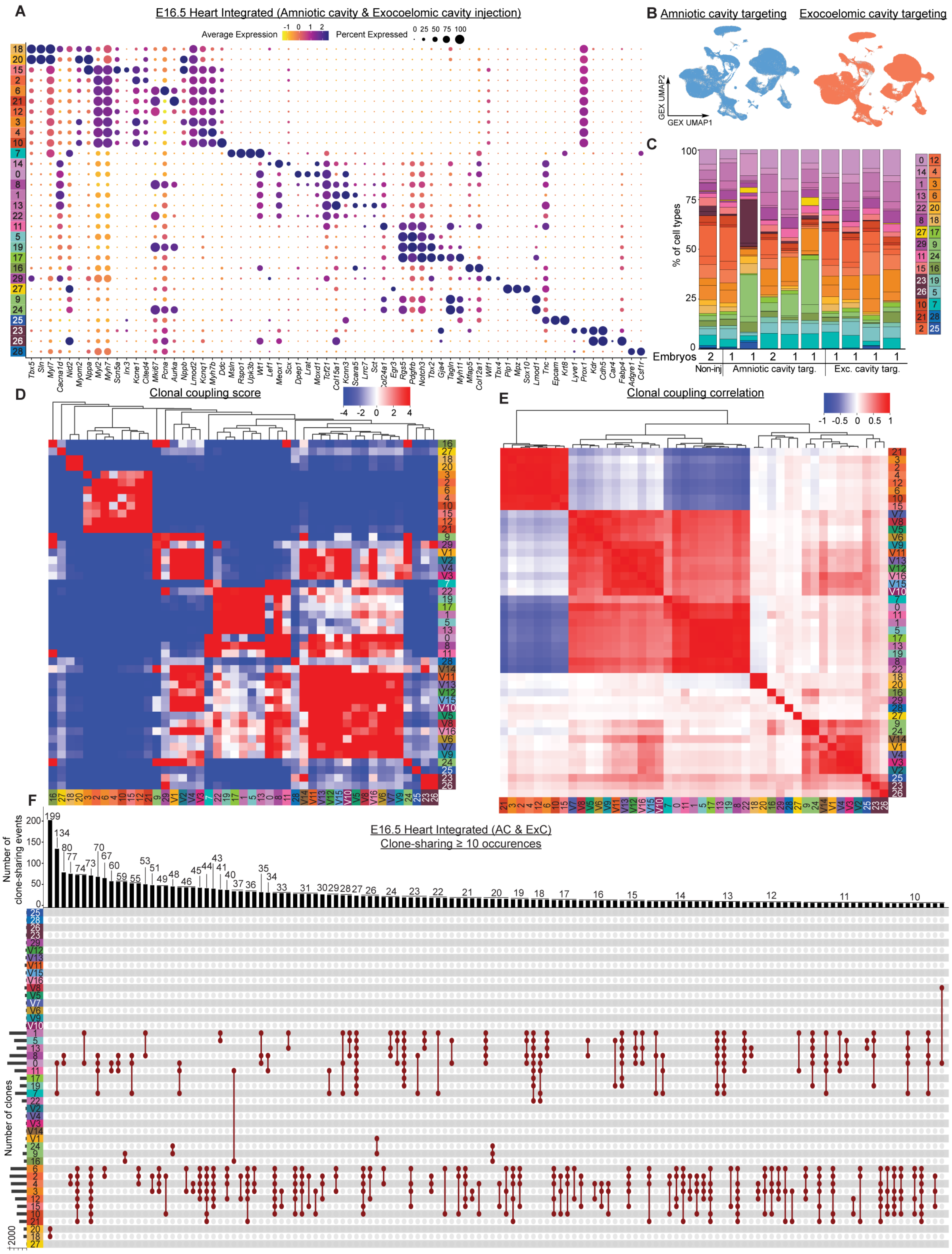


**Figure S21. Integrated cellular and clonal analysis of E16.5 hearts following E7.5 amniotic- and exocoelomic-cavity labelling, related to Figure 7.**

**(A)** Expression of canonical markers used to annotate the integrated exocoelomic and amniotic E16.5 heart populations in **Fig. 7A**.

**(B)** UMAP visualization of targeted clusters by each injection route.

**(C)** Comparison of cell-type representation between injection routes per embryo. Data comprises 12 embryos, including 6 subjected to amniotic-cavity injections, 4 subjected to exocoelomic-cavity injections and 2 non-injected controls.

**(D)** Clonal coupling analysis of integrated exocoelomic and amniotic E16.5 heart populations. Clonal coupling Z-scores quantify the enrichment of observed barcode sharing relative to randomized datasets preserving cell-population abundances, with a positive (red) score indicating enriched coupling and a negative (blue) score indicating under-represented coupling.

**(E)** Correlation of clonal coupling profiles across integrated heart cell populations.

**(F)** UpSet plot showing integrated E16.5 heart clones (amniotic and exocoelomic cavity injections), clonal relationships across cell types (clone-sharing ≥10 occurrences). The top bars indicate the number of clone-sharing events among cell types or within a single cell type, and the left bars indicate the total clone numbers containing the indicated cell types**.**

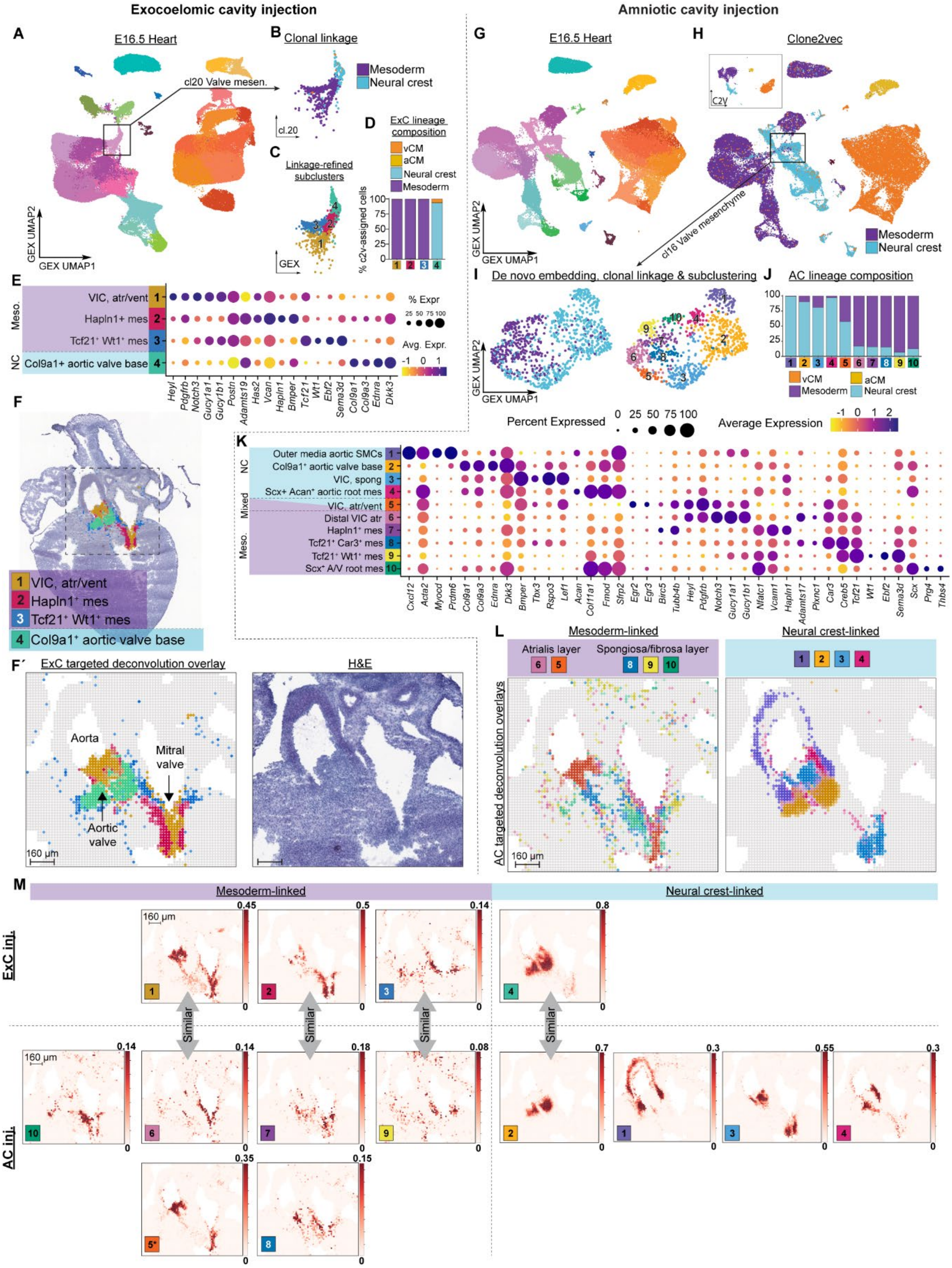

Exocoelomic cavity injection
Amniotic cavity injection
A
E16.5 Heart
B
Clonal linkage
Mesoderm
Neural crest
cl20 Valve mesen.
C
Linkage-refined subclusters
D
ExC lineage composition
vCM
aCM
Neural crest
Mesoderm
% c2v-assigned cells
GEX UMAP1
GEX UMAP2
E
VIC, atr/vent
Hapln1+ mes
Tcf21+ Wt1+ mes
Col9a1+ aortic valve base
% Expr
Avg. Expr.
F
VIC, atr/vent
Hapln1+ mes
Tcf21+ Wt1+ mes
Col9a1+ aortic valve base
F′
ExC targeted deconvolution overlay
H&E
Aorta
Mitral valve
Aortic valve
160 µm
G
E16.5 Heart
H
Clone2vec
cl16 Valve mesenchyme
I
De novo embedding, clonal linkage & subclustering
J
AC lineage composition
K
Percent Expressed
Average Expression
Outer media aortic SMCs
Col9a1+ aortic valve base
VIC, spong
Scx+ Acan+ aortic root mes
VIC, atr/vent
Distal VIC atr
Hapln1+ mes
Tcf21+ Car3+ mes
Tcf21+ Wt1+ mes
Scx+ A/V root mes
L
Mesoderm-linked
Neural crest-linked
Atrialis layer
Spongiosa/fibrosa layer
AC targeted deconvolution overlays
M
Mesoderm-linked
Neural crest-linked
ExC inj.
AC inj.
Similar

**Figure S22. Individual analyses of cell population subsets within cardiac valve mesenchyme clusters at E16.5, from embryos targeted with either exocoelomic cavity or amniotic cavity injections at E7.5, related to Figure 7.**

**(A)** UMAP visualization of E16.5 heart following E7.5 exocoelomic cavity injection**.**

**(B,C)** 1,561 valve mesenchymal cells (cl.20, boxed cluster in **(A)** were lineage-inferred using clone2vec (clone size ≥ 3) **(B)** and re-clustered **(C).**

**(D)** Clone2vec lineage composition comparison of clusters in **(C)**.

**(E)** Expression of canonical markers used to annotate the populations in **(C)**.

**(F)** Spatial localization of re-clustered valve mesenchymal cells **(C).**

**(G)** UMAP visualization of E16.5 heart following E7.5 amniotic cavity injection**.**

**(H)** UMAP visualization of clone2vec-inferred lineages (clone size ≥ 2), and clone2vec embedding of parental UMAP from **(G)** clones (boxed).

**(I)** 976 valve mesenchymal cells (cl.16, boxed cluster) were re-clustered and re-embedded, and had clone2vec lineages **(H)** overlaid**.**

**(J)** Clone2vec lineage composition comparison of clusters in **(I)**.

**(K)** Expression of canonical markers used to annotate the populations in **(I)**.

**(L)** Spatial localization of re-clustered valve mesenchymal cells **(I).**

**(M)** Comparative spatial localization of exocoelomic **(C)** and amniotic **(I)** clusters, the absence of a similarity arrow indicates a population unique to that injection route. *asterisk for population 5 indicates dual lineage coupling.
